# The genetic architecture of the human skeletal form

**DOI:** 10.1101/2023.01.03.521284

**Authors:** Eucharist Kun, Emily M. Javan, Olivia Smith, Faris Gulamali, Javier de la Fuente, Brianna I. Flynn, Kushal Vajrala, Zoe Trutner, Prakash Jayakumar, Elliot M. Tucker-Drob, Mashaal Sohail, Tarjinder Singh, Vagheesh M. Narasimhan

## Abstract

The human skeletal form underlies our ability to walk on two legs, but unlike standing height, the genetic basis of limb lengths and skeletal proportions is less well understood. Here we applied a deep learning model to 31,221 whole body dual-energy X-ray absorptiometry (DXA) images from the UK Biobank (UKB) to extract 23 different image-derived phenotypes (IDPs) that include all long bone lengths as well as hip and shoulder width, which we analyzed while controlling for height. All skeletal proportions are highly heritable (∼40-50%), and genome-wide association studies (GWAS) of these traits identified 179 independent loci, of which 102 loci were not associated with height. These loci are enriched in genes regulating skeletal development as well as associated with rare human skeletal diseases and abnormal mouse skeletal phenotypes. Genetic correlation and genomic structural equation modeling indicated that limb proportions exhibited strong genetic sharing but were genetically independent of width and torso proportions. Phenotypic and polygenic risk score analyses identified specific associations between osteoarthritis (OA) of the hip and knee, the leading causes of adult disability in the United States, and skeletal proportions of the corresponding regions. We also found genomic evidence of evolutionary change in arm-to-leg and hip-width proportions in humans consistent with striking anatomical changes in these skeletal proportions in the hominin fossil record. In contrast to cardiovascular, auto-immune, metabolic, and other categories of traits, loci associated with these skeletal proportions are significantly enriched in human accelerated regions (HARs), and regulatory elements of genes differentially expressed through development between humans and the great apes. Taken together, our work validates the use of deep learning models on DXA images to identify novel and specific genetic variants affecting the human skeletal form and ties a major evolutionary facet of human anatomical change to pathogenesis.

## Introduction

Humans are the only primates who are normally biped, due to our unique skeletal form which stabilizes the upright position. Bipedalism is enabled by unique anatomical properties of the human skeleton, including shorter arms relative to legs, a narrow body and pelvis, and orientation of the vertebral column (*1–3*). These broad changes to skeletal proportions likely began to occur around the separation of the human and chimp lineages and as a result, may have facilitated the use of tools and accelerated cognitive development (*4, 5*). Fossil evidence showing major morphological changes in the length of the limbs, torso, and body width suggest that these changes were gradual, with incremental development over the course of several million years (*6, 7*). Perhaps one of the first bipedal hominins, *A. afarensis*, dated to 3.2 million years before present, exhibited skeletal adaptations such as a shorter iliac blade and wider sacrum in the pelvis as well as a human-like bicondylar angle of the femur and talocrural joint, all of which are crucial for bipedal movement (*8, 9*). Another major change in skeletal proportions can be seen in *H. ergaster,* which exhibit modern human-like intermembral proportions with longer legs relative to the torso and arms than compared to great apes (*6, 10*). However, despite over a hundred years of effort in paleoanthropology documenting morphological change in human evolution, evidence of genomic change shaping the human skeletal form has been elusive. Separately, human body proportions have been the subject of the study of art for several millennia. The Italian polymath Leonardo Da Vinci famously drew one of the most enduring images of the Rennaiscance, the Vitruvian Man in 1490, which depicted a human male with arms and legs inscribed in a circle and a square reflecting the artist’s idealization of the human form.

In developmental biology, the mechanisms and processes underlying animal limb development, morphology, and broad body plan have been studied extensively. Early work using forward genetic screens in Drosophila identified homeobox genes as key regulators of large-scale anatomical development in invertebrates (*11*). Subsequent experiments in a large number of vertebrates including fish, chickens, and mice, identified additional gene families crucial in limb formation and skeletal form regulation (*12, 13*). In addition to this, comparative genomic and evolutionary developmental biology approaches have examined genetic variation or gene expression between related species with differing phenotypes to pinpoint genes associated with the skeletal form. This line of work has produced several insights into the genetic basis of skeletal structure from the underpinnings of convergent limb loss in snakes and limbless lizards (*14, 15*) to increased limb lengths in jerboas when compared to mice (*16*). Analyses studying differences in gene expression and regulation in skeletal tissues between humans and other primates have recently identified a gene involved in tail loss in great apes (*17*), as well as shown that open chromatin regions specific to the human knee and hip joint overlap with regions that are evolutionarily accelerated in humans (*18*). However, these approaches do not provide an unbiased and comprehensive map of genetic variants regulating skeletal proportions and overall body plan, especially in humans. Furthermore, many of these approaches largely focus on examining the impact of loss-of-function mutations, which often have widespread effects on the entire skeleton. The subset of genes responsible for differential and specific growth of individual bones remains unknown.

Genome-wide association studies of human skeletal traits are a direct and complementary approach to identifying biological mechanisms and processes that underlie limb proportions, development, and morphology. Twin studies suggest that the heritability of skeletal proportions range between 0.40 - 0.80 (*19*), about as heritable as standing height (*20*), a skeletal trait that has served as an exemplary quantitative trait in human genetics. Meta-analysis of over 5 million individuals has identified 12,111 independent single nucleotide polymorphisms (SNPs) associated with standing height (*21*). However, height is amongst the most straightforward and accurate of quantitative traits to measure. Other skeletal elements, such as limb, torso, and shoulder lengths, are not typically or comprehensively measured in large sample sizes (*22, 23*). As a result, the genetic basis of such proportions and lengths remains understudied. Furthermore, anthropometric traits, like hip and waist circumference, are measured externally and therefore are intrinsically tied to body fat percentage and distribution, which fails to isolate genetic effects specific to the skeletal frame (*24, 25*).

Applying deep learning methods to non-invasive medical imaging is a powerful way to extract skeletal measures in an accurate and scalable manner. Furthermore, the collection of genetic, phenotypic, and imaging data by national biobanks provides an opportunity to run GWAS for IDPs with sufficiently large sample sizes. Several genetic studies have successfully applied computer vision to generate IDPs of the retina, distribution of body fat, heart structure, and liver fat percentage, and linked significant loci to various disorders (*26–29*).

In the context of musculoskeletal disease, epidemiological data suggests that disorders such as osteoarthritis – the leading cause of adult disability in the United States (*30, 31*) are thought to be influenced by a variety of risk factors ranging from obesity, mechanical stresses, genetic factors, and even the geometric structure of certain bones (*32*). While some small sample studies have examined the relationship of certain skeletal element lengths such as leg length discrepancy and osteoarthritis (*33*), how the skeletal frame may exacerbate an individual’s development of osteoarthritic disease has not been fully studied (*32, 34*).

In this study, we adapt, test, and apply methods in computer vision to derive comprehensive human skeletal measurements from full body DXA images of tens of thousands of adult participants. We then perform genome-wide scans on 23 generated phenotypes to identify loci associated with variation in the skeletal form. Using summary statistics from these IDPs, we identify biological processes linked with human skeletal proportions and study the phenotype and genetic correlation between these measures and a range of external phenotypes, with an emphasis on musculoskeletal disorders. Finally, we investigate the impact of natural selection on these traits to understand how skeletal morphology is linked to human evolution and bipedalism.

## Results

### A deep learning approach for quality control and quantification of biobank scale imaging data

To study the genetic basis of human skeletal proportions, we jointly analyzed DXA and genetic data from 42,284 individuals in the UKB. Individuals from this dataset are aged between 40 to 80 and reflect adult skeletal morphology. We report baseline information about our analyzed cohort in **Methods**: *UKB participants and dataset* and in **Table S1**. We acquired 328,854 DXA scan images across eight imaging modalities comprising full-body transparent images, full-body opaque images, anteroposterior (AP) views of the left and right knees, AP views of the hips, and AP and lateral views of the spine. For quality control, we first developed a deep learning-based multi-class predictor to select full body transparent images from the pool of eight total imaging modalities. We developed a second deep learning classifier to remove cropping artifacts. Finally, we excluded images with atypical aspect ratios and padded them to uniform lengths (**Methods**: *Classification of DXA Images by body part, Removal of poorly cropped X-rays, Image standardization*, **Fig. 1A**). After our quality control process, we were left with 39,469 images for analysis.

**Fig. 1.**
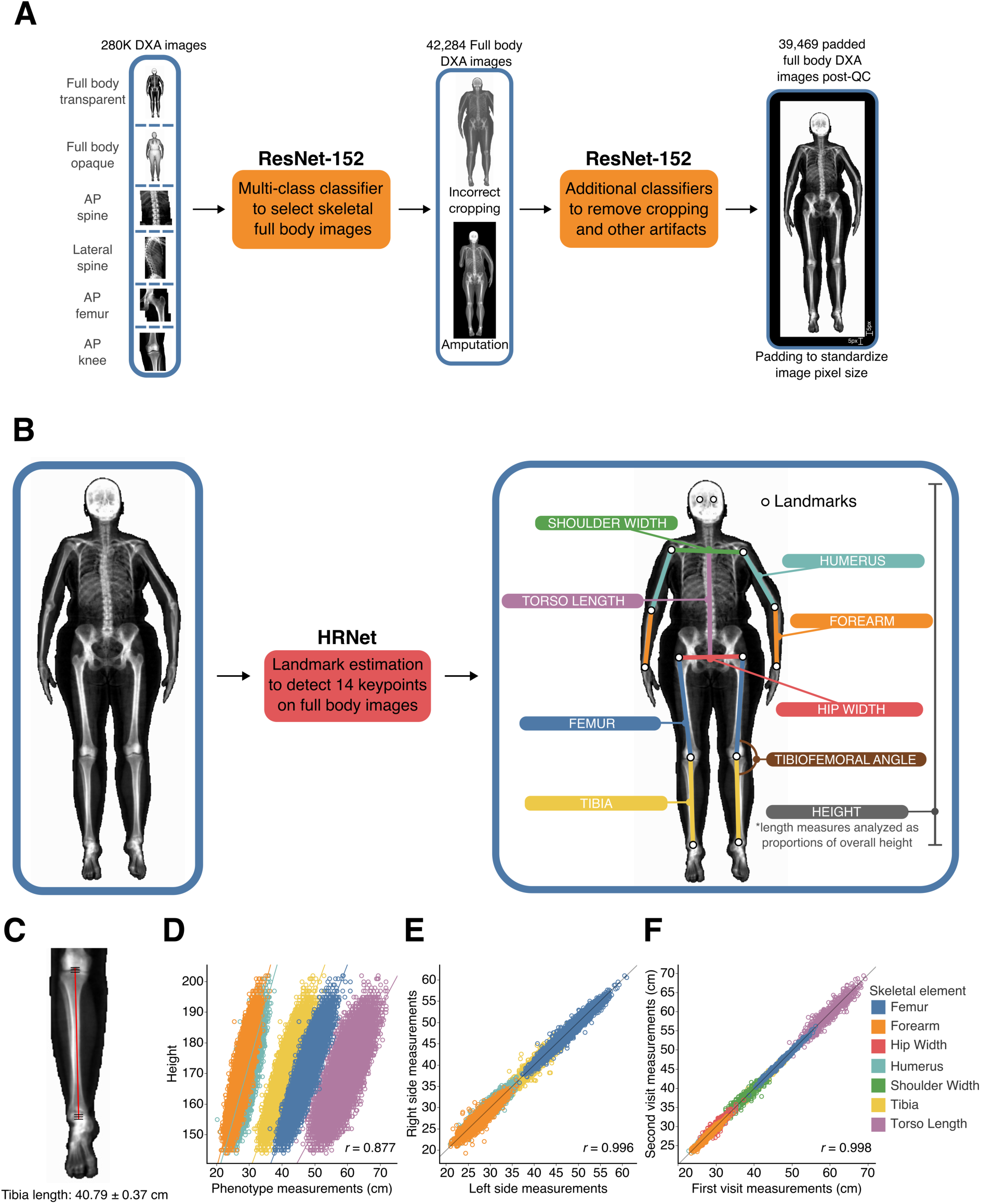
Deep learning-based image quantification. (**A**) *Quality control*. Deep learning-based classifiers to select full body images from a pool of DXA images of different body parts, as well as to remove images with artifacts, resolution, or cropping issues. Full body images were then padded to standardize image pixel size before phenotyping (current image shows padding of 5 pixels on each side). (**B**) *Image quantification*. Deep learning-based image landmark estimation using the HRNet architecture. 297 training images annotated with specific landmarks were used to train the model to perform automatic annotation of landmarks on the rest of images in the dataset from which measurements of skeletal length and other measurements were calculated. (**C**) Average HRNet measurement error when compared to human-derived measurements of the tibia across 100 validation images. (**D**) Correlation of length measurements and height. (**E**) Correlation between left and right-side measurements of the femur, humerus, forearm, and tibia. (**F**) Correlation of lengths measured from the first imaging visit and second imaging visit for the same individual.

**Fig. 2.**
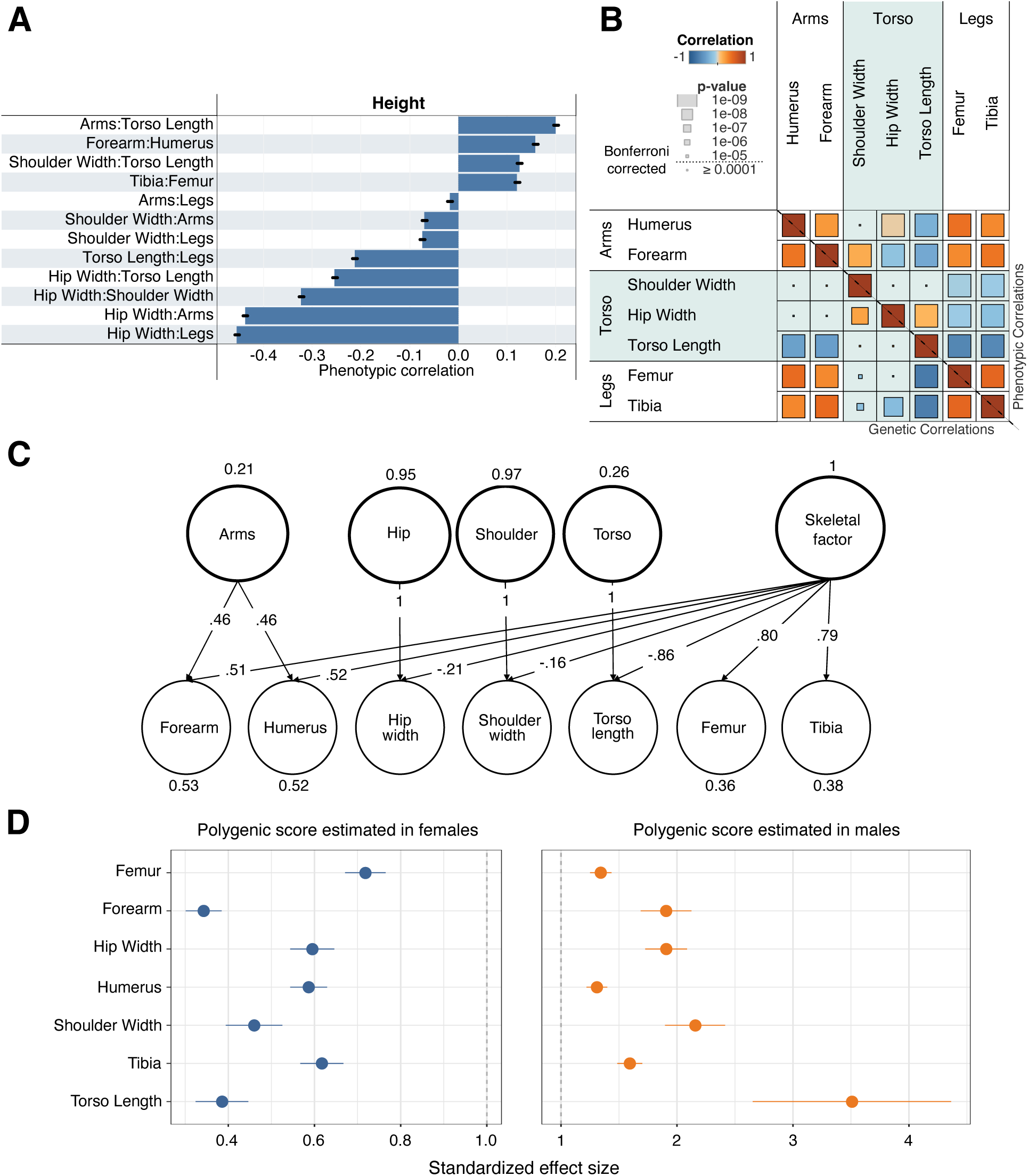
Genetic architecture. (**A**) Correlation of skeletal proportions and overall height. Bars show ±2 SE. (**B**) Genotype (lower-left triangle) and phenotype (upper-right triangle) correlation of skeletal proportions. Overall correlation is shown in color and the p-value of the correlation is visualized by size. A Bonferroni-corrected threshold is also shown. (**C**) Solution for a genomic SEM model for the genetic covariance structure shown in **B** shows one common factor loading for arms, an additional factor for legs, and finally independent factors for each of the torso-related traits (hip width, shoulder width, and torso length). (**D**) Sex-specific analysis showing the ratio of the standardized effect size of the polygenic score on each trait (±2 SE) in males to the effect in females in a hold-out dataset.

After image QC, we manually labeled 14 landmarks at pixel-level resolution on 297 images for use as training data. These labels were independently validated by an orthopedic team. The 14 landmarks include major joints comprising the wrist, elbow, shoulder, hip, knee, ankle, and positions of the eyes. Each landmark represents major joints in the body, and the segments connecting them reflect natural measurements for long bone lengths or body width measures. We assessed the replicability of manual annotation by inserting 20 duplicated images from the 297 training images without the knowledge of the annotator and found that repeat measurements resulted in less than 2 pixels of difference at any landmark (**Methods:** *Manual annotation of human joint positions*, **Fig. 1B**).

We adapted and applied a new computer vision architecture, High-Resolution Net (HRNet), for landmark estimation, or the prediction of the location of human joints (*35*). Our choice was guided by four main reasons. First, HRNet maintains high-resolution representations throughout the model (**Methods:** *A deep learning model to identify landmarks on DXA scans*), and we wanted to utilize the high-resolution medical images produced by the DXA scanner to obtain precise measurement information of bone lengths. Second, the architecture had already been trained on two large imaging datasets, first on imageNet (*36*), a general natural image dataset, and then subsequently on Common Objects in Context (COCO) (*37*), a dataset of over 200,000 images of humans in natural settings with joint landmarks classified. These two previous layers of training enabled us to perform transfer learning to fine tune the architecture on our training data and reduce the total amount of manual annotation to just 297 images. Third, HRNet has among the best performance for a similar task of labeling human joints on two large-scale benchmarking data sets of human subjects (*38, 39*). Finally, we directly compared the performance of the HRnet architecture with a more traditional architecture on our dataset (ResNet-34) (*40*) and obtained significantly better results across different training parameter choices (**Methods:** *A deep learning model to identify landmarks on DXA scans*, **Table S2**). Upon training, the model achieved greater than 95% average precision on hold-out validation data across all body parts (**Table S2**).

### Validation of human skeletal length estimates

After training and validating the deep learning model on the 297 manually annotated images, we applied this model to predict the 14 landmarks on the rest of the 39,172 full body DXA images. We then calculated pixel distances between pairs of landmarks that corresponded to 7 bone and body lengths segments (**Fig. 1B**, **Methods:** *Obtaining skeletal element length measures, Obtaining a set of body proportion traits from raw length measures,* **Table S3**). We also computed an angle measure between the tibia and the femur (tibiofemoral angle or TFA) (**Fig. 1B**). To standardize images with different aspect ratios, we rescaled pixels into centimeters for each image resolution by regressing height in pixels against standing height in centimeters as measured by the UKB assessment (**Methods:** *Adjusting for scaling differences across imaging sizes and modes*). We then removed individuals with any skeletal measurements that were more than 4 standard deviations from the mean (**Methods:** *Removal of image outliers*).

After outlier removal, we validated the accuracy of our measurements on the remaining samples in four ways. First, the error rate for segment length from the model compared to manual annotation was at maximum 3 pixels or 0.7 cm, which is similar to the variation from manual annotation of the 20 duplicate images. Reliability (100%-variance in measurement/variance of a segment length) was greater than 95% across all length measures (**Fig. 1C**, **Methods:** *Validation metrics comparing automated annotation to manual annotation*, **Table S4-Table S6**). Second, the correlation between long bone lengths and height as measured in the UKB was around ∼0.88, which falls within expectation observed in the literature (*22*) (**Fig. 1D**). Third, the correlation between left and right limb lengths was above 0.99 (**Fig. 1E**). Fourth, a subset of 667 individuals had undergone repeat imaging an average of two years apart, with different image aspect ratios, DXA machines, software models, and technicians carrying out the imaging (**Fig. 1F**). The correlation in these technical replicates across skeletal elements was also above 0.99. Taken together, these results suggest that the IDPs from our deep learning model are highly accurate and highly replicable.

### Characteristics and correlations of human skeletal proportions with sex, age, and height

From the 7 bone and body segment lengths, to examine these IDPs as proportions instead of lengths (or to control for variation in overall height which is highly correlated with each of these lengths) we took simple ratios of each IDP with overall height (**Fig. 1B**, **Methods:** *Obtaining a set of body proportion traits from raw length measures*). As expected, this greatly reduced the overall correlation of our traits with height (**Table S7**). We also carried out this normalization analysis in alternate ways, including using height as a covariate in association tests as well as regressing each IDP with height and obtaining residuals. All three approaches were highly correlated, and we used the simple approach of taking proportions for most analyses (**Methods:** *Adjusting for height correlation in GWAS using ratios*). In addition to obtaining ratios of each segment length with overall height, we also computed ratios of segments with each other obtaining a total of 21 different ratio IDPs along with the angle measure, TFA (**Table S3**). These ratios are referred to in the text as Segment:Segment (Hip Width:Height, Torso Length:Legs, etc). In **Fig. 3B**, we show our mean proportions of each skeletal element across all of our individuals of white British ancestry (*41*).

**Fig. 3.**
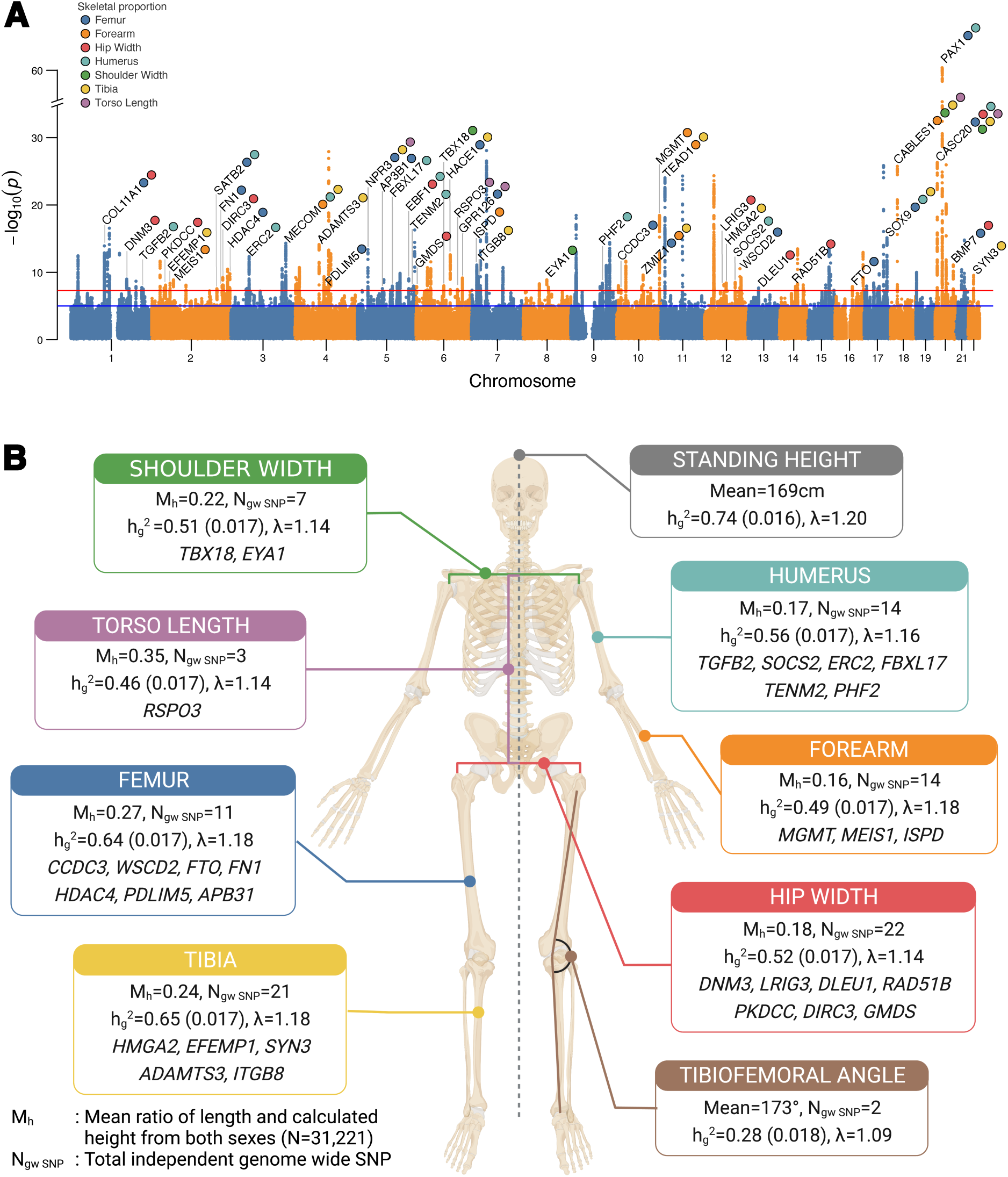
Genome-wide association results. (**A**) Manhattan plot of GWAS performed across 7 skeletal proportions and TFA, with the lowest p-value for any trait at each SNP annotated. Loci over the genome-wide significance threshold that are in close proximity to only a single gene are annotated. Colors show the traits for which each SNP is genome-wide significant. (**B**) Mean values of proportion and angle traits across individuals, total number of genome-wide significant loci per trait, heritability, lambda (from LDSC), and associated genes of loci independent of height that are specific to each skeletal trait (again annotating only loci that map to a region with a protein-coding gene within 20 kb of each clumped region).

We then examined differences in skeletal proportions across sex and age. In line with well-known observations, Hip Width:Height (t-test p < 10^-15^) and Torso Length:Height (t-test p < 10^-15^) were significantly larger in women than in men (*42*), but we also observed that Humerus:Height was also significantly larger in women than in men (t-test p = 1.45 × 10^-5^) (**Methods:** *Correlations of skeletal proportions with age and sex*, **Table S8**). In addition, we found that all body proportions vary slightly but significantly as a function of age (**Methods:** *Correlations of skeletal proportions with age and sex*, **Table S9**). We also examined how body proportions vary as a function of overall height and found that Torso Length:Legs decreases with height (Pearson correlation r = -0.21), suggesting that increases in height are driven more by increasing leg length rather than torso length (**Fig. 2A**). Arms:Legs also decreases with height (Pearson correlation r = -0.02) meaning that leg length also outpaces arm length as height increases. Within each limb, for both arms and legs, lower to upper limbs ratios (Tibia:Femur, Forearm:Humerus) increase with overall limb length. These increases also correspond with correlations with height, with Tibia:Femur increasing when height increases (Pearson correlation r = 0.12).

### GWAS of human skeletal proportions

We performed GWAS using imputed genotype data in the UKB to identify variants associated with each skeletal measure. We applied standard variant and sample QC and focused our analyses on 31,221 individuals of white British ancestry as determined by the UKB genetic assessment and 7.4 million common bi-allelic SNPs with minor allele frequency > 1% (*41*) (**Methods:** *Genetic data quality control, Heritability analysis and GWAS*, **Table S1 and Table S10**). We used Bolt-LMM (*43*) to regress variants on each skeletal measure using a linear mixed-model association framework (**Methods:** *Heritability analysis and GWAS*). After generating summary statistics for each skeletal measure, we estimated SNP heritability using LD Score regression (LDSC) (*44*) and GCTA-REML (*45*). All traits were highly heritable, with SNP heritability 30% - 60% for LDSC and 40% - 70% for GCTA-REML (**Table S11 and Table S12**). We detected inflation in test statistics in our QQ plots (mean lambda = 1.20); however, minimal deviation of univariate LDSC intercepts from 1.0 suggested that this inflation was consistent with polygenicity rather than confounding (**Methods:** *Heritability analysis and GWAS*, **Fig. 3B**).

In the seven skeletal proportions as a ratio of height (Forearm:Height, Humerus:Height, Tibia:Height, Femur:Height, Hip Width:Height, Shoulder Width:Height, Torso Length:Height) and TFA, we identified 223 loci at p *<* 5 × 10^-8^ and 150 loci at p < 6.25 x 10^-9^ (Bonferroni correction for eight traits). Of these loci, 179 are independently significant across all eight phenotypes (116 after Bonferroni correction for eight traits) (**Methods:** *Heritability analysis and GWAS*, **Supplementary Data - GWAS Summary statistics**). Of the 179 independent loci, 77 are also associated with standing height, and 102 loci are only significant in skeletal proportions or TFA (**Methods:** *Clumping, independence analysis and removing previous height associated loci*). As sensitivity analysis we also examined the genetic effect of skeletal lengths before and after height adjustment and found that 95% of genome-wide significant loci had the same direction of effect when carrying out GWAS in these alternate ways (**Methods:** *Sensitivity analysis for height adjustment*).

### Genetic correlations and factor analysis of skeletal proportions

We calculated the genetic correlation between each pair of traits to investigate the degree of genetic sharing between each skeletal measure. Estimates from LDSC and GCTA-REML were virtually identical (**Fig. S10**); here we report estimates from GCTA-REML. Limb proportions had positive genetic correlations with each other (r_g_ = 0.34-0.55). Upper arms and legs (Humerus:Height-Femur:Height r_g_ = 0.55, p = 1.59 × 10^-66^) and lower arms and legs (Forearm:Height-Tibia:Height r_g_ = 0.51, p = 6.01 × 10^-50^) were significantly more correlated than upper arms and lower legs (Humerus:Height-Tibia:Height r_g_ = 0.38, p = 5.18 × 10^-23^) or lower arms and upper legs (Forearm:Height-Femur:Height r_g_ = 0.34, p = 1.49 × 10^-18^). Body width proportions, Hip Width:Height and Shoulder Width:Height, were largely uncorrelated with limb length proportions. No correlations involving any pairwise combination of arm and width traits were significant (minimum p-value across all such correlations was ≥ 0.0022, above our Bonferonni threshold). Correlations between leg and width traits were marginally significant in three out of four comparisons with the maximal correlation (Hip Width:Height-Tibia:Height) being 0.23 (**Fig. 2B, Table S13**). In addition, we also computed phenotypic correlations between our traits which were highly concordant with genetic correlations (r = 0.98).

We used Genomic Structural Equation Modeling (Genomic SEM) to identify latent factors that represent shared variance components between skeletal proportions (*46*) (**Methods:** *Multivariate genetic architecture of skeletal endophenotypes*). We performed exploratory factor analysis to identify the likely number of factors and built confirmatory models using odd-numbered chromosomes for model building and even-numbered chromosomes for validation, which we compared using a range of model fit indices. Our preferred model of the genetic covariance structure of the seven skeletal proportions indicates that all limb traits (both arms (Humerus:Height and Forearm:Height) and legs (Femur:Height and Tibia:Height) load positively on a general skeletal factor (on which Torso Length:Height loads negatively) and that the arm traits additionally load on a second general factor, whereas torso length and body width traits (Hip Width:Height and Shoulder Width:Height) only load appreciably on trait-specific factors (**Fig. 2C**). This analysis reinforces our observations from the univariate genetic correlation analysis, in which arm and leg proportions exhibited strong genetic sharing but were largely independent of torso and body width proportions.

### Sex-specific heritabilities and genetic effects of skeletal proportions

Anthropometric and skeletal traits, such as hip width, are common examples of sexual dimorphism. We found that for most traits, the genetic correlation of skeletal proportions between males and females was not statistically different from 1 except for TFA (r_g_ = 0.89) (**Methods:** *Sex-specific analysis*, **Fig. S16**). For five out of the seven skeletal proportions, the sex-specific SNP heritabilities were both greater than the heritability estimated jointly with both sexes (**Fig. S17**).

To test for pervasive differences in the magnitude of genetic effects, we performed sex-specific GWAS of all the skeletal traits and evaluated these polygenic scores in both sexes in a hold-out data set (**Methods:** *Sex-specific analysis*). This method had recently been applied to examine sex specific effects in biobank traits (*47*). Across all skeletal proportions that we tested, polygenic scores had a significantly larger standardized effect size (standardized in males and females separately) in males compared with females (t-test p < 1 × 10^-3^ for all comparisons) (**Fig. 2D**). These results are in-line with previous work suggesting that skeletal proportions, like other anthropometric traits, have clear differences in the magnitude of sex-specific effects when compared to other quantitative traits in the UKB (*47*).

### Biological insights from skeletal associations

We performed gene set enrichment analyses in 10,678 gene sets using FUMA to identify biological processes and pathways enriched in each skeletal trait (*48*) (**Methods:** *Functional mapping and gene enrichment analysis*). After FDR correction (FDR < 0.05), we found 195 gene sets to be significantly enriched across our 7 skeletal traits. Several gene sets related to development were common across the majority of traits such as skeletal system development, connective tissue development, chondrocyte differentiation, and cartilage development (**Table S14**).

Furthermore, common alleles associated with skeletal proportions were significantly enriched in 701 autosomal genes linked to “Skeletal Growth Abnormality” in the Online Mendelian Inheritance in Man (OMIM) (*49*) database (p < 5.0 × 10^-2^) except for torso length (p = 0.22) **(Methods:** *OMIM gene set enrichment analysis*, **Table S15 and Table S16**). Combined, these results indicate that common variants associated with skeletal proportions pinpointed genes in which rare coding variants contribute to Mendelian musculoskeletal disorders.

Out of the 223 total loci identified across GWAS (**Table S17**), 45 loci overlapped a single protein-coding gene within 20 kb of each clumped region. Notably, of these 45 genes, 32 (or 71%) resulted in abnormal skeletal phenotypes when disrupted in mice using the Human-Mouse Disease Connection database (*50*). Four of these genes (*COL11A1*, *SOX9*, *FN1*, *AGDRD6*) were associated with rare skeletal diseases in humans, annotated in OMIM (**Table S18**). In some cases, a gene linked with a specific skeletal proportion in our GWAS resulted in a defect in the same skeletal trait in mouse models. We found a common variant (rs6546231) near *MEIS1*, a homeodomain transcription factor, is associated with increased Forearm:Height. Mouse models of *MEIS*-/-mice are specifically associated with abnormal forelimb development (*51*). Similarly, a common variant (rs1891308) near *ADGRG6*, a G protein-coupled receptor, is associated with increased torso length. Mice with conditional knockouts in *ADGRG6* have spine abnormalities and spine alignments directly correlated with reduced torso length (*52*). Thus, GWAS of skeletal proportions pinpoints genes previously associated with skeletal developmental biology and Mendelian skeletal phenotypes and identifies candidates for future functional and knockout studies.

Next, we conducted a transcriptome-wide association study (TWAS), linking predicted gene expression in skeletal muscle (based on the Genotype-Tissue Expression project (GTEx v.7) (*53*) with our skeletal proportion GWAS (**Methods:** Transcriptome-wide associations (TWAS)). In total we identified 30 genes that were significantly associated with any one of our skeletal traits at a Bonferroni-corrected significance threshold across the total number of gene and trait combinations (**Table S19**). Among the strongest TWAS associations were *PAX1* (TWAS z-score = 12.6, p = 1.31 × 10^-36^), a transcription factor critical in fetal development and associated with development of the vertebral column, and *FGFR3* (TWAS z-score = 6.5, p = 8.52 × 10^-11^), a fibroblast growth factor receptor that plays a role in bone development and maintenance.

### Genetic and phenotypic association of skeletal phenotypes with musculoskeletal disease

To investigate the clinical relevance of human skeletal proportions, we examined their genetic and phenotypic associations with musculoskeletal disease and with joint and back pain. We used logistic regression to examine phenotypic associations between skeletal morphology and these musculoskeletal disorders (**Fig. 4A**) while controlling for age, sex, bone mineral density, BMI, and other major risk factors for OA (*54*) (**Methods:** *Phenotypic association of skeletal phenotypes with musculoskeletal disease*). We found one standard deviation in Hip Width:Height was associated with increased odds of hip OA (p = 3.16 × 10^-5^, OR = 1.34). Similarly, Femur:Height, Tibia:Height, and the TFA, skeletal measures of the knee joint, were associated with increased risk of knee OA (p = 2.24 × 10^-15^, OR = 1.34; p = 6.09 × 10^-5^, OR = 1.16; p = 1.64 × 10^-35^, OR = 1.49). Femur:Height and the TFA were also significantly associated with internal derangement of the knee (p = 4.03 × 10^-6^, OR = 1.19; p = 1.43 × 10^-17^, OR = 1.34). Pain phenotypes for hip and knee joints, were also associated with the specific skeletal proportions that make up each joint (hip pain with Hip Width:Height: p = 8.53 × 10^-5^, OR = 1.12; knee pain with Femur:Height, Tibia:Height, and TFA: p = 8.13 × 10^-6^, OR = 1.09; p = 2.89 × 10^-5^, OR = 1.09; p = 1.66 × 10^-46^, OR = 1.31) (**Fig. 4A**) (**Table S20**).

**Fig. 4.**
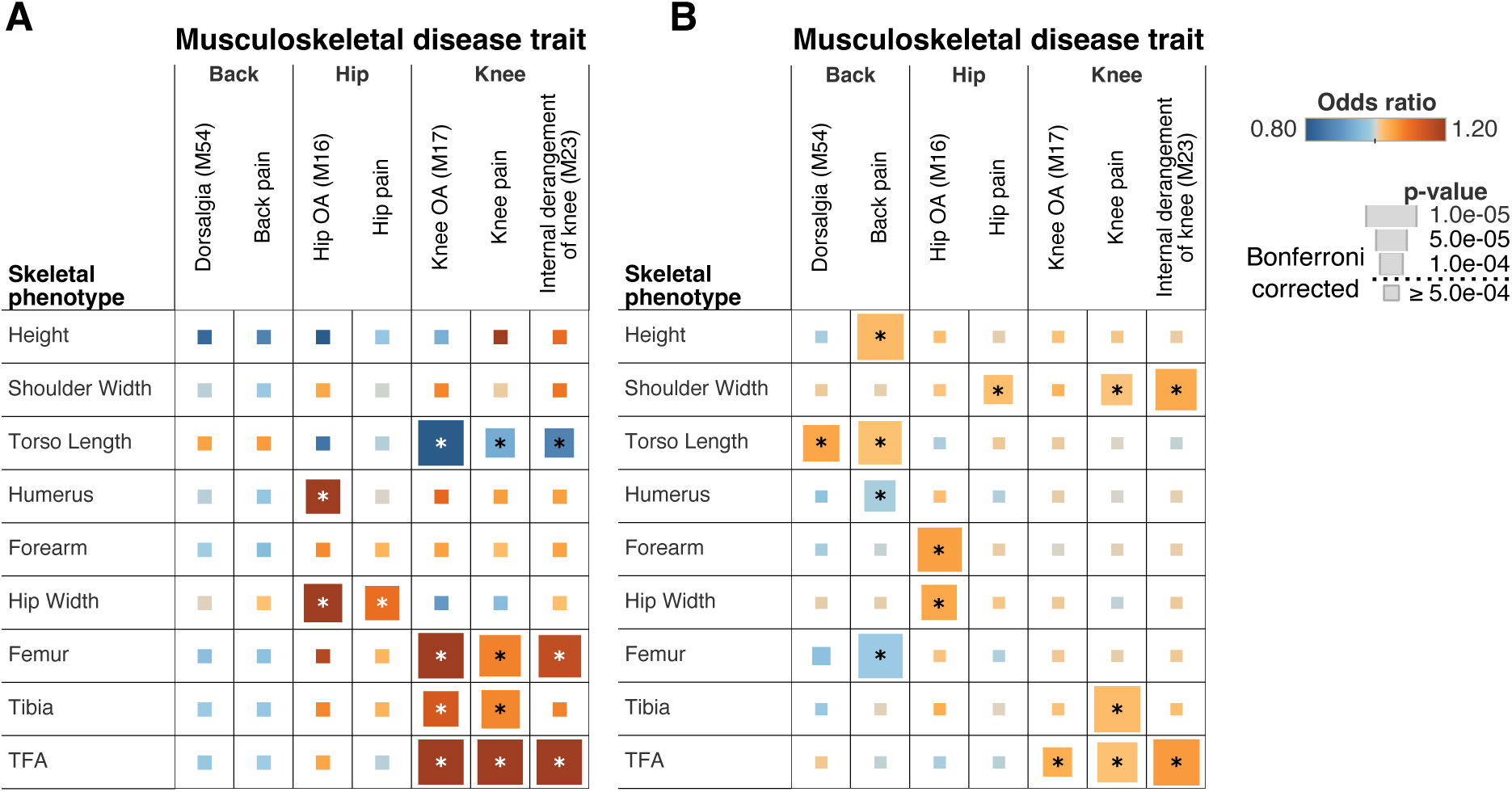
Association between skeletal traits and musculoskeletal disease. For both panels associations that are significant after Bonferroni correction are annotated with a *. Odds ratios for the phenotypic associations and PRS are shown in colors, and the p-values are represented by size. (**A**) Phenotypic associations from logistic regression analysis of musculoskeletal disease traits on skeletal phenotypes. (**B**) Polygenic risk score associations between musculoskeletal disease traits and skeletal phenotypes.

Next, we analyzed 361,140 UKB participants who had not undergone DXA imaging and were of white British ancestry for predictive risk based on polygenic scores derived from our GWAS on skeletal proportions on the imaged set of individuals (**Fig. 4B**). We generated polygenic scores via Bayesian regression and continuous shrinkage priors (*55*) using the significantly associated SNPs and ran a phenome-wide association study of the generated risk scores and traits, adjusting for the first 20 principal components of ancestry, and imputed sex (**Methods:** *Polygenic risk score (PRS) prediction in UKB*). Polygenic scores of Hip Width:Height and TFA were associated with increased incidence of hip and knee OA respectively (p = 7.92 × 10^-5^, OR = 1.04; p = 1.73 × 10^-4^, OR = 1.04) in line with the phenotypic associations. In addition, we also saw significant association between back pain (both recorded on the ICD-10 code and self-reported) and Torso Length:Height (p = 5.59 × 10^-5^, OR = 1.05; p = 5.71 × 10^-6^, OR = 1.02) (**Table S21**). Neither OA nor musculoskeletal pain phenotypes we tested were significantly associated with overall height in this analysis (phenotypic associations: 1.10 × 10^-2^ < p < 8.51 × 10^-1^; PRS associations: 2.17 × 10^-3^ < p < 3.88 × 10^-1^) except for polygenic risk score (PRS) of height and back pain (p = 5.76 × 10^-10^) (**Table S20 and Table S21)**. In Genomic SEM analyses, we observed similar patterns of genetic associations with musculoskeletal diseases at the level of general genetic factors (**Table S22**, **Fig. S13)**

Taken together, these analyses suggest that increases in the length of skeletal elements associated with the hip, knee and back as a ratio of overall height were exclusively associated with increased risk of arthritis and pain phenotypes in those specific areas.

### Evolutionary Analysis

As human skeletal proportions are an important part of our transformation to bipedalism, we next investigated whether variants associated with skeletal proportions have undergone accelerated evolution in humans in two ways. First, following a procedure by Richard et al. (*18*) and Xu et al. (*56*), we examined if genes associated with skeletal proportions overlapped human accelerated regions (HARs) more than expectation. HARs are segments of the genome which are conserved throughout vertebrate and great ape evolution but strikingly different in humans. We generated a null distribution by randomly sampling regions matched for overall gene length (**Fig. 5A**, **Methods:** *Enrichment analysis for HARs*). For comparison, we also performed the same analysis on summary statistics from the ENIGMA consortium (*57*), and several quantitative and common quantiative and disease traits from the UKB (**Table S23**). Genetic signals from several of the skeletal proportion traits, in particular arm or leg length, were significantly enriched in HARs (Arms:Legs, Humerus:Height, Arms:Height, Hips:Legs, Tibia:Femur, Hip Width:Height, had FDR-adjusted p < 0.05). We also observed nominal enrichment for traits related to hair pigmentation (FDR-adjusted p = 0.013), which has also changed dramatically compared with the great apes, and for schizophrenia (FDR-adjusted p = 1.61 × 10^-34^). However, no enrichment (FDR-adjusted p > 0.05) was observed for HARs in autoimmune disorders, cardiovascular disease, cancer, and overall height (**Fig. 5A**).

**Fig. 5.**
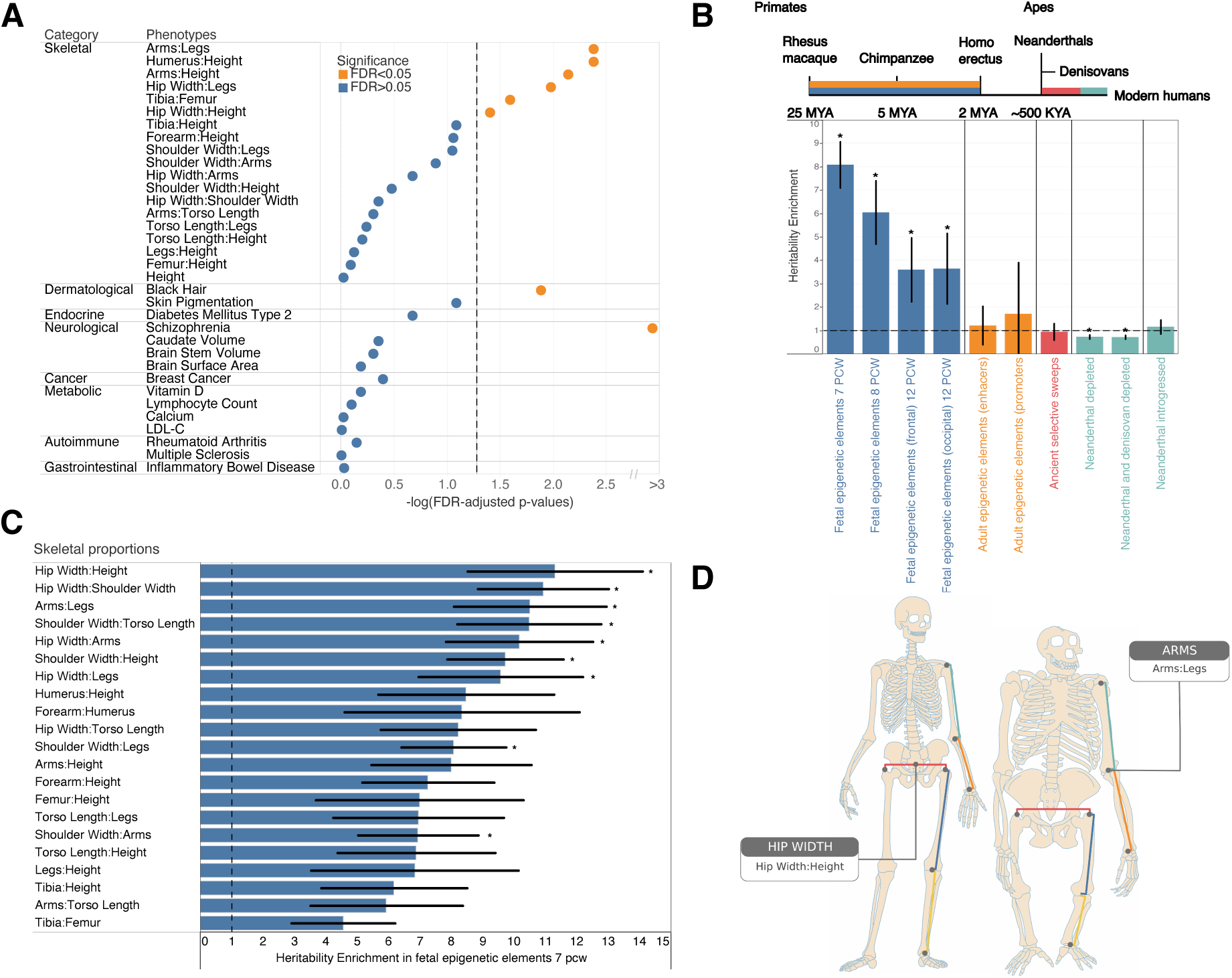
Evolutionary analyses. (**A**) P-values of enrichment for overlap of HARs with genes associated with skeletal proportions, autoimmune, dermatological, neurological, endocrine, gastrointestinal, metabolic, psychiatric, and cancer-related traits compared to randomly sampled genes of comparable length. Traits above the FDR-corrected threshold (0.05) shown in orange, and non-significant traits shown in blue. (**B**) Meta analysis of LDScore Heritability Enrichment across 21 skeletal proportion traits for different evolutionary annotations representing different divergence points in human evolution. Annotations represented in colors refer to fetal human-gained enhancers and promoters (*blue*), adult human-gained enhancers and promoters (*orange*), ancient selective sweeps (*red*), putatively introgressed variants from Neanderthals (*teal*), and genomic regions depleted in Neanderthal and Denisovan ancestry (*teal*). Blue and orange intervals mark epigenetic annotations while the other color intervals mark genetic annotations. Asterisks show significance at FDR < 0.05. A dashed line is drawn at y = 1 (no heritability enrichment). This analysis was jointly performed with all genomic annotations in the baseline LDv2.2 model. (**C**) Heritability enrichment analysis in human-gained enhancers and promoters at 7 PCW for each trait analyzed. Asterisks show significance at FDR < 0.05 across all genomic annotations and traits analyzed in this study. A dashed line is drawn at x=1 (no heritability enrichment). Error bars show 1 standard error around each estimate. (**D**) Arm:Leg ratio and Hip Width:Height are the only two skeletal traits that show significant enrichment in both types (HARs and heritability across differentially regulated regions at 7 PCW) of evolutionary analysis showing convergence of genomic change and some of the best-known anatomical differences between in humans and the great apes as shown by illustrations of ape and human skeletons.

Second, we examined heritability enrichment using LD score regression on genomic annotations reflecting divergence at different time points in human evolution (**Fig. 5B**) following an approach outlined in Sohail and Hujoel et al. (*58, 59*). These annotations include regions that differ in gene regulation between humans and primates through stages of early development (*60*), regions that differ in expression between adult humans and macaques (*61*), and regions that are enriched and depleted of ancestry from archaic humans (*62, 63*) (**Methods:** *LDScore heritability enrichment in regions of evolutionary context*). We then computed heritability enrichment, *h^2^*(*C*), that measures the proportion of heritability in an annotation set divided by the proportion of SNPs in the annotation (**Methods:** *LDScore heritability enrichment in regions of evolutionary context*). In our analysis we also simultaneously incorporated other regulatory elements, measures of selective constraint, and linkage statistics (baselineLDv2.2 with 97 annotations) (*59, 64–66*) to estimate heritability enrichment *h^2^*(*C*) while minimizing bias due to model misspecification.

Meta-analyzing across all our skeletal proportion traits we found enrichment in fetal human-gained enhancers and promoters in early time points (7, 8.5, and 12 post-conception weeks (PCW), *h^2^*(*C*) = 8.08, p = 5.91 × 10^-44^; *h^2^*(*C*) = 3.60, p = 2.55 × 10^-4^; *h^2^*(*C*) = 3.65, p = 3.55 × 10^-4^, **Table S24**) but not in adults suggesting that genes associated with skeletal proportions are differentially expressed in early development between apes and humans. While we acknowledge that the annotations of differentially regulated elements are from developing brain and not skeletal tissues, fetal human-gained brain regulatory elements and adult human skeletal regulatory elements are correlated at 58% (*58, 67*). Moreover, our observation of only observing enrichment in developing but not adult tissues suggests that enrichment is not driven by confounders of tissue type but by differences in development between the two species. As a second line of analysis, we also examined enrichment of individual traits across the different annotations controlling for multiple hypothesis correction at the level of FDR<0.05. 9 out of 21 of our skeletal proportion traits (Hip Width:Height, Hip Width:Shoulder Width, Arms:Legs, Shoulder Width:Torso Length, Hip Width:Arms, Shoulder Width:Height, Hip Width:Legs, Shoulder Width:Legs, Shoulder Width:Arms) were significantly enriched at 7 weeks PCW at FDR < 0.05 (**Fig. 5C, Table S25**). In addition, we saw depletion for enrichment in regions of the genome that were depleted for Neanderthal and Denisovan ancestry, particularly for overall leg length (*h^2^*(*C*) = 0.44, p = 5.89 × 10^-5^) (**Table S25)**. These results were consistent with other analysis which showed a depletion of Neanderthal informative markers in contrast with modern human mutations particularly for anthropometric traits (*68*) and are suggestive of purifying selection.

The proportion traits that were significantly enriched across both types of evolutionary analysis were associated with Arms:Legs and Hip Width ratios (**Fig. 5D**). These results suggests that specific skeletal proportions, but not overall height or several other quantitative and disease traits examined by us or (*58*) underwent human lineage-specific evolution since the separation of humans from the great apes.

## Discussion

In this study, we used deep learning to understand the genetic basis of skeletal elements that make up the human skeletal form using DXA imaging data in a large population-based biobank. We demonstrate that deep learning is useful not just in phenotyping individuals, but also as a tool for quality control at scale, including the capture of heterogeneous types of error modes. Our work also demonstrates the importance of having an interconnected dataset incorporating 3 different types of data - imaging, genetic, and health record/metadata - to best leverage biological insights - the scaling and resolution issues presented by the imaging data would have been impossible to correct for without external information about individual height in the biobank metadata. Through transfer learning we also show that accurate and replicable phenotyping can be achieved despite limited manual annotation. The fast and flexible architectures we present here can be deployed rapidly at population scale enabling their utility for automated phenotyping as imaging data becomes more integrated into large population biobanks.

Beyond methodological improvements for biobank-scale analysis, our results provide new insights into musculoskeletal biology. Despite over a century of work in genetics investigating the development of limbs and the overall body plan, beginning first in invertebrates and then later in vertebrate model organisms, a comprehensive genetic map of variation that shapes the overall skeletal form has been absent. More importantly, how the expression of various genes regulates modular development of the forelimb, hindlimb, and other long bones has not been fully characterized. Additionally, the broad set of genes and genetic variants that are responsible for the morphological changes in body proportions that allow us to walk upright has remained unknown. To the best of our knowledge, our work provides the largest genome-wide genotype to phenotype map of skeletal proportions in any vertebrate and lays the foundation for future functional assays of the genes discovered to understand how they contribute mechanistically to overall phenotype.

The moderate genetic correlations (a maximum of 0.55) observed between skeletal proportions indicates genetic sharing, particularly among limb length traits, while also highlighting the unique biology behind the growth of each element. Our results for genetic correlations are in line with artificial selection experiments in multiple mouse lines showing that selection for tibia length increased the trait by more than 15% across 14 generations but did not result in significant change in overall body mass (*69*) - a trait highly correlated with body width (r_g_ = 0.25, p-value = 1 × 10^-21^) but not limb length (r_g_ = -0.01, p-value = 0.53) proportions. Thus, these genetic correlation and factor analysis models provide insight into constraints placed on the evolutionary trajectory of the skeletal form both in humans and in vertebrates more broadly.

One important issue that affects the interpretation of our results is the normalization for height for each skeletal length measure we obtained. We did this to look at our primary outcome of interest, skeletal proportions that are independent of height. Several papers have cautioned that the interpretation of associations studies performed with adjustment should be carefully considered (*70, 71*). While this issue affects virtually every genome-wide association study that uses age as a covariate in the model (where age is a proxy for survivability - a complex trait with a heritable basis), our analysis is most similar to GWAS conducted for BMI, also a trait where body weight is computed as a proportion of height. Our results largely showed consistent direction of effect for loci before and after height adjustment. This suggests our GWAS for skeletal proportions are largely identifying loci that are directly associated with overall length of that particular skeletal element. However, a minority of these signals could still arise from pleiotropic increases or decreases in other skeletal elements that affect overall height. Thus, in interpreting our results, it is important to only view each of our phenotypes as proportions of height rather than directly associated with particular skeletal element lengths themselves.

Epidemiological studies indicate that OA of the hip as well as the knee frequently occur in the absence of OA in each other as well as other large joints, suggesting that local factors are important in OA pathogenesis (*72–77*). Specific abnormalities in skeletal morphology are now recognized as major biomechanical risk factors for the development of OA. Based on improved understanding of these morphological variations, parameters have been introduced to quantify them and enable classification of patients presenting with early OA (*78–83*). The findings presented here of the association between skeletal proportions, but not overall height, and joint-specific osteoarthritis highlight the biomechanical role these proportions play in shaping stresses on the joints themselves and highlight unique risk factors of clinical relevance.

Across both types of evolutionary analyses, the most significant skeletal proportion traits were those associated with the proportions of arms and legs, as well as proportions of hip width. These results are concordant with some of the most striking morphological differences between the two species being arm-to-leg ratio, as well as the change in the human pelvis which has allowed for a transition from knuckle-based walking to bipedalism (**Fig. 5D**). In addition, our results for heritability depletion in leg length-related traits in regions depleted for archaic ancestry are consistent with skeletal differences between anatomically modern humans and Neandertal s – who have shorter total limb length relative to body size and lower distal to proximal limb proportions (*84, 85*). Numerous studies have proposed a thermo-regulatory hypothesis that accompanied the primary biomechanical energy efficiency hypothesis for the evolution of these traits in early homonin evolution as well as to explain differences in anatomy between humans and Neanderthals (*86, 87*). However, only one extremely small sample study of 20 individuals, has been conducted to attempt to test these thermo-regulatory theories (*88*). Here, we conducted large sample size genetic correlation analysis between skeletal proportions and basal metabolic rate as well as whole-body fat-free mass in humans using genetic correlation (**Methods**: *Genetic correlation of skeletal proportions with external phenotypes*). We found that increased Arms:Legs ratio was associated with lower basal metabolic rate and lower whole-body fat-free mass (p = 9.37 × 10^-16^; p = 4.05 × 10^-16^) in line with the theory that these changes in early human evolution would have also increased heat dissipation in early hominins (**Table S26**). Similarly, increased distal to proximal limb proportion (Tibia:Femur) was associated with increased basal metabolic rate and increased whole-body fat-free mass (p = 2.23 × 10^-14^; p = 1.18 × 10^-14^) also consistent with the theory of Neanderthals being selected for survival in cold climates adding additional support to a thermo-regulatory mechanism for evolution of these traits (**Table S26**).

To our knowledge, these results provide the first genomic evidence of selection shaping some of the most fundamental anatomical transitions that have been observed in the fossil record in human evolution - changes in the overall skeletal form which confers the unique ability of humans to walk upright.

## Materials and Methods

### UKB participants and dataset

All analyses were conducted with data from the UKB unless otherwise stated. The UKB is a richly phenotyped, prospective, population-based cohort that recruited 500,000 individuals aged 40–69 in the UK via mailer from 2006 to 2010 (*41*). In total, we analyzed 487,283 participants with genetic data who had not withdrawn consent as of April 9, 2021, out of which 42,284 had available DXA imaging data. Access was provided under application number 65439. The baseline participants metadata including age and sex and other variables related to our study are in **Table S1**.

### Dual-energy X-ray Absorptiometry (DXA) Imaging

The UKB has released DXA imaging data for a total of 50,000 participants as part of bulk data field ID (FID) 20158. The DXA images were collected using an iDXA instrument (GE-Lunar, Madison, WI). A series of 8 images were taken for each patient: two whole body images - one of the skeleton and one of the adipose tissue, the lumbar spine, the lateral spine from L4 to T4, each knee, and each hip. Dual-energy X-ray absorptiometry (DXA) images were downloaded from the UKB bulk data FID 20158. The bulk download resulted in 42,284 zip files, each corresponding to a specific patient identifier otherwise known as each patient’s EID, and each file contained several DXA images of the patient as described above. All images were exported and stored as DICOM files which were later converted to high resolution JPEG files for image analysis and quantification.

### Phenotype and clinical data acquisition

The binary classification of patient disease phenotypes was obtained from a combination of primary and secondary ICD-10 codes (FID 41270) and the non-cancer self-assessment (FID 20002). Self-assessment codes were translated to three-character ICD-10 codes (Coding 609) and ICD-10 codes were truncated to only be the initial three characters. Patients received one if a disease code appeared in either self-assessment visit or their hospital records and zero otherwise. Reports of a fracture from a simple fall (FID 3005) or within the last 5 years (FID 2463) of any visit (instance 0 to 3) was considered a case. Falls in the last year (FID 2296) from any visit were considered a single case, regardless of a patient having more than one fall within the year. Our classification of fractures and falls increases case counts while excluding any childhood incidence. **Table S27 and Table S28** contain all ICD-10 and FID codes we used in our analysis.

### Computing infrastructure

All analysis was carried out on the Corral and Frontera system of the Texas Advanced Computing Cluster. The deep learning analysis was carried out on NVIDIA Quadro RTX 5000 GPUs using the CUDA version 11.1 toolkit.

### Classification of DXA Images by body part

Each individual had a DXA image folder containing up to 8 different body parts. In order to check the labels of these body parts that were defined using their file name, we built a convolutional neural network (CNN) to sort the images by body part through the use of a multi-class classification model using Python libraries FastAIv2 (*89*) and pydicom (*90*). We selected 1,600 total images - around 200 images per body part - and randomly split them into 1,280 images for training and 320 images for validation. These training and validation images were labeled by hand and cross-referenced with the label associated with the DXA image metadata. Training was run for 3 epochs using ResNet-152 (*40*) as the CNN and we obtained a validation accuracy of 100%. This classifier was run on all DXA images obtained from the UKB and 150 images were discovered that were correctly identified by the classifier, but incorrectly labeled in the DXA metadata. These images were removed from all future analyses. After sorting and removal of images, we were left with 42,228 full skeleton x-rays (**Table S29**).

### Removal of poorly cropped X-rays

After we determined the final set of full body x-ray images, we performed additional quality control to remove images that were poorly cropped and cut off parts of the arms on the image. To do this we created a binary classifier using FastAIv2 to differentiate between cropped and non-cropped images. 600 images were selected by hand - evenly split between cropped and non-cropped images - to use for training and validation. These images were randomly split into 480 training images and 120 validation images. The images were also all labeled by hand and trained for 30 epochs using a CNN with a ResNet-152 architecture. The final results had an accuracy on validation data of 100% on validation data. Removal of all the cropped images resulted in a total of 39,644 full-body images that we used for analysis (**Table S29**).

### Image standardization

From the pool of remaining full-body x-ray images, we discovered that the images varied in both pixel dimension and background. Broadly, the images fell into two main categories: (a) images that were on a black background with sizes between 600-800 by 270 pixels and (b) images on a white background with sizes between 930-945 by 300-370 pixels. The overall distribution of images by pixel ratio and an example of each type of image is shown in **Table S30** and **Fig. S1**. In order to process these images and remove effects of scaling and resolution change during the deep learning process, we chose to pad all the images to be of consistent size. We removed images that had sizes far out of the normal range and processed each of the two categories of images separately. The black background images were padded equally on all sides of the image to a final resolution size of 864×288 pixels while the white background images were padded in the same fashion to a resolution size of 960×384 pixels. We carried this out by converting each of individual DICOM files obtained from the UKB into numpy arrays and added additional rows and columns of black or white pixels as appropriate using standard functions from numpy (*91*), scipy (*92*), and skimage (*93*). These final resolution sizes were chosen based upon image size requirements for our deep learning model for landmarking and image quantification. Padding and removing individuals with sizes that did not fit into the two major categories resulted in a final total of 39,469 images - 21,981 images of 864×288 and 17,488 images of 960×384. In our deep learning model for landmarking, we trained two separate models, one for each pixel ratio, as these images were different not just in their size but also in their background.

### Manual annotation of human joint positions

To train our deep learning model, we manually annotated a total of 297 images (with 148 images padded to 960×384 pixels on a white background, and 149 images padded to 864×288 pixels on a black background). We used 100 images of each type for training and the rest for validation. The images that were chosen for this training dataset had an equal number of male and female individuals, had equal numbers of individuals who had an OA diagnosis in their ICD-10 codes, were from the white British population group (as determined by genetic PCA), and sampled equally across the age distribution of the UKB cohort. Out of the 297 total images, 10 images were duplicated in each of the image sizes to measure the replicability of our process. We used a single human annotator for all training data and provided an initial dataset of 317 (297 +2×10 duplicate images) without the annotator’s knowledge. We used a standard annotation scheme in computer vision, the Common Objects in Common (COCO) (*37*) scheme which provides a rubric for joint landmark estimation on the human body. The positions in the body we chose to annotate were the: left eye, right eye, left shoulder, right shoulder, left elbow, right elbow, left wrist, right wrist, left hip, right hip, left knee, right knee, left ankle, and right ankle, which have been long used in benchmarking analysis of human pose estimation - designed to label joints and other features in natural images of humans. For annotating each of these landmarks, the locations specified below were chosen because they were the easiest and most consistent to identify across all the images, which featured slightly different poses. The center of the orbit was chosen to be labeled for each of the eye landmarks. The center of the head of the humerus was chosen to be labeled for each of the shoulder landmarks. A location near the elbow joint closest to the olecranon fossa was chosen to be labeled for each of the elbow landmarks. A location near the scaphoid bone near the wrist was chosen to be labeled for each of the wrist landmarks. The topmost tip of the femur was chosen to be labeled for each of the hip landmarks. The middle of the femur and tibia was chosen to be labeled for each of the knee landmarks. The point where the ends of the tibia, fibula, and talus converge was chosen to be labeled for each of the ankle landmarks. An example of the annotation of one image is shown below in **Fig. 1B** with landmarks placed at each of the locations listed above.

We measured the replicability of our annotations by taking the Euclidean distance of pixels between the corresponding key points across 10 images that were duplicated amongst the 864×288 image set without knowledge of whether the image was a duplicate. Our replication analysis of 10 duplicate images was under 3 pixels across the different points that were estimated. Across the body parts, the farthest deviation across annotations was seen in the ankles, but the mean replicability across 10 images was under 3 pixels for both the right and left ankles. **Table S31** shows the mean pixel differences of 10 images across these duplicate annotations in the 864×288 dataset.

### A deep learning model to identify landmarks on DXA scans

In order to perform joint/landmark estimation on the entire UKB skeletal X-ray dataset using our manually annotated training data, we compared two different neural network architectures that have been used for previously for landmark detection on human subjects, ResNet-34 (*40*) and HRNet (*35*) **Fig. S2**. To arrive at the best possible architecture and training process for our task we utilized transfer learning and began with pre-trained models which were trained first on ImageNet and then trained on 123,847 images from the COCO dataset (*37*) which have been annotated with landmarks of humans performing tasks in various natural settings such as playing sports, driving, or seated indoors. On these pre-trained models, we adopted two approaches, one where we fine-tuned these networks using our manual annotation on the fully body X-rays as above, and one where we did not perform additional fine tuning/training. In addition, for each architecture and each choice of training approach, we varied the heatmap resolution size (an area around each landmark that was predicted) and overall image size. Our results across architectures, including/not-including fine-tuning, heatmap resolutions and image size are in **Table S2**. Based on these results we found that larger image input sizes and heatmap resolution sizes performed better and that accuracy even with smaller input sizes were over 95% across body parts. We also found that the HRNet based architecture performed better across parameter choices and therefore for the final analysis, we used the HRNet model that had been pretrained on the COCO dataset, and then fine-tuned on our manually annotated images. We also used the two post-padded image sizes and used the largest heatmap size available to us. The 864×288 and 960×384 models were run at a batch size per GPU of 12 and 8 respectively for 210 epochs.

### Validation metrics comparing automated annotation to manual annotation

As an initial examination of the performance of our model, we visualized the location of the landmarks on the original X-ray image to confirm that our labeling was qualitatively accurate on both imaging modes. We quantified the pixel-level accuracy of our model by comparing the automated annotation versus a set of 50 images that were manually annotated and computed landmark differences in pixels for each landmark. The results of this analysis for both image modes are shown in **Table S5 and Table S6**. Following the training process, we deployed our model on all 39,469 images of full body X-rays from the UKB.

### Adjusting for scaling differences across imaging sizes and modes

A major issue in combining our analysis across input pixel ratios was that these pixel ratios represented different resolution scalings, perhaps due to distances that the scanner was held above the patient (**Fig. S3**). That is, in one image a pixel could represent 0.44 cm and in another 0.46 cm. To control for this scaling issue and to standardize the images, we chose to regress height measured directly on our image using the midpoint of the eyes and the midpoint of the two ankle landmarks that could be taken across all image pixel ratios and overall height (FID 50) computed externally from the UKB (**Fig. S4**). While the height measure we utilized did not include the forehead, it was a relative measure that we used to obtain a scaling factor for each image pixel ratio that we could for normalization. Measurement error of individuals either in our image-based height measure or as reported in the UKB is not expected to affect our conversion from pixels to cms as we are regressing over many individuals. Importantly, we validated this regression and normalization using duplicate individuals taken by different scanners, imaging modes and technicians (**Fig. 1D-F**).

### Obtaining skeletal element length measures

From each of the 14 landmarks, we generated a total of seven skeletal length measures and one angle measure in pixels which we converted to centimeters using coefficients from the regressions with height. 4 of these measures were for each bone that makes up the limbs, the humerus, forearm, femur, and tibia. We averaged these lengths across the left and right side of the body for all analysis. We generated measurements of shoulder width and hip width using the shoulder and hip joint landmarks. From the midpoint of the shoulder and hip landmarks we also generated a torso length measure. Finally, we measured the angle between the femur and the tibia by obtaining the average across two legs, with angles greater than 180 corresponding to knees bent outward (bow-knees) while angles less than 180 correspond to knees bent inward (knock-knees). For all measurements, mean and standad deviations are shown in **Table S4**.

### Removal of image outliers

We removed individuals who were more than 4 standard deviations from the mean for any skeletal length measure from the analysis. Examination of these outliers by comparing left and right symmetry as well as comparison of other body proportions revealed a heterogeneous set of issues that were associated with the poor prediction by our deep learning model. In some cases, individuals had a limb, or another body part amputated. Some poorly classified images were individuals who had had major hip or knee replacement surgery or had various implants that were causing incorrect model landmark prediction (**Fig. S5**). Another class of outlier images were those that were too poor in quality for any landmarking of any of the points on the image. A distribution of outlier individuals as well as possible reasons for their removal is in **Table S32.**

### Obtaining a set of body proportion traits from raw length measures

From the seven skeletal length measures we calculated 21 different body proportion measures representing ratios of one measure with the other or with overall height. A list of these proportions can be found in **Table S3**. Ratios were generated by dividing smaller lengths by larger lengths to generate body proportions as a phenotype.

### Correlations of skeletal proportions with age and sex

Each of the skeletal proportion phenotypes were correlated with age, and p-values were calculated to see how body proportions are affected by age. Furthermore, we carried out t-test analyses on each phenotype to look at differences in body proportions based on sex. Both analyses were carried out on white British patients only (n = 31,221). Results are shown in **Table S8 and Table S9.**

### Participant data quality control

For all genome-wide association analyses, we filtered the participants with correctly labeled full body DXA images (FID 20158 and 12254) to individuals from just Caucasian individuals (FID 22006) from the white British population as determined by genetic PCA (FID 21000). We removed individuals whose reported sex (FID 31) did not match genetic sex (FID 22001), had evidence of aneuploidy on the sex chromosomes (FID 222019), were outliers of heterozygosity or genotype missingness rates as determined by UKB quality control of sample processing and preparation of DNA for genotyping (FID 22027), had individual missingness rates of more than 2% (FID 22005), or more than nine third-degree relatives or any of unknown kinship (FID 22021). In total 31,221 individuals remained.

### Genetic data quality control

Imputed genetic data for 487,253 individuals was downloaded from UKB for chromosomes 1 through 22 (FID 22828) then filtered to the quality-controlled subset using PLINK2 (*94*). All duplicate single nucleotide polymorphisms (SNPs) were excluded (--rm-dup ’exclude-all’) and restricted to only biallelic sites (--snps-only ’just-acgt’) with a maximum of 2 alleles (--max-alleles 2), a minor allele frequency of 1% (--maf 0.01), and genotype missingness no more than 2% (--maxMissingPerSnp 0.02). In total 8,638,168 SNPs remained in the imputed dataset. Non-imputed genetic data (genotype calls, FID 22418) did not contain duplicate or multiallelic SNPs but were filtered to the quality-controlled subset; 652,408 SNPs remained.

### Heritability analysis and GWAS

GWAS was performed with BOLT-LMM (*95*). Heritability, genetic correlation, LD Score and PCA analyses were carried out with the non-imputed data, but the imputed data was used for the final association testing using a linear mixed model. LD Score v1.0.1 was used to compute linkage disequilibrium regression scores per chromosome with a window size of 1 cM (*44*). PLINK2 --indep-pairwise with a window size of 100 kb, a step size of 1, and an r^2^ threshold of 0.6 was used to create a list of 986,812 SNPs used as random effects in BOLT-LMM. Covariates were the first 20 genetic principal components provided by UKB (FID 22009), sex (FID 31), age (FID 21003), age-squared, sex multiplied by age, sex multiplied by age-squared, and estimated height from eyes to ankles. In addition, the DXA scanner’s serial number and the software version used to process images were combined into one covariate, resulting in 5 factor levels.

The heritability of each phenotype was assessed with non-imputed data using BOLT-REML with the same covariates. SNPs in each resulting GWAS were clumped using --clump with a significance threshold of 5.0 × 10^-8^, a secondary significance threshold of 1.0 × 10^-4^ for clumped SNPs, an r^2^ threshold of 0.1, and a kilobase window of 1 Mb. SNPs were assigned to genes with --clump-verbose --clump-range glist-hg19 downloaded from PLINK gene range lists (*96*). The genomic inflation factor of each phenotype was assessed in R version 3.6.1 as the ratio of the median of the observed chi-squared distribution (an output of BOLT-LMM --verbose) to the expected median of the chi-squared distribution with one degree of freedom. We examined the pairwise genetic correlation of traits using GCTA version 1.93.2 beta for Linux (*97*). (*97*)We created the genetic relationship matrix for our quality-controlled subset but without any related individuals (21,248 total individuals remained) and a minor allele frequency of 0.01, then ran GCTA for each phenotype pair with the first ten genetic principal components provided by UKB (FID 22009).

We also estimated heritability using LDSC (*44*) and found similar but slightly lower heritabilities (30-60%) compatible with either reduced power for LDSC based methods or due to assortative mating increasing the estimate for REML-based methods (*98, 99*).

A major contribution of noise in GWAS comes from measurement error. We wanted to see if heritability estimates of height measured in pixels calculated directly on the skeleton which and therefore have lower measurement error could be greater than measurements carried out externally by UKB. To do this we compared the heritability of height computed in three ways: raw pixel lengths, FID 50 standing height, and FID 12144 height from the first imaging visit on the 864-pixel image size subset of 16,623 imaged white British individuals all meeting imaging and genetic QC outlined above. FID 12144 only reports height in integer cm, whereas FID 50 reports to the first decimal. All these 16,623 individuals were imaged at the same pixel ratio and thus were unaffected by resolution scaling issues. The heritability measures and standard deviations in brackets were as follows:

Heritability of height measured in pixels 0.75 ± 0.03

Heritability of height from FID 50 = 0.74 ± 0.03

Heritability of height from FID 12144 = 0.67 ± 0.03

The heritability of FID 12144 was the lowest of all measures (h^2^=0.67, se=0.03) as expected from the course measurement. However, the difference between height measured externally by the biobank (FID 50) and our pixel-based measurements was non-significant suggesting that we did not see improvements in measurement error compared with external measurements. One possibility could be that height, unlike waist size or hip size, is well-measured externally and that having skeletal measures does not add significant improvement to measurement accuracy.

### Adjusting for height correlation in GWAS using ratios

We were broadly interested in human body proportions, that is, how various lengths in our body change as a proportion of overall height. However, the common denominator of height used in these proportions might induce spurious correlations across these proportion phenotypes (*100*). In practice this is less of a problem as the overall variation in height was only a small portion of the overall height. However, in carrying out correlation analysis we attempted to normalize for height in three different ways. First in examining phenotypic correlations we show that residualizing each measure by height and then taking correlations does not induce spurious correlation. To do this, we simulated data of 31,221 individuals under the mean and standard deviation of femur length and humerus length as well as height on a standard normal distribution. As these three measures were randomly generated, we do not expect to see correlation between them. However, on taking ratios with height we observed a correlation of 0.25 between Humerus:Height and Femur:Height, but a correlation of 0.00012 when examining correlation on residualized humerus and femur measures.

Second, we attempted to carry out GWAS in three different ways and used those to compute genetic correlations between skeletal traits. First, we divided each trait by the overall height of each person and carried out GWAS on the proportion phenotype. Second, we tried added height as a covariate as part of the GWAS along with the other covariates. Third, we regressed each trait on the overall height and then performed a GWAS on just the residuals of that regression. Genetic correlations between all three GWAS results were highly correlated with one another (r_g_ 0.96 (ratio with height covariate), 0.97 (ratio with residual), and 0.97 (residual with height covariate) for the 3 pairs (**Table S33 and Table S34**). We then compared genetic and phenotypic correlations across skeletal proportion phenotypes controlling for height as simple ratios and then as residuals and found they were highly similar (Overall Pearson correlation r between the two matrices was 0.969) (**Fig. S6 and Fig. S7**).

For simplicity we decided to use the first approach of obtaining each trait as a ratio of height for all analyses other than examining genetic correlations between two phenotypes that were both proportions of height where such an approach could lead to spurious correlations. For example, when looking at the genetic correlation between Femur:Height and Humerus:Height we computed phenotype and genotype correlations on the residuals of regressing femur on height rather than just examining correlations on the raw ratios.

### Sensitivity analysis for height adjustment

To test for possible bias due to running GWAS using bone length and body width measurements as proportions of height (sometimes called collider bias), we carried out sensitivity analysis outlined by Aschard et al. (*70*) to test the effect of each SNP in the same sample population on the raw phenotype (femur), the covariate itself (height), as well as the adjusted analysis (Femur:Height) (**Table S35)**. To verify this, we conducted a GWAS of femur length, height, and Femur:Height as well as torso length and Torso:Height for 31,221 individuals on more than 7 million SNPs. We observed that for both our torso and femur phenotypes, we see that the vast majority (>95%) of genome-wide significant signals are in the same direction as the non-adjusted phenotypes (**Fig. S8**).

### Multivariate genetic architecture of skeletal endophenotypes

To investigate the joint genetic architecture of skeletal traits, identifying clusters of skeletal traits with a shared genetic component, and elucidating biological pathways of genetic risk for musculoskeletal diseases, we used genomic SEM to analyze the genetic factor structure of the limb and body measurements independent of height. We also analyzed associations of these factors to musculoskeletal disease. Links to details on case ascertainment, genotyping, and quality control are provided in **Table S36**. Inclusion criteria for summary data were: mean χ^2^ > 1.03, LDSC h^2^ Z-statistic > 2, and mean χ^2^ / LDSC intercept ratio > 1.02.

For LDSC genomic SEM analyses, the included SNPs were restricted to HapMap3 common SNPs (1,215,001 SNPs) (*101*). MHC region SNPs and SNPs with MAF < 1% or information scores < 0.9 were excluded. We first conducted exploratory modeling including SNPs on odd numbered autosomes. We reserved SNPs on even numbered autosomes for confirmatory modeling to assess model fit.

We employed the multivariate extension of LDSC to estimate SNP-based heritabilities and co-heritabilities across skeletal proportions in odd-numbered autosomes. The estimated LDSC S matrix containing the genetic variances and covariances was smoothed to the nearest positive definite matrix using the Higham algorithm (*102*). The maximum difference in Z statistics between the pre- and post-smoothed S matrix was 0.00001, suggesting very little distortion of the original matrix. The smoothed S matrix was then standardized to compute the genetic correlation matrix S_std_. We additionally compared the LDSC- and the GCTA-estimated genetic correlations.

Using the LDSC-estimated genetic correlation matrix, we conducted a Parallel Analysis (*103*) to determine the number of factors to retain in a subsequent Exploratory Factor Analysis (EFA). We compared the eigenvalues from the LDSC-estimated genetic correlation matrix to a distribution of eigenvalues from null correlation matrices (1s on the diagonal, 0s off the diagonal) sampled with random noise drawn according to the multivariate sampling covariance matrix, V_std_.

The genetic correlation matrix revealed substantial genetic sharing among the 9 skeletal traits, with varying degrees of genetic overlap across traits (**Fig. S9**). Arm- and leg-related traits showed substantial positive genetic correlations with each other. We found positive but modest genetic correlations among torso-related traits. Torso length and hip width presented negative genetic correlations with arm- and leg-related traits. Shoulder width presented negligible genetic correlations with arm-related traits, and small and negative genetic correlations with leg-related traits. There was a close correspondence between the GCTA and the LDSC-derived genetic correlations (r = 0.99, linear regression model intercept = < .001, linear regression model slope = 0.972, linear regression model R^2^ = 0.981, **Fig. S10**).

Results from the Parallel Analysis revealed two principal components from the LDSC-estimated genetic correlation matrix presenting eigenvalues exceeding 95% of the corresponding eigenvalues from the simulated matrices (**Fig. S11**), with the first principal component accounted for 54.28% of the genetic variance among skeletal traits. The second principal component accounted for an additional 15.66% of genetic variation. Based on results of the Parallel Analysis, a 2-factor EFA model with *promax* oblique rotation was fit to the LDSC-estimated genetic correlation matrix (**Table S37**).

In the 2-factor EFA solution, Factor 1 consisted of arm-related skeletal traits, including average arms, humerus, and forearm. Factor 2 was mostly comprised of leg-related traits, including average legs, tibia and femur. Torso-related traits presented cross-factor loadings on both the arms and legs factors. There was a medium-size correlation between the arms and legs factors (r_g_ = 0.51).

### Confirmatory models comparison within even-numbered autosomes

We next specified and compared the goodness-of-fit of three types of Confirmatory Factor Analysis (CFA) models within the even-numbered autosomes based on the 2-factor EFA solution for the odd-numbered autosomes (**Fig. S12**). Performing our exploratory analyses in an independent set of autosomes rather than the set in which we estimate model fits helps us to avoid inflation of goodness-of-fit that would otherwise result from estimating model fit in the same dataset on which the model was trained.

First, we fitted a factor model comprising two correlated factors of Arms and Legs with cross-factor loadings for the torso-related traits (Model A in **Fig. S12**). Next, we fitted a 3-factor CFA model consisting of three correlated factors of Arms-, Legs-, and Torso-related traits (Model B in **Fig. S12**). Finally, we fitted a bifactor model, comprising a common factor of genetic sharing among all phenotypes, and 2 specific groups factors of Arms and Legs accounting for the genetic variance unique to the arms and legs-related traits (Model C in **Fig. S12**). We selected the model presenting the better goodness-of-fit indices (i.e., highest CFI, and lowest χ^2^, AIC, and SRMR values), and applied it to the complete dataset including the 22 autosomes.

Goodness-of-fit indices for the 5 CFA models are reported in **Table S38**. The best fitting model was the bifactor model with 2 specific factors, Model C (χ^2^ [21] = 23233, AIC = 23281, CFI = 0.991, SRMR = 0.068).

An inspection of the standardized parameter estimates for Model C from even-numbered autosomes (**Table S39**) indicated that the Legs-specific factor was isomorphic with respect to the general factor of skeletal traits (λ_Legs,G_ = 0.999), with no significant genetic variance accounted for by the specific factor of leg-related traits (i.e., all shared genetic variance among the indicators of the Legs factor was accounted for by the common factor). Moreover, the residual correlations involving shoulder width were comparable in magnitude to the shoulder loadings on SK (**Fig. S12**), and the loadings for shoulder, hip, and torso on SK were also negative. These results hindered the substantive interpretation of the common factor of skeletal endophenotypes SK and provided no evidence for a specific factor of leg-related skeletal traits. Given these results, we decided to fit an additional model to provide a different conceptualization of the genetic covariance structure (Model D). Model D consists of 1) a leg factor that the other arm- and torso-related skeletal traits are residualized for (similar to a Cholesky decomposition), 2) an arms factor, and 3) three residual factors representing the genetic variance unique to hip, shoulder, and torso length. Model D exhibited good approximation to the genetic covariance structure with a more reasonable substantive interpretation (Model D fit indices: χ2 [24] = 118989, AIC = 119031, CFI = 0.953, SRMR = 0.074), and was thus carried forward for subsequent analyses. **Table S40** contains the parameter estimates for Model D in even autosomes.

We applied the preferred model (Model D) to the complete dataset including the 22 autosomes. The model fit the data well (χ^2^ [24] = 98520, AIC = 98562, CFI = 0.955, SRMR = 0.069). All arm and leg traits loaded positively and substantially on the common factor SK, whereas hip width, shoulder width, and torso length traits loaded negatively. We can conceptualize SK as a general propensity toward longer limbs relative to total height. We note that all skeletal traits were corrected for total height, which may help to explain the opposing factor loadings for limb and torso/width traits on the common factor. We can most straightforwardly conceptualize the general factor as representing overall limb length relative to height. The loadings of arms, forearm, and humerus on a separate Arms factor were also large, positive, and significant (**Table S41**). These results together indicate that 1) arms’ and legs’ skeletal length relative to height present substantial genetic overlap, 2) the genetic component of hip and shoulder width relative to height is mostly unique to each trait, 3) there is a specific source of genetic variation that is unique for arm relative length, and 4) there is strong and negative genetic associations between torso length and arms and legs’ skeletal structure relative to height.

### Genetic associations between musculoskeletal disease and skeletal trait factors

We conducted a series of genomic SEM models to assess the generality vs. specificity of the associations between the latent dimensions of skeletal structure specified in Model D and 18 musculoskeletal diseases (**Table S36**). To do so, we first estimated the observed effects of individual skeletal traits across musculoskeletal diseases using univariate regression models. We then calculated the effects mediated by common factor Model D, in which associations between musculoskeletal diseases and skeletal traits are fully mediated by the common factors SK and Arms, and the three unique factors of torso-related traits. Observed effects of individual skeletal endophenotypes on musculoskeletal diseases were estimated from univariate genomic regression models. Model-implied effects were obtained from multivariate genomic regression models, where the 5 genetic factors included in Model D (**Fig. S12**) are regressed on 18 musculoskeletal diseases (**Table S36**). We additionally calculated and compared the observed effects of the skeletal traits on a set of common musculoskeletal diseases in the UKB and FinnGen (i.e., coxarthrosis, gonarthrosis, dorsalgia, fibroblastic disorders, internal derangement of knee, intervertebral disk disorders, other joint disorders, rheumatoid arthritis, and spondylopathies).

We employed a correlated vectors method to quantify the correspondence between the observed and the effects implied by the 5 genetic factors specified in Model D, using correlations, Tucker’s congruence coefficients (CC), and linear regression models to quantify the correspondence between the two vectors of regression coefficients. We employed a Bonferroni correction for the p-values to control for multiple comparisons. We additionally conducted an outlier detection analysis to identify musculoskeletal diseases whose observed effects on individual skeletal traits differ substantially from those implied by the factor model, thus indicating potential specific pathways that are not mediated by the skeletal genetic factors.

Outliers were defined based on a standardized difference between model implied (**B_MI_** and observed **B_O_** effects, thus highlighting substantial deviations from perfect correspondence between observed and model-implied effects (blue dashed line in **Fig. S13**). First, we calculate the vector of absolute differences between standardized regression coefficients for model implied and observed effects as follows:

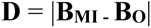

Then we standardized the vector of differences between **B_MI_** and **B_O_**, **D**, using the pooled standard deviation of model implied and observed effects:

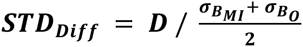

Musculoskeletal diseases were considered outliers if ***STD_Diff_***> 2 (see labeled skeletal traits across scatterplots in **Fig. S13**). **Table S42** contains the observed effects of musculoskeletal diseases on skeletal traits and the common factor estimates derived from Model D. **Fig. S13** displays the scatterplots of observed vs model-implied effects by common factor Model D between skeletal traits and musculoskeletal diseases.

The common genetic propensity toward longer relative limb length (SK) was associated with an increased genetic liability risk for arthropathies (0.266, p = .001), arthrosis (0.306, p = <.001), gonarthrosis (0.294, p < .001), hallux valgus (0.237, p = .008), internal derangement of knee (0.256, p = .001), and other joint disorders (0.228, p = 0.003). The vectors of observed and model-implied effects for associations involving these diseases were very similar in ordering (r range: 0.92 – 0.96) and magnitude (linear model intercept range: -0.01 – -0.03; linear model slope range: 0.99 – 1.07), presenting a close correspondence (CC range: 0.95 – 0.97), with no appreciable evidence of disease associations with individual skeletal traits operating through specific pathways not included in our modeling (**Fig. S13** and **Table S22**), indicating that the factors plausibly act on those diseases. On the contrary, the diseases for which there is lower correspondence (i.e., fibroblastic disorders, polyarthropathies, rheumatoid arthritis, soft tissue disorders, and systemic connective tissue disorders), tended not to have significant associations with the skeletal factors, suggesting more specific pathways of association with the individual skeletal traits. There was a moderate correspondence between the observed effects for the set of common diseases in UKB and FinnGen (r = 0.567, linear regression model intercept = -0.018, linear regression model slope = 0.570, R^2^ = 0.313, **Fig. S14**).

### Sensitivity analysis using height-residualized skeletal traits

We compared the genetic correlations among skeletal traits using height scaling (measurement/height) versus height residualization, where skeletal traits were first residualized by height before conducting GWAS. We additionally excluded the traits average arms and average legs from the confirmatory model of the preferred model (Model D). This further aspect of the sensitivity analysis allowed us to investigate the potential impact of collinearity among the arm- and leg-related skeletal traits on the model fit and the factor structure of the preferred model. High genetic overlap among perfectly collinear traits (i.e., average arm length = humerus length + forearm length; average leg length = femur length + tibia length) may inflate the proportion of shared genetic variance among such traits, thus potentially leading to spurious factor identification.

We found a close correspondence between the height-scaled and the height-residualized genetic correlations among skeletal traits (r = 0.98; **Fig. S7**) in both ordering and magnitude (linear model intercept: 0.002; linear model slope: 0.94). These findings suggest that both approaches produce very similar patterns of genetic overlap across skeletal traits. We next fitted the preferred confirmatory factor model (Model D) on the set of height-residualized genetic correlations, after excluding average arms and average length from the model (**Fig. S15**). To identify the arms-specific factor we constrained the factor loadings of the humerus and forearm to be equal. The model presented an adequate fit to the height-residualized data (χ2 [13] = 197.76, AIC = 227.76, CFI = 0.93, SRMR = 0.060).

### Sex-specific analysis

BOLT-REML was used to assess genome-wide SNP heritability of phenotypes in both sexes (N=31,221), males (N=15,279), and females (N=15,941) (**Fig. S16**). Standard errors for the ratio of sex-specific heritability to that of the heritability in both sexes was calculated using a 2nd order Taylor approximation for the standard error of a ratio of estimators of x and y, where x is a sex-specific heritability estimate and y is the heritability estimate across both sexes (*47*). We assessed male-female genetic correlation (r_g_) with GCTA bivariate GREML with the first ten principal components as covariates, no constraint on r_g_ (--reml-bivar-no-constrain), and against the hypothesis r_g_ is 0 (--reml-bivar-lrt-rg 0) (*104*).

Sex-specific GWAS were run in BOLT-LMM on a subset of 10,000 individuals per sex with a MAF of 0.1%, SNP missingness of 5%, and individual missingness of 2%. The first twenty principal components, age, age^2^, the serial number of DXA machine, and the software version for image processing were used as covariates. Using the GWAS performed in these samples, we computed out-of-sample polygenic risk scores for an independent sample of 5,000 males and 5,000 females. GWAS were clumped using an r^2^ threshold of 0.1 and a 250 kb threshold of physical distance for clumping, and a significance threshold of 1 × 10^-6^ was used to compute the PRSs in each sample. Next, we regressed the normalized PRSs (in standard deviations) obtained in each sample with the skeletal proportion phenotypes as a function of height (e.g., the ratio of average tibia length to calculated height) **Fig. S17**. From the estimates obtained in this analysis, we computed the ratio of the effect of the polygenic score on the trait (± 2 standard errors). This was computed as the ratio of the effect in the male samples to the effect in females across the skeletal proportion traits. We derived the standard errors for the ratio of male to female variance using the 2nd order Taylor approximation for the standard error of each sex, assuming independence between the estimated values for males and females (as they were obtained from independent sampling distributions).

### Clumping, independence analysis and removing previous height associated loci

To obtain a set of independent SNPs associated with each skeletal proportion phenotype, we first performed clumping analysis for each phenotype using plink and assigned SNPs to genes with --clump-verbose --clump-range glist-hg19 with an r^2^ window of 0.1 and a 1 Mb threshold of physical distance for clumping. We downloaded gene ranges from plink for hg19 (*105*).

Following clumping, we looked at a subset of 8 phenotypes, 7 limb and body lengths and widths regressed against height as well as TFA and combined the significant SNPs across the chosen phenotypes resulting in 212 unique SNPs. Overlapping clump regions were unioned using BEDtools (*106*). The --indep function in PLINK was used to prune out SNPs that were in approximate linkage disequilibrium with each other, leaving only independent SNPs (*105*). This function was carried out on the 212 SNPs chosen, resulting in 179 independent SNPs remaining. We then removed any of the 179 SNPs that were also found to be significant in a GWAS for height with greater than 10 times our sample size (Neale lab height GWAS), resulting in 102 SNPs remaining (**Table S43)**. The genes associated with each SNP as determined earlier by the clump range function in PLINK are also listed as well as each phenotype that each SNP was found to be significantly associated with.

### Functional mapping and gene enrichment analysis

For this analysis, out of the 23 phenotype GWAS results, we looked at the subset of phenotypes that were either limb or body lengths as a ratio of height which resulted in 7 phenotypes (Forearm:Height, Humerus:Height, Tibia:Height, Femur:Height, Hip:Height, Shoulder:Height, and Torso:Height) as well as the TFA. Using the GWAS output for each phenotype, we took the lowest p-value associated with each SNP to generate a combined GWAS output file across phenotypes. We then ran FUMA (*48*) without any predefined lead SNPs on a sample size of 31,221 individuals. GENE2FUNC was run with all types of genes selected as background genes using Ensembl v92 with GTEx v8 gene expression data sets.

### OMIM gene set enrichment analysis

We used FUMA (*48*) to generate gene level p-values from SNP p-value data. We then used Mare thought to beAGMA (*107*) gene set enrichment analysis to examine enrichment in 701 genes associated with abnormalities in skeletal growth in OMIM (*49*).

### Transcriptome-wide associations (TWAS)

We conducted a TWAS on 8 skeletal proportions to link imputed cis-regulated gene expression taken from expression quantitative trait locus (eQTL) data in skeletal muscle tissue with increased bone lengths. We carried out this analysis using FUSION (*108*) which also provided precomputed transcript expression reference weights for skeletal muscle tissue (n = 7408 genes). The analysis was run only on GTEx v7 muscle skeletal genes with significant heritability on the default FUSION settings as recommended by the authors of FUSION.

### Transcriptome analysis

To connect the genetics of skeletal proportions and growth plate biology, we looked for enrichment of genes associated with our skeletal proportion GWAS in gene expression data in three dissected layers of murine newborn tibial growth plate following an analysis described in Renthal et al. (*109*). Specifically, we were interested to see if we could identify which layers of the growth plate (i.e., the resting (round), proliferative (flat) or hypertrophic layer) would associate with increased limb length. The previous analysis in Renthal et al. used only overall height GWAS to examine these but we were interested to see if specifically obtaining GWAS for each limb proportion would provide additional insights. We downloaded microarray data of mouse tibial growth plate dissections from GEO data repository GSE87605 and normalized the data using Robust Multiarray Averaging (RMA) with the affy (version 1.72.0) package in R (version 4.1.3). Mouse gene IDs for each microarray probe were obtained from the GEO feature data for the Affymetrix Mouse Genome 430 2.0 Array. Mouse genes were then converted to human genes using the biomaRt (version 2.50.3) package in R (*110*). A specificity score for each growth plate (epiphyseal) layer was calculated as the proportion of total gene expression found in each layer. A score of 0 meant none of the total gene expression was found in the layer while a score of 1 indicated that all gene expression was found in that layer. We then carried out MAGMA gene property analysis to examine enrichment between genes expressed in particular growth layers and each skeletal proportion. However, unlike enrichment seen in Renthal et al. for overall height, we saw no significant enrichment for growth plate layers using our skeletal proportion GWAS after Bonferroni correction for the number of trait and layer pairs. In **Table S44** we report the results for all layer and proportion pairs for this analysis.

As different long bones differ dramatically in overall size, we examined whether we could correlate our GWAS results with RNA-Seq data comparing gene expression with age (1-vs 4-week-old mouse) in longer bones (tibia) and short bones (phalanx). To do this, we downloaded RNA-seq data from 1 and 4-week-old mouse tibial growth plates as well as 1-week-old mouse tibial and phalanx growth plates from GEO data repository GSE114919 (*111*). The data were normalized before upload to GEO, and mouse genes were converted to human genes using the biomaRt (version 2.50.3) package in R. The fold changes for each gene from 1-versus 4-week-old tibial growth plates and tibial versus phalanx growth plates were then calculated.

We used gene property analysis in MAGMA (version 1.08) to determine associations between genes implicated in 7 of our skeletal proportions GWAS (Forearm:Height, Humerus:Height, Tibia:Height, Femur:Height, Hip:Height, Shoulder:Height, and Torso:Height) and genes expressed in various bone layers and time points. Gene level p-values for our skeletal phenotype GWAS were first calculated using the positional mapping tool with default settings in SNP2GENE (version 1.3.7) (*48*). We then ran MAGMA’s gene property analysis method, which performs a one-sided association test between a covariate and phenotype. We used bone layer specificity score and RNA-seq fold change values as our covariates and used various skeletal traits as phenotypes. We also carried out this analysis using a GWAS on height as measured in our DXA image population as well as a GWAS on height across the UKB population carried out by Neale et al., as controls.

### Phenotypic association of skeletal phenotypes with musculoskeletal disease

To examine correlations between our skeletal phenotypes with musculoskeletal disease, musculoskeletal or connective tissue diseases related to the hip, knee, and back we obtained data from UKB Chapter XIII (FID 41270) ICD-10 codes as well as self-reported pain phenotypes (FID 6159) for the hip, knee and back. We then regressed the binary outcome of disease or reported pain against skeletal proportions controlling for clinically relevant covariates that are known to affect OA (*112*) including age, sex, diet, BMI, and other factors. A full list of variables we controlled for are reported in **Table S45**. All covariates were obtained from the notated FIDs in the UKB in **Table S28**. After running the regressions, we used Bonferroni correction for significance at the level of the total number of disease/pain traits multiplied by the total number of skeletal phenotypes.

### Polygenic risk score (PRS) prediction in UKB

This analysis only utilized the ∼300,000 white British individuals who were not included in our imaging dataset for which GWAS was conducted. Prior to testing for associations, on these individuals, we applied stringent sample quality control steps to infer global ancestries and exclude related and low-quality samples. We leveraged filters performed at the Wellcome Trust Center for Human Genetics, Oxford, UK. Filters included removing closely related individuals, individuals with sex chromosome aneuploidies and individuals who had withdrawn consent from the UKB study. To minimize the impact of confounders and unreliable observations, we used a subset of individuals that (1) had self-reported white British ancestry, (2) were used to compute principal components, (3) did not show putative sex chromosome aneuploidy.

Outcomes were pre-processed with the open-source software tool PHEnome Scan ANalysis Tool (PHESANT) (*113*). Phenotypes were converted into normally distributed quantitative or a collection of binary (TRUE/FALSE) categorical variables. Full details of the phenotype pipeline are summarized here (*114*). We further excluded continuous phenotypes with fewer than one hundred samples and binary phenotypes with fewer than one hundred cases.

We generated polygenic risk scores for each of the generated traits with Bayesian regression and continuous shrinkage priors (*55*) using the significantly associated single-nucleotide polymorphisms. We ran a logistic or linear regression of the polygenic risk score on traits across all individuals, adjusting for the first 20 principal components of ancestry, and imputed sex.

### Genetic correlation of skeletal proportions with external phenotypes

We utilized cross-trait LD score regression for estimating genetic correlations between each of our skeletal proportions (*115*) and up to 700 additional quantitative and case-control phenotypes from the UKB that were precomputed by the Neale lab. Unlike polygenic risk score or phenotype association analysis, sample sizes for the case-control musculoskeletal disease traits were too low to assess genetic correlations between our skeletal proportion phenotypes and these disease traits in the UKB.

When examining other quantitative traits and applying a Bonferroni threshold correcting at the level of the number of skeletal proportion and biobank phenotype pairs, we saw well-known associations of skeletal proportions with puberty timing also previously associated with overall height. Here we were able to assess the impact of puberty timing on overall body proportions. While long bone proportions such as Femur:Height (r_g_ = 0.24, p = 1.77 × 10^-17^, r_g_ = 0.41, p = 1.71 × 10^-10^) and Humerus:Height were positively correlated with later onset of puberty overall body width measures such as Shoulder Width:Height were negatively correlated with age of puberty (r_g_ = -0.15, p = 2.50 × 10^-7^). We also saw that walking pace was increased by longer arms, and legs but decreased with torso length as a function of height.

We consistently found that increased Torso Length:Height was positively associated with body fat, BMI, and blood pressure (r_g_ = 0.09, p = 7.00 × 10^-4^; r_g_ = 0.16, p = 1.27 × 10^-8^; r_g_ = 0.15, p = 1.71 × 10^-6^). Overall, traits related to body mass were genetically correlated to several of our skeletal phenotypes such as Tibia:Height with left and right leg fat-free mass (r_g_ = 0.16, p = 4.90 × 10^-8^; r_g_ = 0.16, p = 2.36 × 10^-7^) or Humerus:Height with left and right arm fat-free mass (r_g_ = -0.14, p = 1.09 × 10^-6^; r_g_ = -0.14, p = 1.38 × 10^-6^), suggesting a possible link between skeletal body proportions and obesity. A full set of trait and skeletal proportion pairs of genetic correlation can be found in **Table S26.**

### Enrichment analysis for HARs

In order to investigate the evolution of body proportions in humans, we scanned for elevated levels of intersections between genes containing genome-wide significant SNPs and HARs through a modified version of the method outlined in Ke et al. (*56*). HARs are defined as regions of at least 100 base pairs (bp) which are conserved within the common ancestor of chimpanzees and humans but have increased rates of base pair substitutions in the human genome (*116–120*). For each phenotype, we first created annotations of protein coding regions that lie on our genome-wide significant SNPs using Ensembl’s GRCh37 Variant Effect Predictor version 103.4 (*121*). We selected the closest protein coding feature within 5,000 base pairs up- or downstream of the SNP. Using biotype categorizations identified by VEP, these protein coding features were: (“protein_coding”, “IG_C_gene”, “IG_D_gene”, “IG_J_gene”, “IG_LV_gene”, “IG_M_gene”, “IG_V_gene”, “IG_Z_gene”, “nonsense_mediated_decay”, “nontranslating_CDS”, “non_stop_decay”, “polymorphic_pseudogene”, “TR_C_gene”, “TR_D_gene”, “TR_J_gene”). We refer to the list of features for all independent genome-wide significant loci significantly associated with the trait as the *element set* for the phenotype being analyzed. Phenotypes with fewer than 50 elements in their set were removed from the analysis due to insufficient power. We then used BioMart (*110*) command line queries to generate the genomic locations (chromosome, start, stop) of each feature within the human genome. In order to scan for selection, we used BEDTools ‘intersect’ to compute the number of intersections found in the gene set with HARs sourced from literature.

To generate a background distribution of intersections per bp, we computed the HAR-element intersections per bp of 5,000 length-matched element sets. Because the distribution of these feature lengths is non-normal, we binned the element sets into deciles based on gene length and computed the average length *l* within each bin of size *n*. For each bin in the simulation, we sampled *n* random elements of length *l* to create our complete element set which was then used to compute the intersections per base pair of the simulated set. Due to the large differences in element set sizes and lengths across phhenotypes, a background distribution was generated independently for each phenotype analyzed (**Fig. S18**). On this background we fit Weibull distribution for computation of p-values of the observed intersections in comparison to the background. A comprehensive table of analyses performed can be found in **Table S23**.

### LDScore heritability enrichment in regions of evolutionary context

We applied stratified linkage disequilibrium score (S-LDSC) regression, which estimates whether a genomic region is enriched or depleted in heritability for a set of traits, capturing the contribution of variants in that genomic region towards trait variation, and whether this contribution is more or less than expected given the relative proportion of variants in that region. We used the following genomic annotations marking different evolutionary periods: (A) epigenetic elements that gained novel function in the fetal brain since our divergence with rhesus macaque (H3K27as and H3K4me2 histone modification peaks in the fetal cerebral cortex gained in humans compared to mouse and rhesus macaque) at different developmental stages or post-conception weeks (PCW) (Fetal human-gained (HG) enhancers and promoters at 7 PCW, 8.5 PCW, and 12 PCW) (*60*), (B) epigenetic elements that gained novel function in the adult brain since our divergence with rhesus macaque (Adult human-gained (HG) enhancers and promoters) (*61*), (C) ancient selective sweeps from the extended lineage sorting method capturing human-specific sweeps relative to Neanderthal/Denisovan (*63*), (D) regions depleted in Neanderthal ancestry (*122, 123*) (E) regions depleted in Neanderthal and Denisovan ancestry (*123*), and (F) putatively introgressed variants from Neanderthals (*62*). We did not include HAR annotations as part of this analysis as these annotations were small and the use of such annotations in this context might not always control type 1 error (*124*).

Using S-LDSC for our skeletal traits, we analyzed our test annotations in a model simultaneously incorporating several other regulatory elements, measures of selective constraint, and linkage statistics (baselineLDv2.2 with 97 annotations) (*59, 64–66*) to estimate heritability enrichment while minimizing bias due to model misspecification.

## Supplementary Tables

**Table S1** - GWAS population summary

This table contains summary data on the population subset used in our GWAS from the UKB

**Table S2** - Architecture comparison

This table contains data comparing the performance of HRNet against ResNet on our landmark estimation task

**Table S3** - IDPs

This table contains a list of all generated IDPs

**Table S4** - HPE measurement error

This table contains various error metrics comparing human-derived measurements of bone and body lengths to HRNet-derived measurements

**Table S5** - 864 Model landmark results

This table contains a comparison of landmark estimations between human annotated images and HRNet prediction for 864×288 images

**Table S6** - 960 Model landmark results

This table contains a comparison of landmark estimations between human annotated images and HRNet prediction for 960×384 images

**Table S7** - Adjusted IDPs to Height

This table contains correlations between measurements as ratios of height and height itself

**Table S8** - Sex difference t-tests

This table contains the results from T-test comparisons between sex for each IDP

**Table S9** - Age phenotype correlations

This table contains the results from linear regression analyses between age and each IDP, also separated by sex

**Table S10** - GWAS QC

This table contains data showing how many individuals in the UKB were removed from our final GWAS population for each QC step

**Table S11** - GCTA GWAS heritability

This table contains the heritability for each IDP as determined by GCTA

**Table S12** - LDSC GWAS heritability and lambdas

This table contains the heritability and lambda values for each IDP as determined by LDSC

**Table S13** - Phenotype and genetic correlations

This table contains correlations, standard errors, and p-values of genotype and phenotype correlations for IDPs included in Figure 2B

**Table S14** - MAGMA GO terms GSA

This table contains output from MAGMA GSA for each phenotype as well as gene set

**Table S15** - OMIM gene set

This table contains all genes from OMIM database search “Skeletal Growth Abnormality” that were used in a GSA

**Table S16** - OMIM skeletal GSA

This table contains output from a GSA over the OMIM gene set for each phenotype

**Table S17** - Clumped SNPs

This table contains output from PLINK --clump ranges command including lead SNP, p-value, and number of kilobases in each clump

**Table S18** - Gene ranges

This table contains data regarding gene mapping for each clump range as well as whether the single clump range genes are related to known mouse phenotypes and rare human disease

**Table S19** - TWAS results

This table contains Bonferroni corrected TWAS output

**Table S20** - Musculoskeletal regressions

This table contains output from logistic regression analyses for each skeletal proportion and musculoskeletal disease or area of pain

**Table S21** - PRS analysis

This table contains output from PRS analyses for each skeletal proportion and musculoskeletal disease or area of pain

**Table S22** - Multivariate genetic architecture of skeletal endophenotypes table 1

This table contains statistics of correspondence between direct and model-implied associations across musculoskeletal diseases and skeletal endophenotypes.

**Table S23** - HAR analysis results

This table contains information regarding source and publication of GWAS summary statistics as well as output of enrichment overlap analysis

**Table S24** - LDSC heritability enrichment meta-analysis

This table contains output from S-LDSC heritability enrichment meta-analysis

**Table S25** - LDSC heritability enrichment analysis

This table contains output from S-LDSC heritability enrichment analysis for all traits and annotations

**Table S26** - Genetic correlations UKB

This table contains output from genetic correlations between skeletal proportions and all other traits in the UKB

**Table S27** - ICD10 Codes

This table contains all ICD10 codes used in our analyses

**Table S28** - UKB phenotypes FID

This table contains the FID of each UKB traits used in our analyses

**Table S29** - Initial deep learning QC

This table contains the number of patients removed from each QC step before landmark estimation

**Table S30** - Image pixel data

This table contains the number of full body skeletal DXA images for each pixel aspect ratio in the UKB

**Table S31** - Annotation reproducibility

This table shows annotator accuracy for each landmark on 10 duplicate images

**Table S32** - Outlier image removal

This table shows the number of patients removed due to outlier values following image measurement

**Table S33** - Genetic correlations ratios residuals covariates

This table shows genetic correlation values between skeletal measurements as a ratio of height, skeletal measurements with height as a covariate, and residuals of skeletal measurements regressed against height

**Table S34** - Residual correlations

This table shows genotype and phenotype correlations between skeletal measurements with height as a covariate, mimicking Table S13

**Table S35** - Sensitivity analysis for height adjustment

This table contains summary statistic data from skeletal measurements alone, skeletal measurements as a ratio of height, skeletal measurements with height as a covariate, and height

**Table S36** - Multivariate genetic architecture of skeletal endophenotypes table 2

This table contains the summary statistics of skeletal traits and musculoskeletal diseases and results from LD Score Regression (LDSC) in all autosomes.

**Table S37** - Multivariate genetic architecture of skeletal endophenotypes table 3

This table contains standardized factor loadings from two-factor EFA solution of skeletal traits within odd-numbered autosomes.

**Table S38** - Multivariate genetic architecture of skeletal endophenotypes table 4

This table contains goodness-of-fit indices for confirmatory factor models of skeletal endophenotypes on even autosomes.

**Table S39** - Multivariate genetic architecture of skeletal endophenotypes supplementary table 5

This table contains the results of applying the genetic bifactor model of skeletal endophenotypes on even autosomes (Model C)

**Table S40** - Multivariate genetic architecture of skeletal endophenotypes supplementary table 6

This table contains the results of applying the genetic bifactor model of skeletal endophenotypes on even autosomes (Model D)

**Table S41** - Multivariate genetic architecture of skeletal endophenotypes table 7

This table contains the results of applying the genetic bifactor model of skeletal endophenotypes on all autosomes (Model D)

**Table S42** - Multivariate genetic architecture of skeletal endophenotypes supplementary table 8

This table contains results showing observed effects of skeletal endophenotypes on musculoskeletal (MSK) diseases and common factor estimates as estimated from Model D.

**Table S43** - Height independent SNPs

This table contains the p-value of independent SNPs from our skeletal elements as ratios of height and TFA as well as the p-value of each SNP in a GWAS for height

**Table S44** - Transcriptome analysis

This table contains the results of MAGMA GPA for skeletal proportions and various gene expression data including expression from various bone layers, different time points, and different types of bones

**Table S45** - Musculoskeletal covariates

This table contains the list of covariates used in our logistic regression analyses and the FID from the UKB

## Acknowledgements

This research has been conducted using the UKB Resource under Application Number 65439. We thank Carrie Zhu and Arbel Harpak for insightful discussions and comments on sex-specific analysis. We thank Phillip Wooley and Muyoung Lee for early implementations of our deep learning models.

*Funding:* V.M.N was supported on a grant from the Allen Discovery Center program, a Paul G. Allen Frontiers Group advised program of the Paul G. Allen Family Foundation and a Good Systems for Ethical AI grant from the University of Texas at Austin. O.S. and B.F. were supported on an NSF Graduate Research Fellowship DGE 2137420 and DGE 2137420. E.M.J. and B.F were supported by an NIH T32 grant 5T32LMO012414. B.F. was also supported on a UT Austin Provost’s Graduate Excellence Fellowship. E.M.T.D. and J.F. were supported by NIH grants R01MH120219, R01AG054628 and RF1AG073593. Additionally, E.M.T.D and J.F. are members of the University of Texas Center on Aging and Population Sciences and the University of Texas Population Research Center, which are supported by NIH grants P30AG066614 and P2CHD042849, respectively.

*Author contributions:* E.K, T.S. and V. M. N. wrote the paper with input from all co-authors. E.K., E.M.J., O.S, F.G., B.F, Z.T. K.V., J.F., E.M.T.D., M.S., P.J., T.S. and V.M.N. performed analysis.

*Competing interests:* The authors declare no competing interests.

*Data and materials availability:* Code used for performing the deep learning based key point identification and QC of the DXA data can be found at https://github.com/EucharistKun/Human-Skeletal-Form/, https://github.com/briannaflynn/dxaconv/. Code for the HAR analysis can be found here: https://github.com/ossmith/HARE/. GWAS Sumstats are available here: https://utexas.box.com/s/vli4rb4ise7qbdx5gmgpakga5n9ce2lr. Individual level information of skeletal lengths has been reported back to the UKB and will be available upon publication.

---

**Fig. S1.**
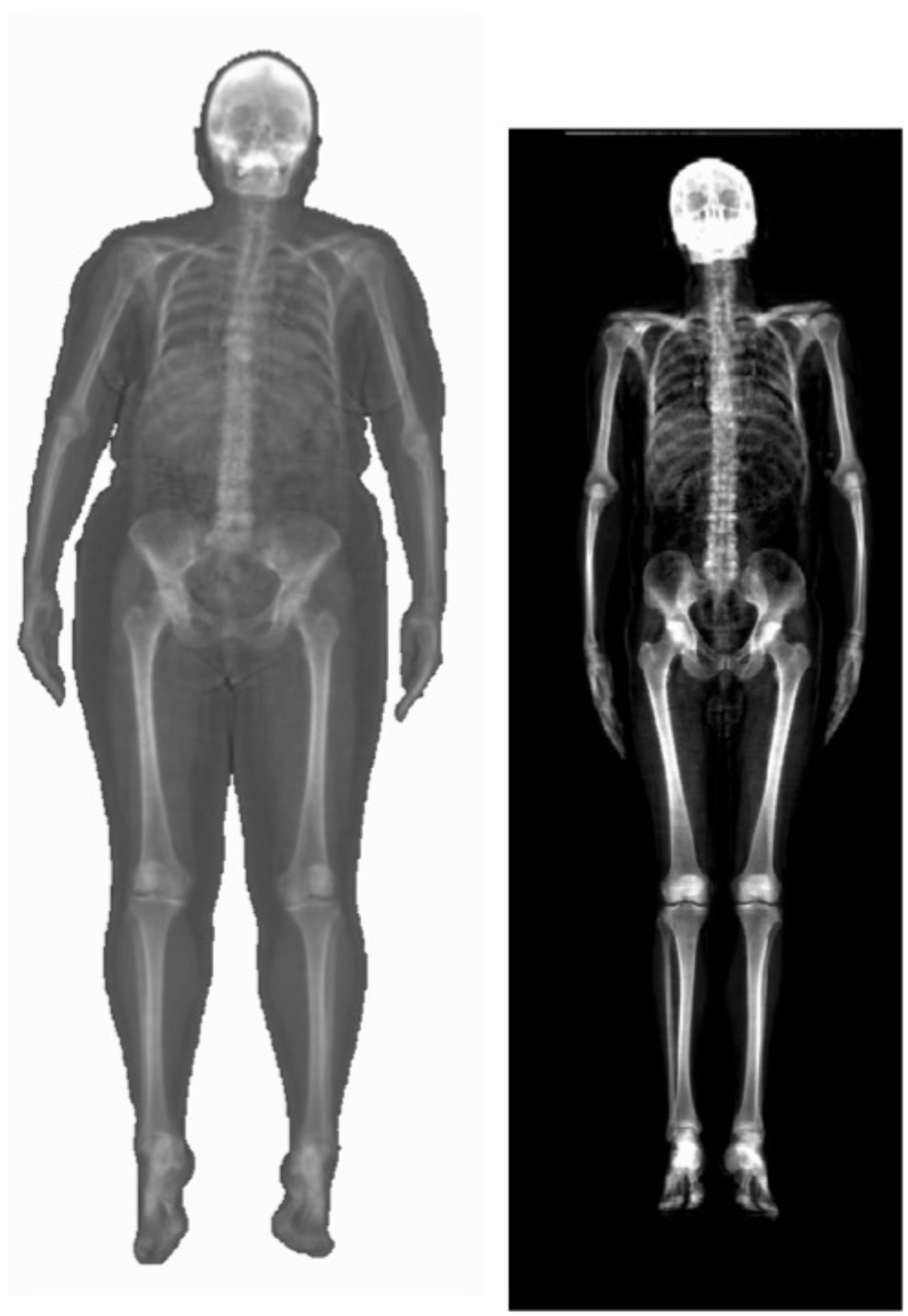
Types of DXA images acquired from the UKB. (Left) Image of patient imaged on white background. (Right) Image of patient imaged on black background. Sizes of images are true to scale.

**Fig. S2.**
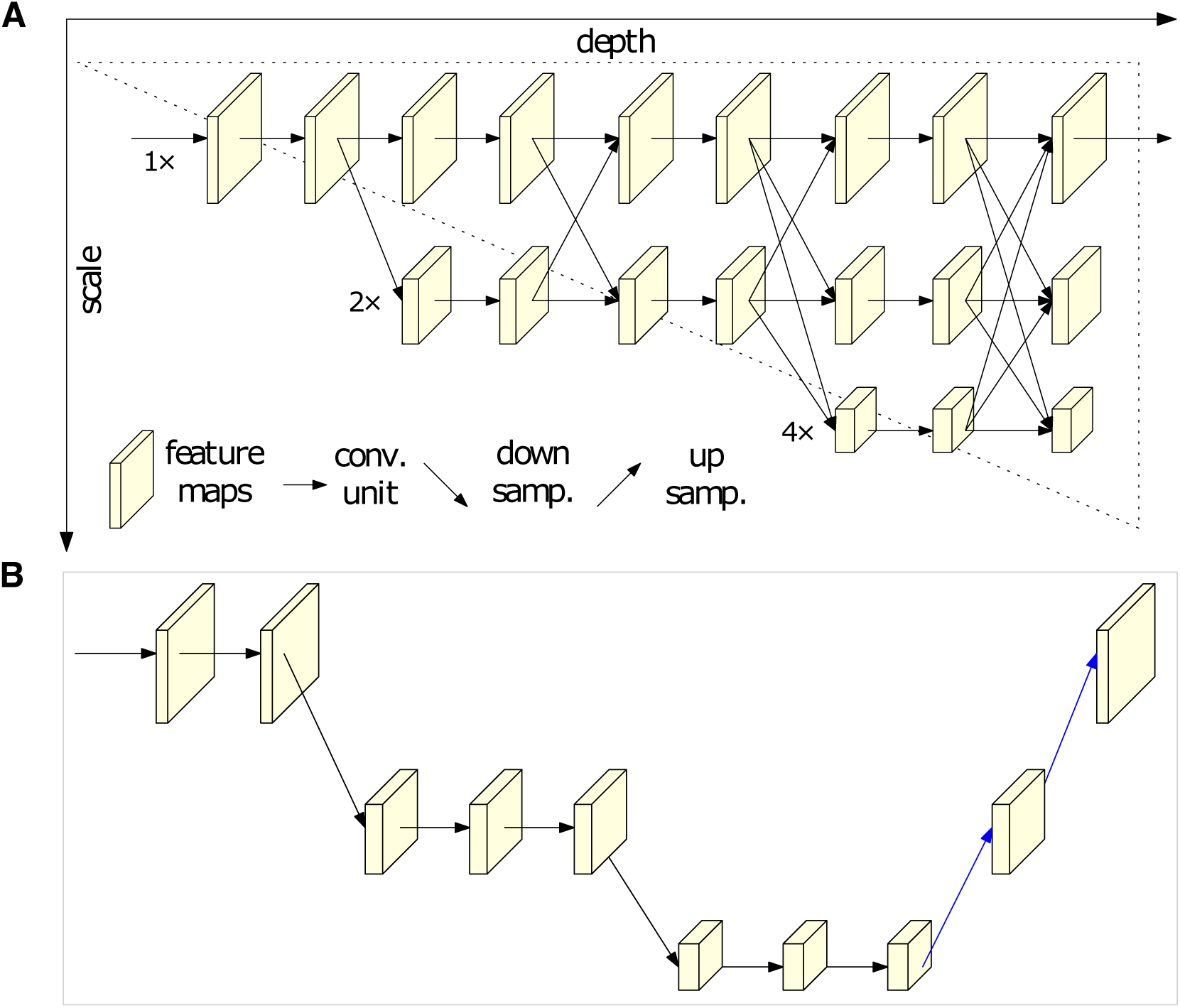
A comparison of HRNet and ResNet deep learning architectures. (**A)** High-Resolution Network (HRNet) architecture maintains parallel high to low resolution subnetworks. (**B)** Simple Baseline deep learning architecture (ResNet) which relies on a high-to-low and low-to-high framework. Both images are taken directly from Sun et al., (*35*) to illustrate the architectural differences between HRNet and a standard architecture for this prediction task.

**Fig. S3.**
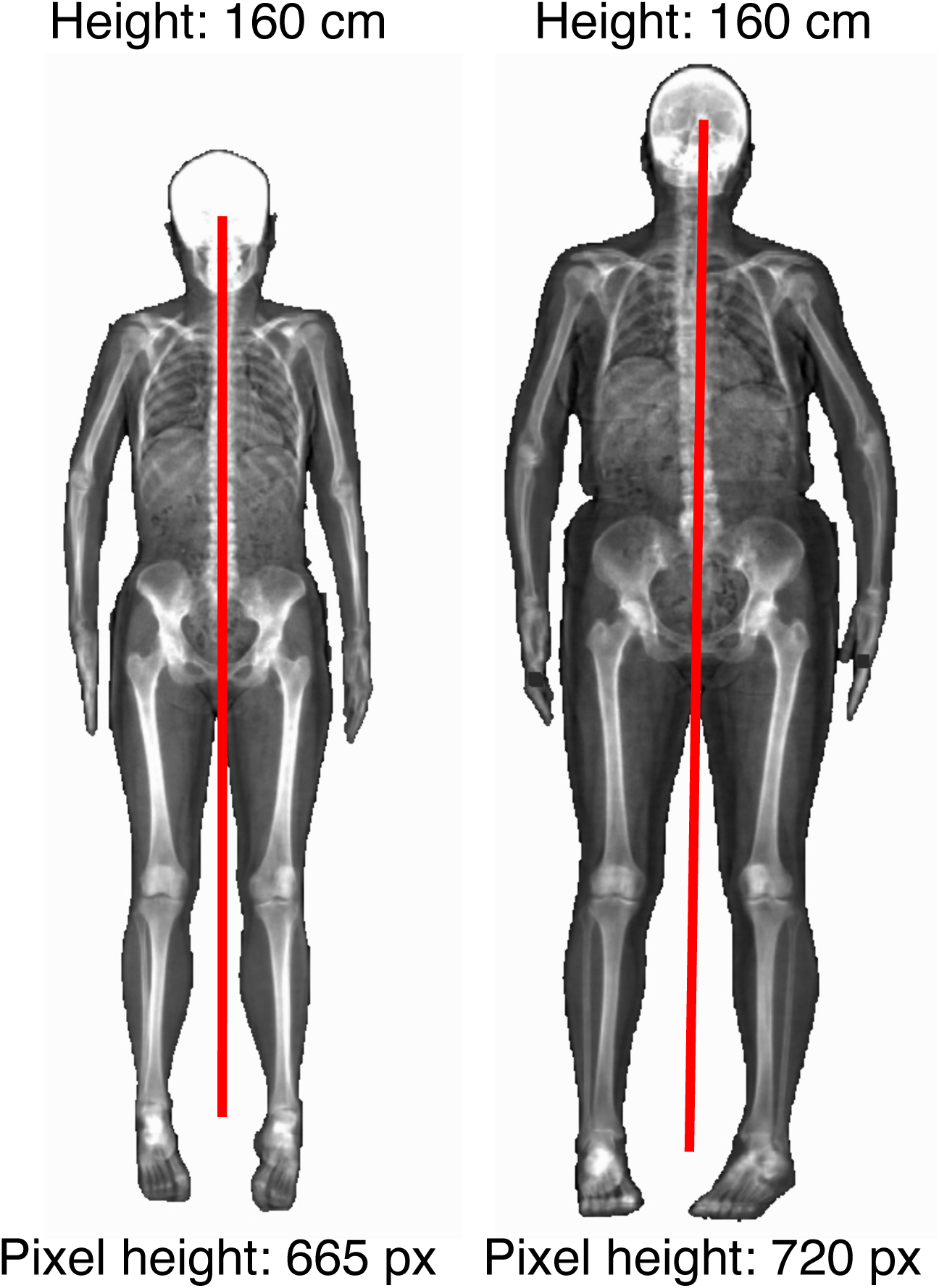
DXA images from the UKB that have undergone different image scaling. Example of two individuals who were measured to be the same height in the FID 50 in the UKB (overall height) but pixel-based measurements of one image were considerably smaller than the other due to image scaling/resolution differences.

**Fig. S4.**
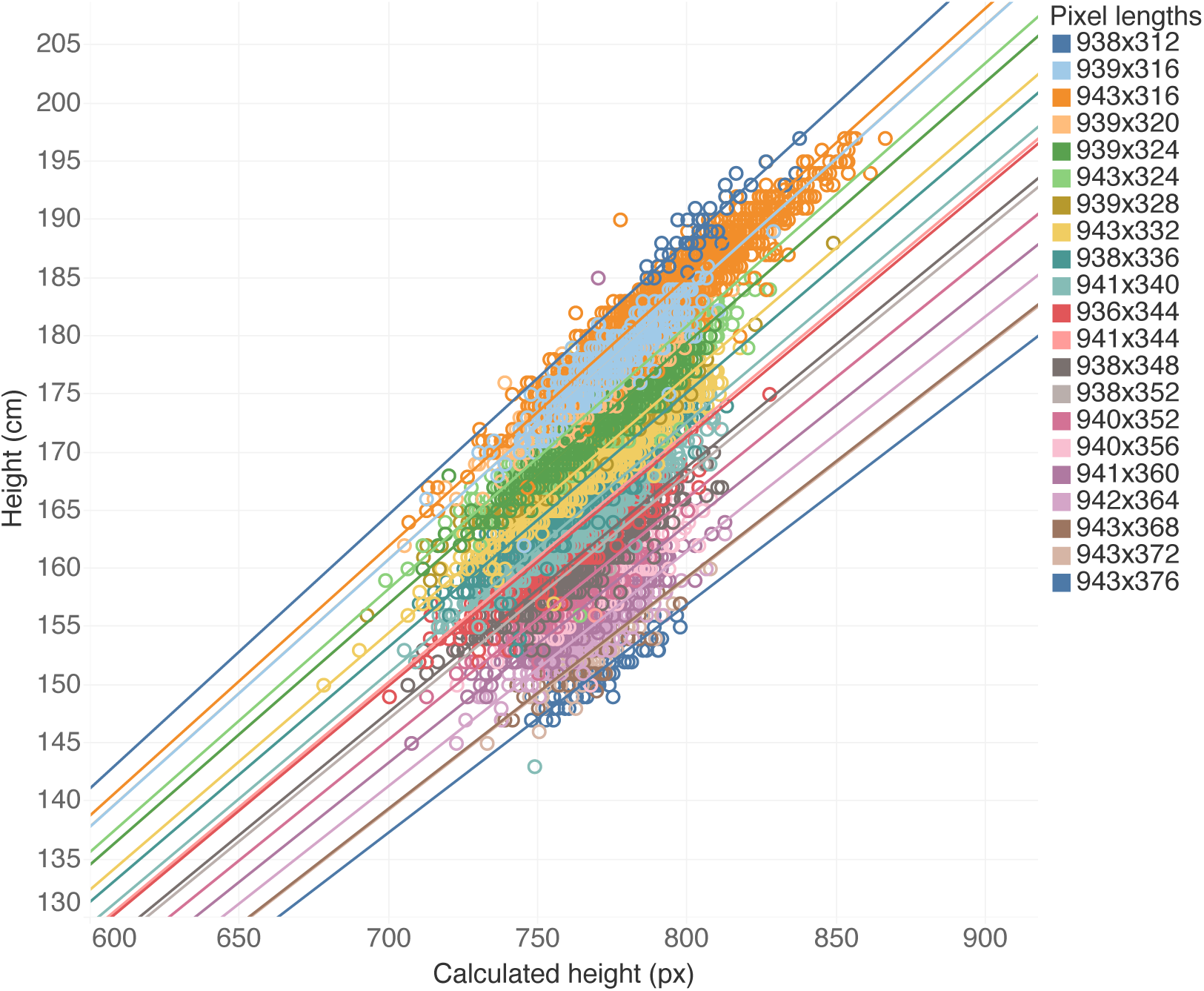
A linear regression of image-measured height against UKB-measured height. For each image pixel-ratio, we regressed height measured in the UKB with height we calculated in pixels from the DXA scan. This provided a conversion from pixels to cm that we used as a normalization factor to correct for differences in resolution.

**Fig. S5.**
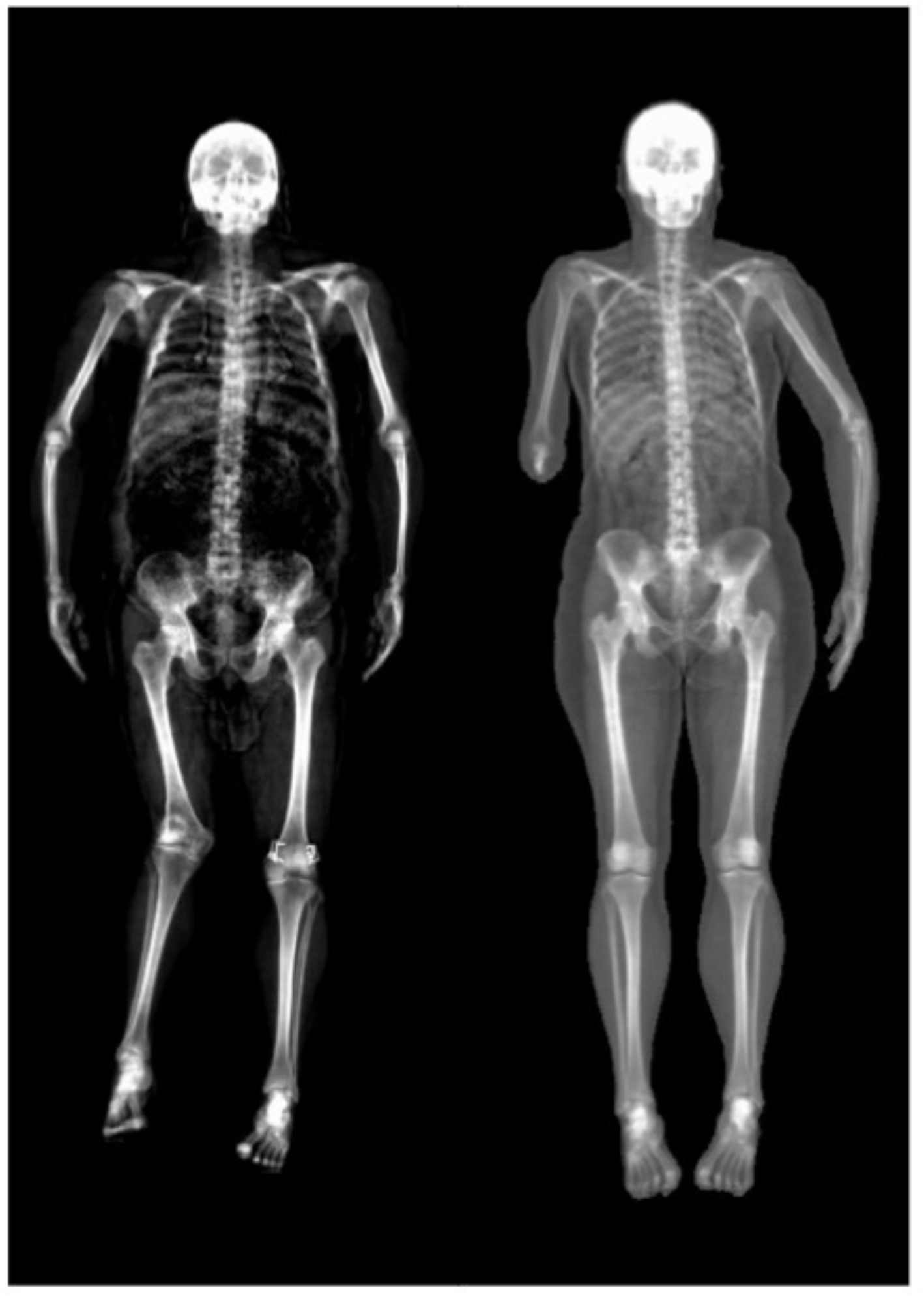
Examples of individuals who were outliers on our measurement and were removed from analysis. (Left) Individual with femur deformity and metal implants. (Right) Individual with missing forearm.

**Fig. S6.**
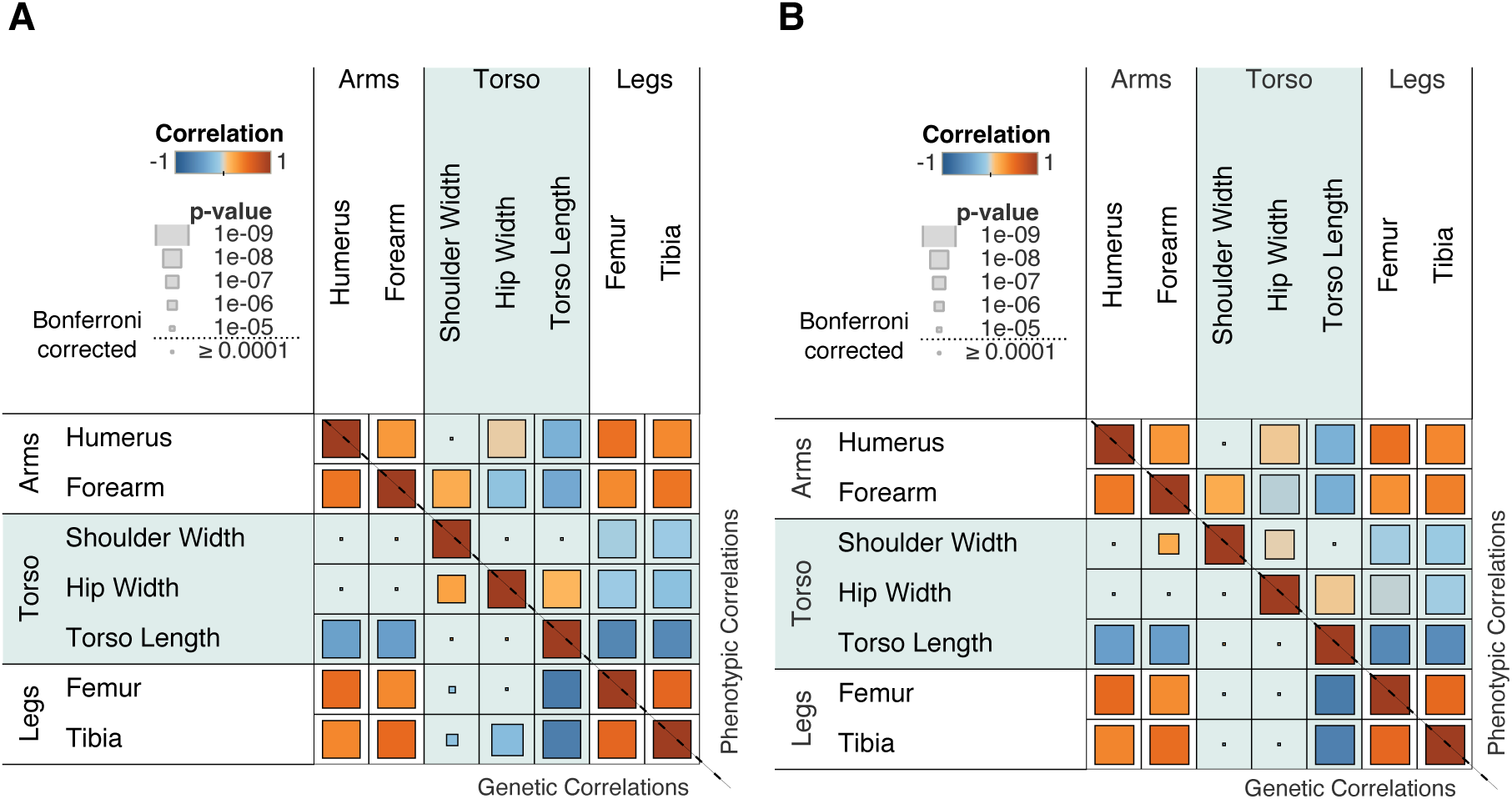
A heatmap comparison of genotype and phenotype correlations between ratios and residuals. (A) Matrix of genotype and phenotype correlation with each phenotype computed as a ratio of height. (**B)** Matrix of genotype and phenotype correlation with each phenotype computed by regressing the phenotype with height and then obtaining residuals.

**Fig. S7.**
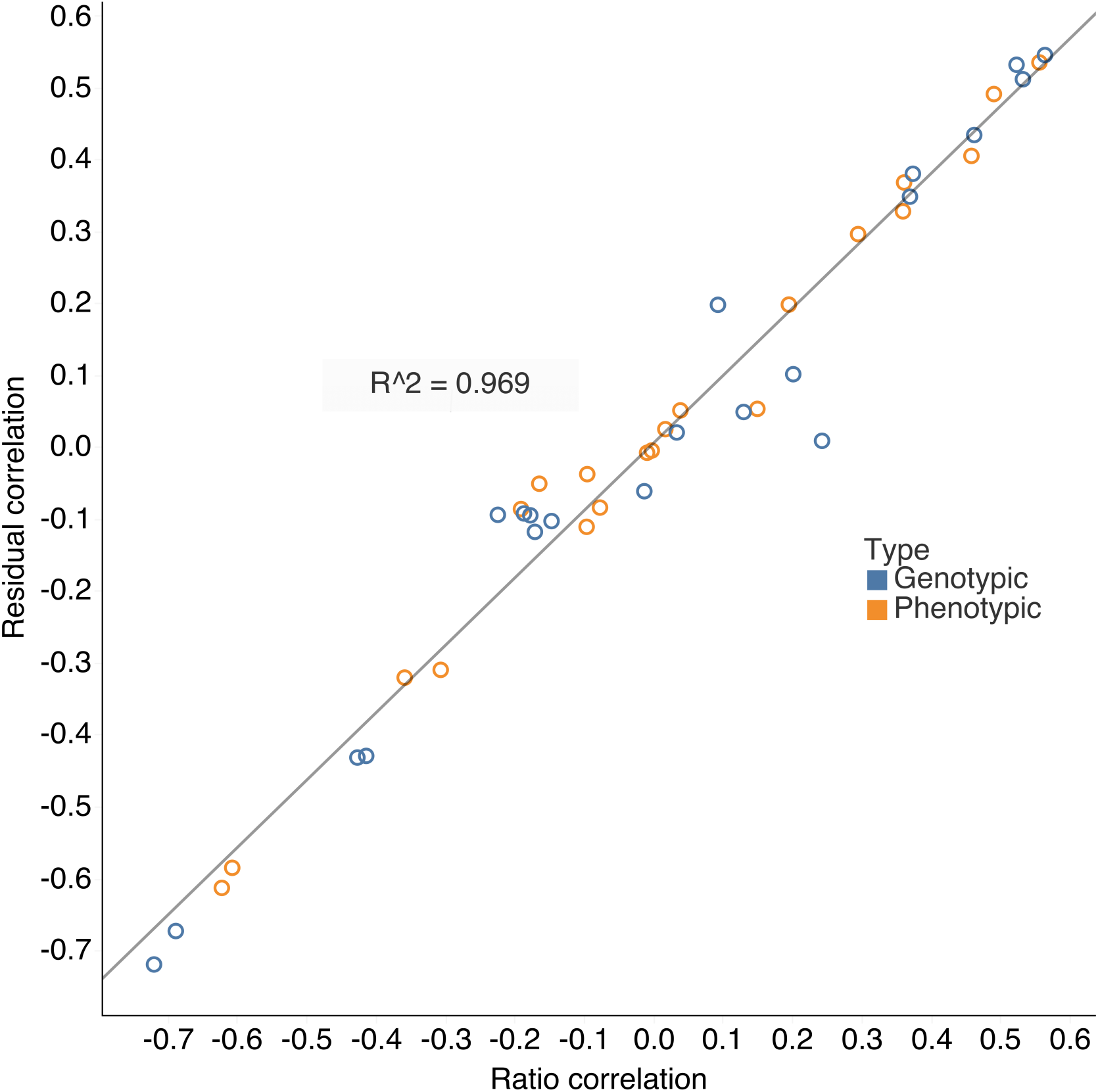
Correlation of genotype and phenotype correlations across skeletal traits, computed using ratios with height and second residualizing for height

**Fig. S8.**
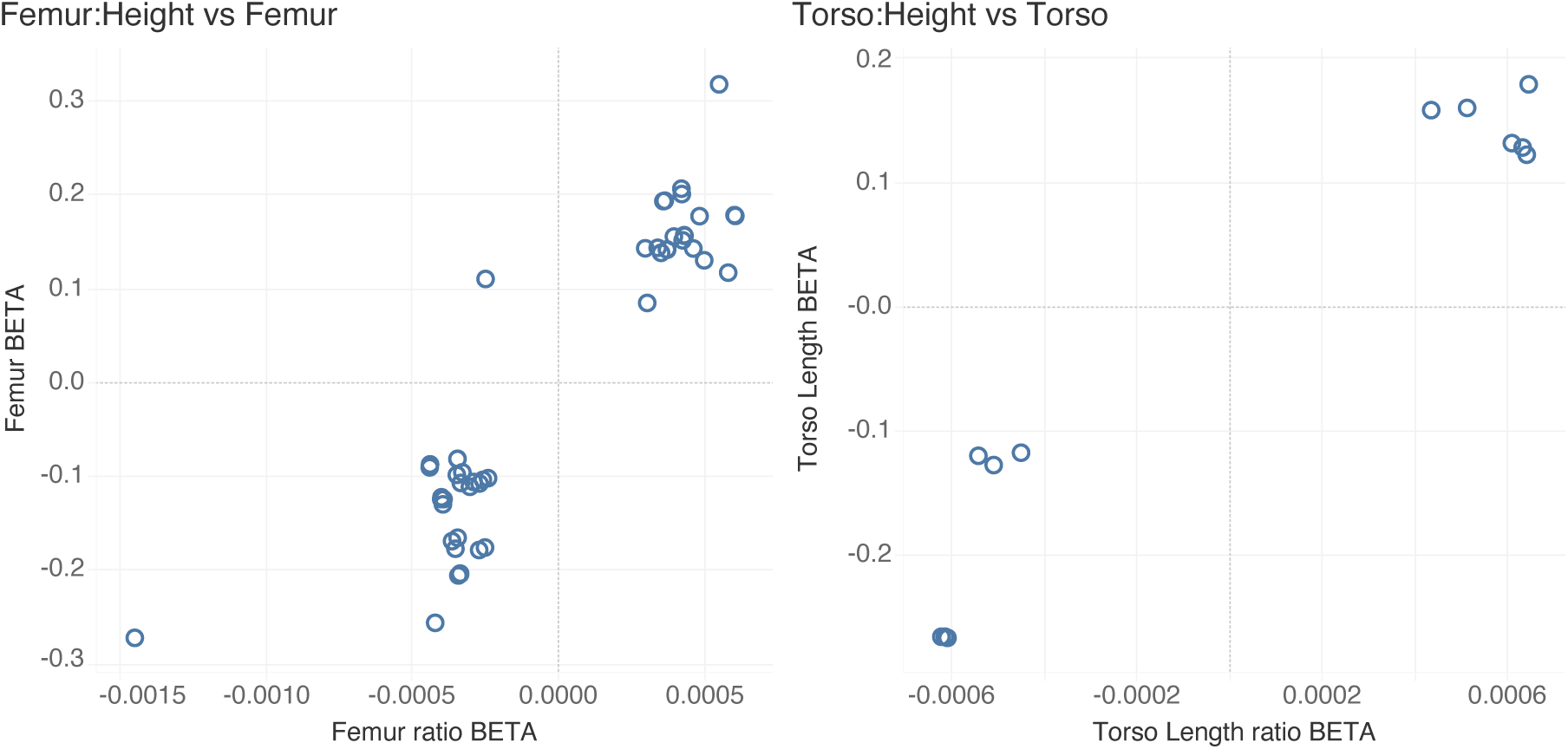
Comparison of effect estimates of independent genome-wide significant SNPs across different phenotypes. Effect estimates of genome-wide significant SNPs for each phenotype (p < 5e-08) showing same effect directionality for skeletal proportions and raw measurements.

**Fig. S9.**
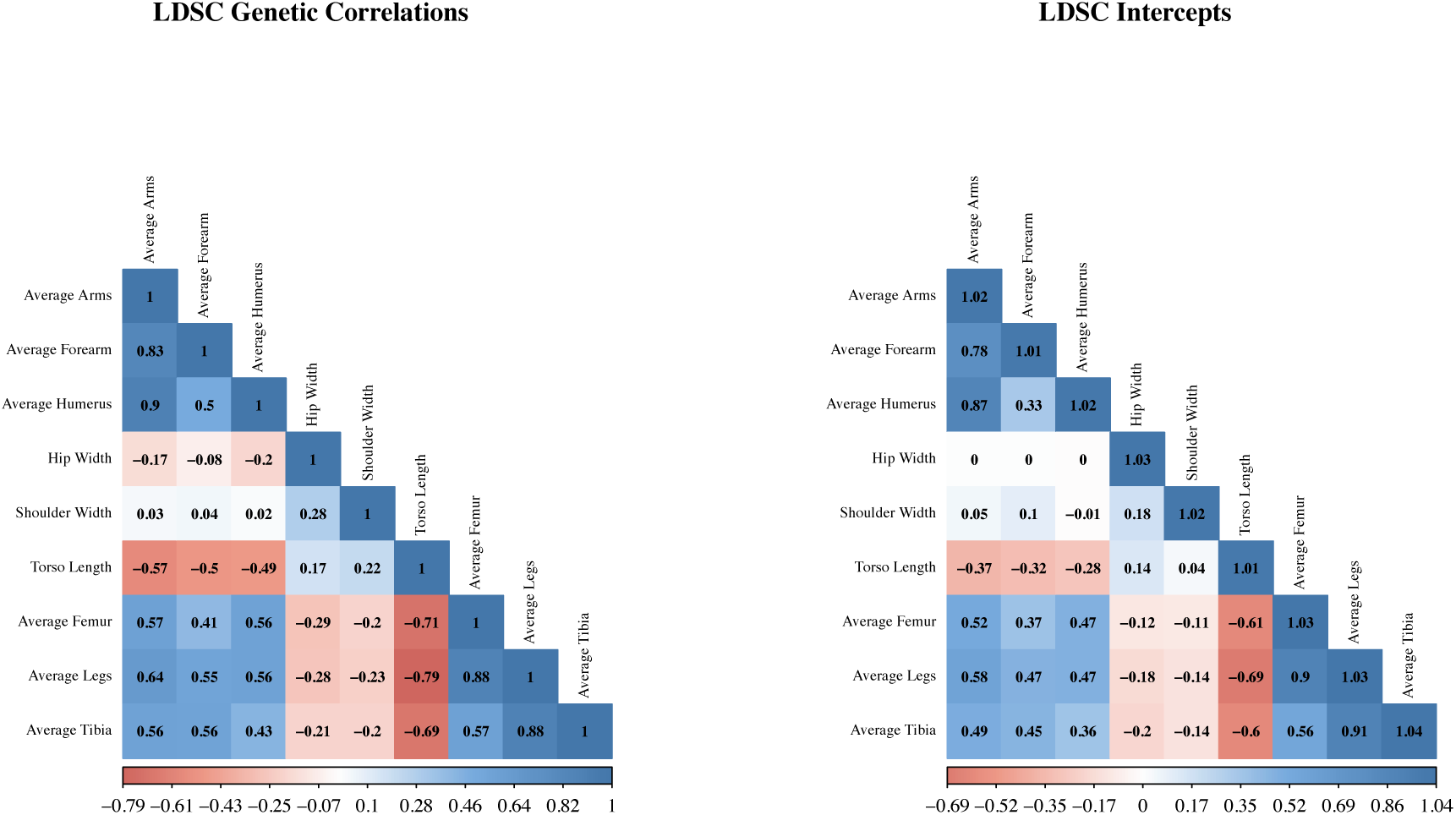
Heatmap of genetic correlations and LDSC cross-trait intercepts across skeletal proportion phenotypes within odd-numbered chromosomes

**Fig. S10.**
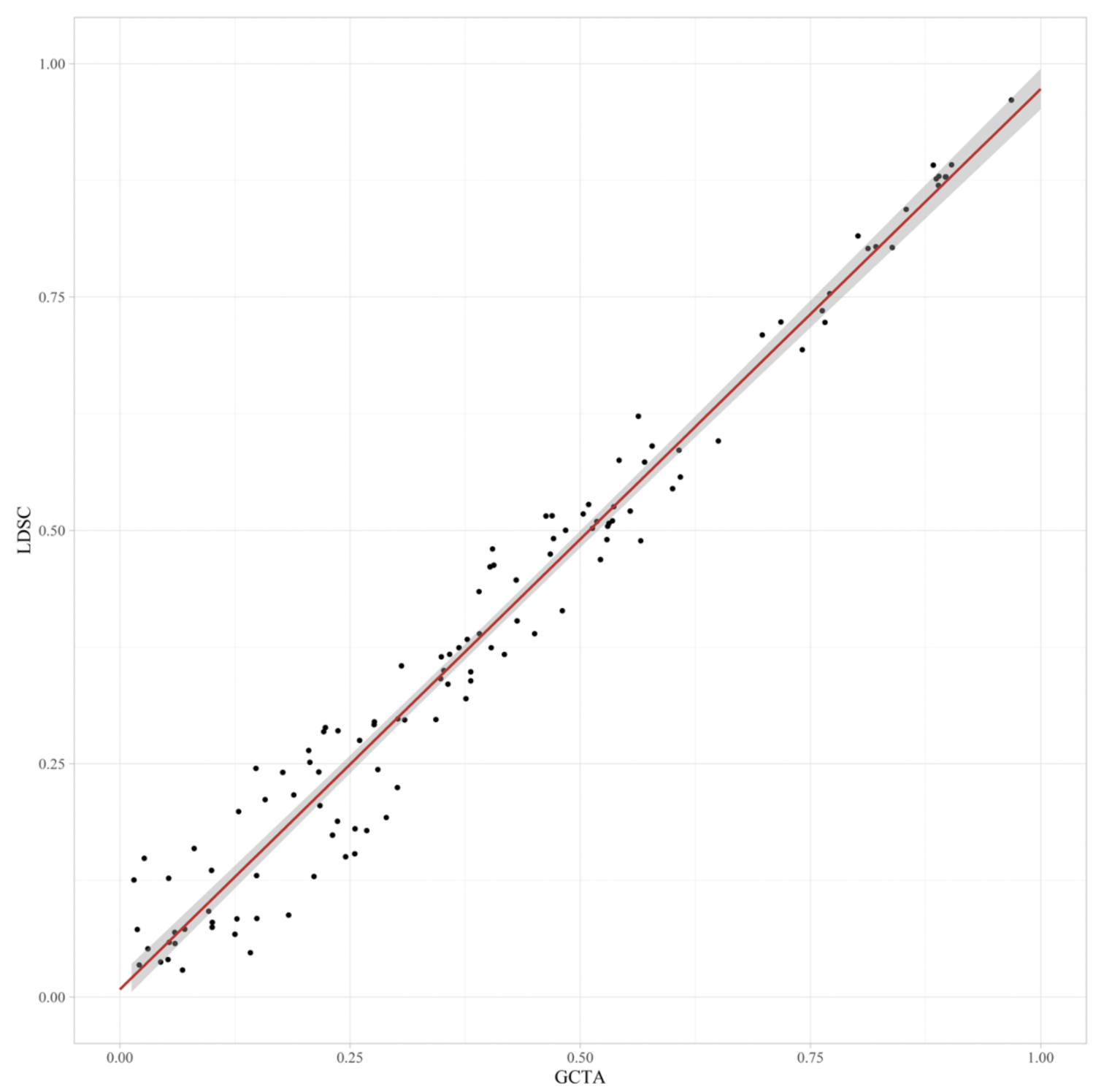
Scatterplot of GCTA and LDSC genetic correlation estimates across skeletal ratios.

**Fig. S11.**
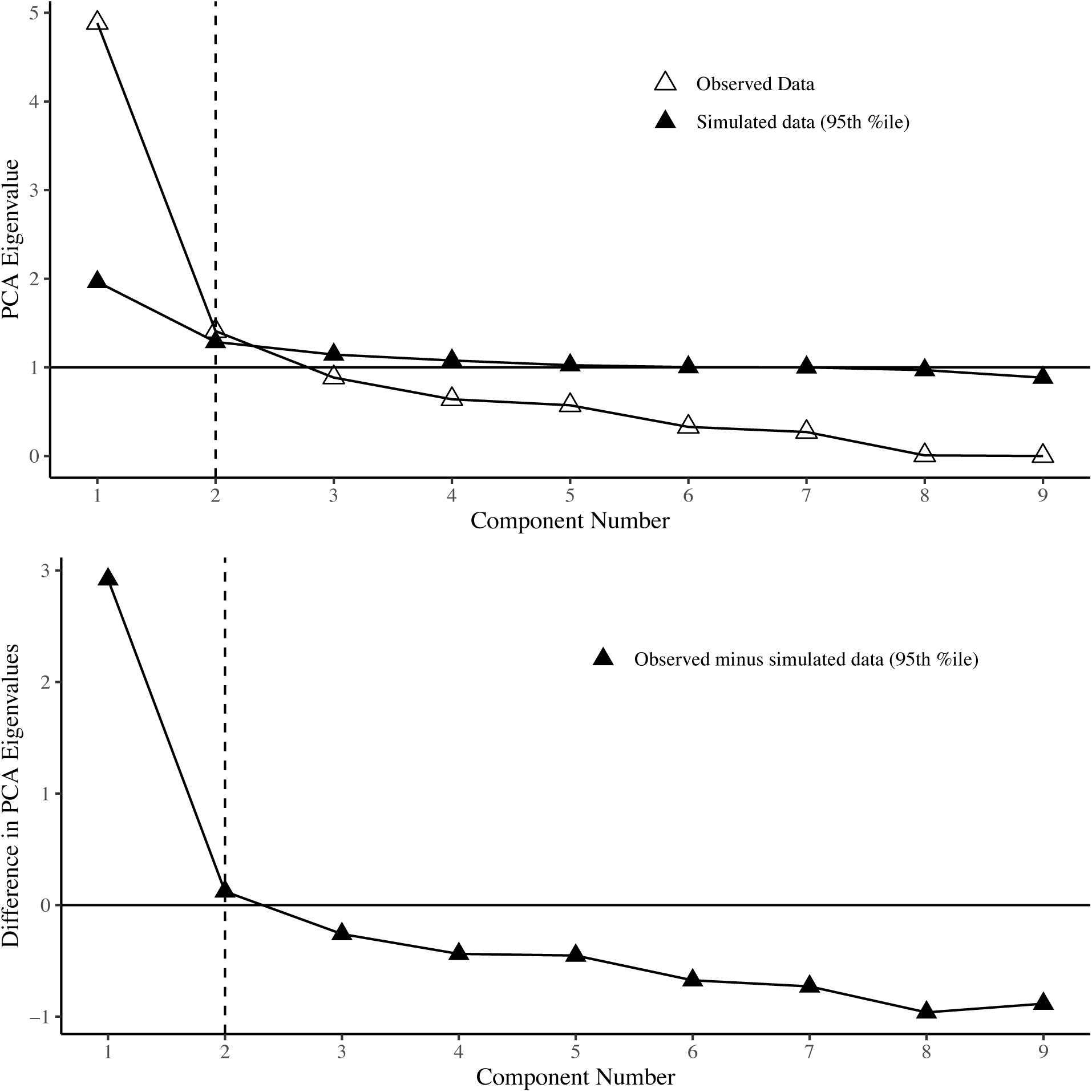
Screeplots of PCA and difference in PCA from LDSC Parallel Analysis

**Fig. S12.**
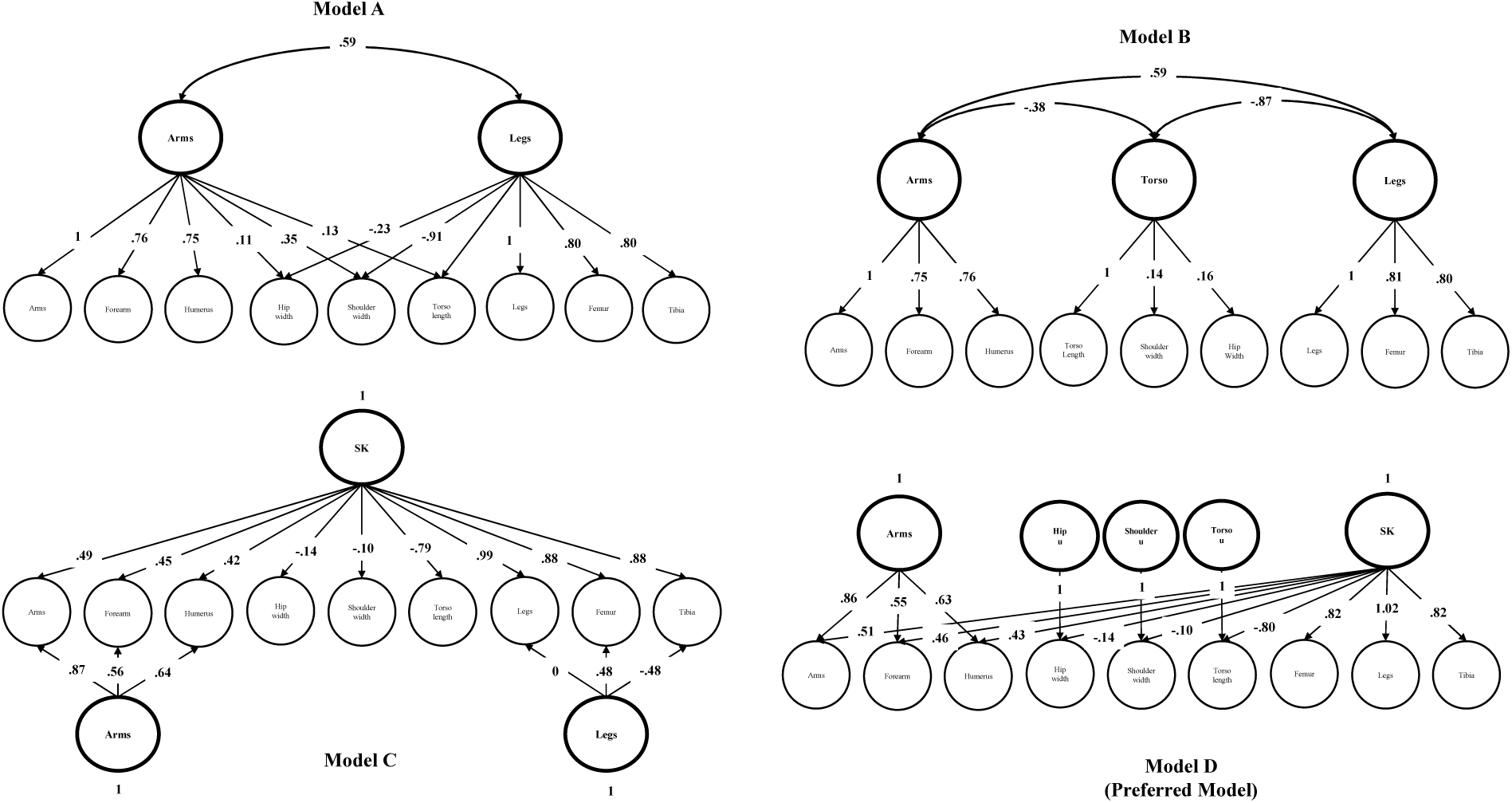
Confirmatory factor models of skeletal traits (even autosomes)

**Fig. S13.**
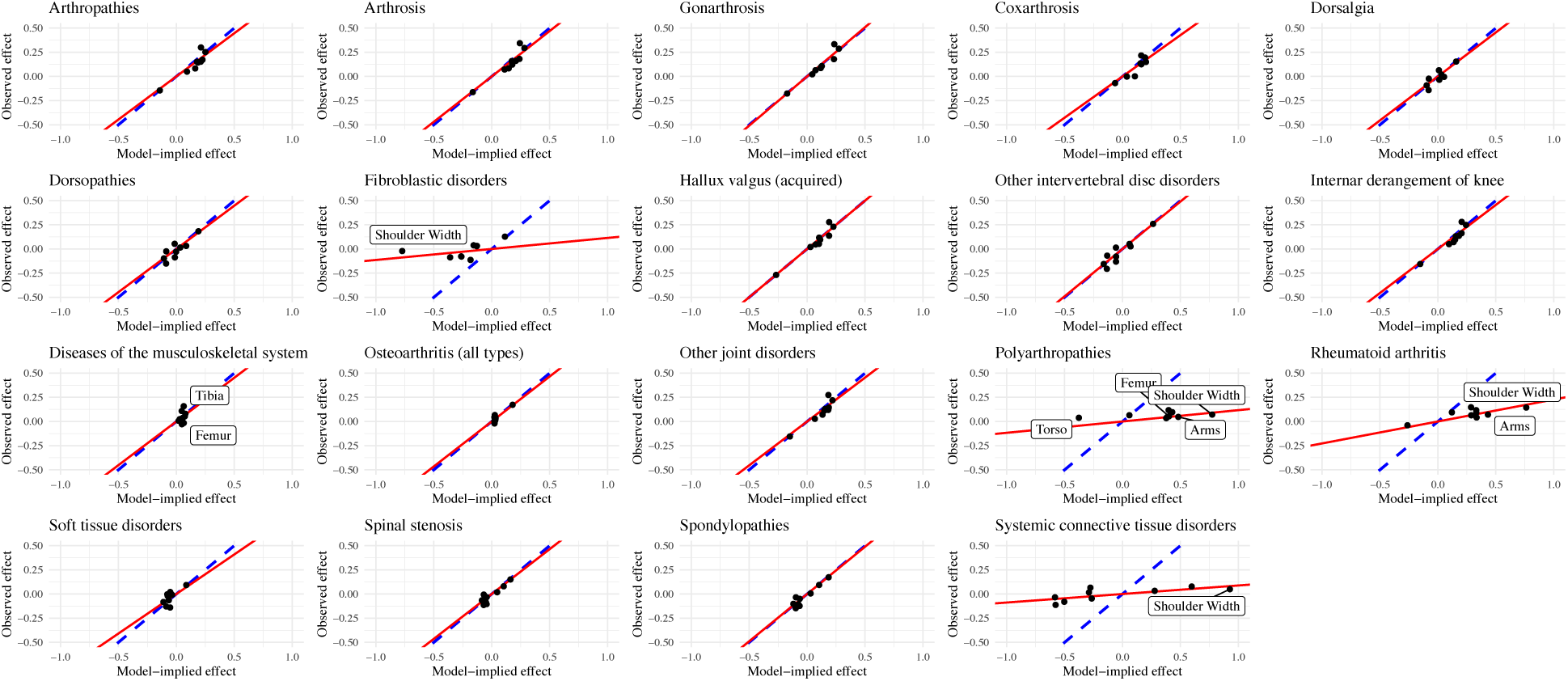
Scatterplots of observed and model-implied effects between musculoskeletal diseases and skeletal endophenotypes. Red lines represent best fitting regression lines. Blue dashed lines represent perfect fit (Observed effects = Model-implied effects). Labeled traits are outliers detected based on standardized differences between the observed and the common factor model-implied effects for the skeletal traits > 2.

**Fig. S14.**
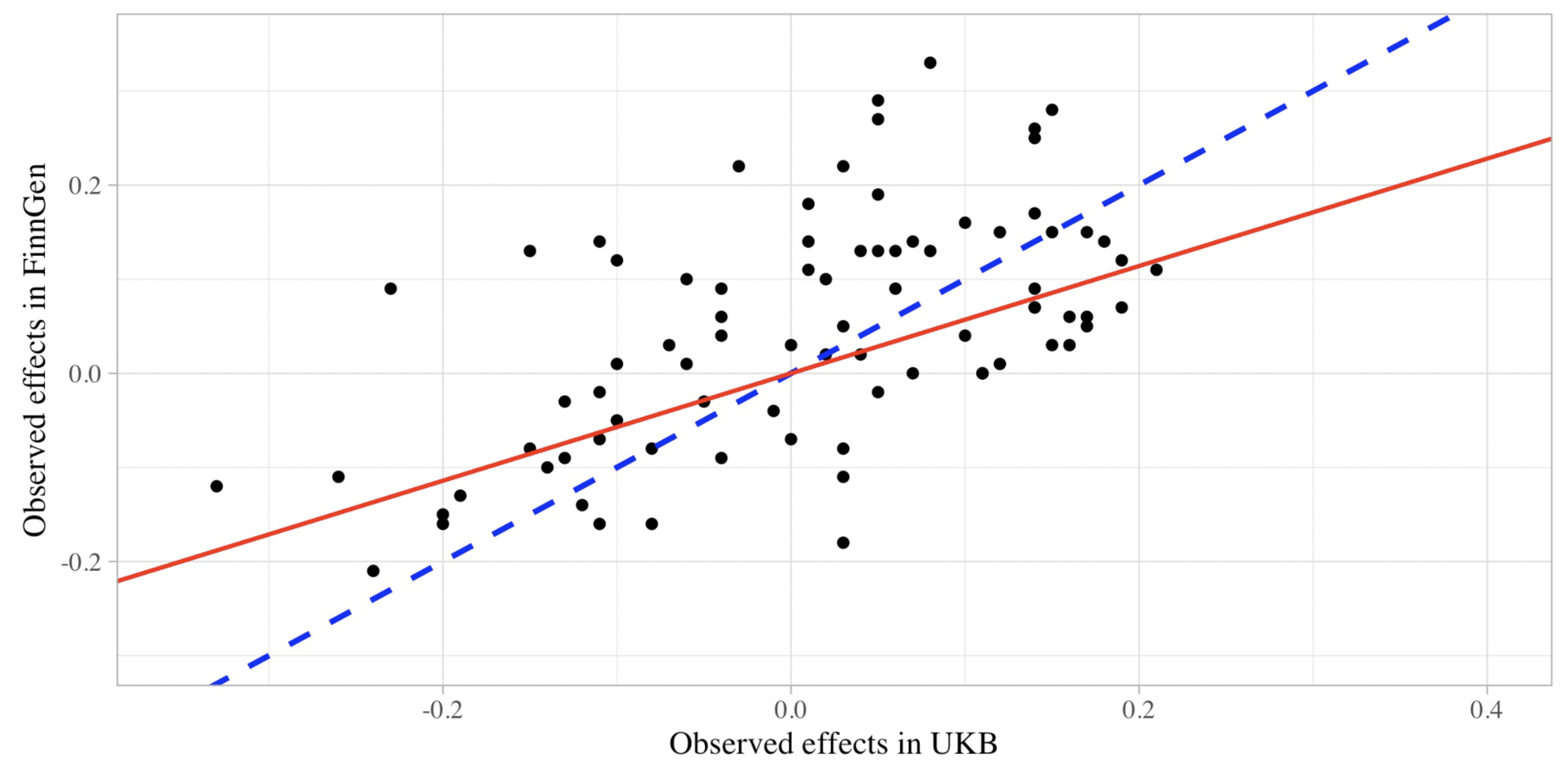
Observed effects of skeletal traits on musculoskeletal diseases common in UKB and FinnGen. Red lines represent best fitting regression lines. Blue dashed lines represent perfect correspondence (intercept = 0, slope = 1).

**Fig. S15.**
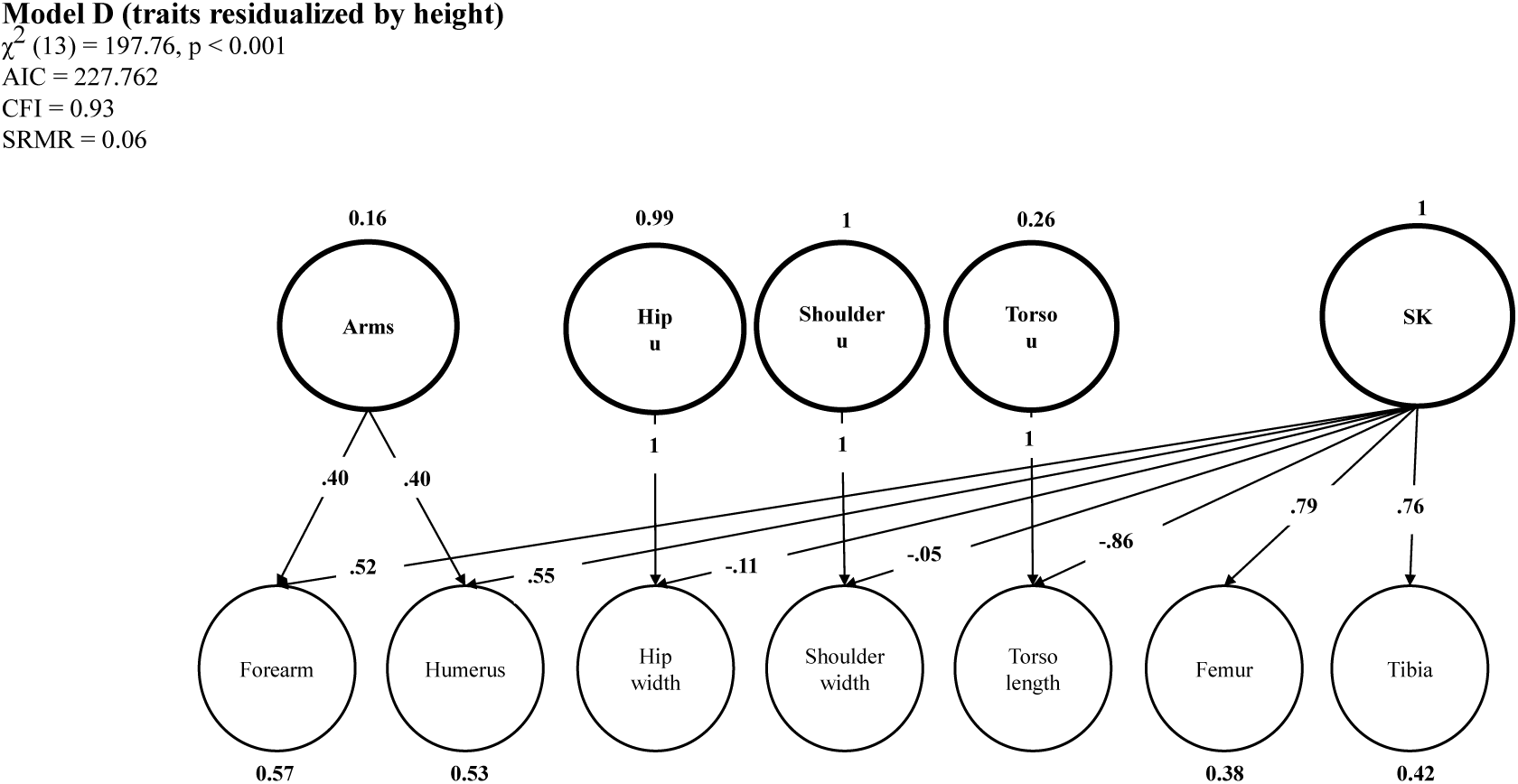
Confirmatory Factor Model D applied to residuals. Preferred model D fully standardized parameter estimates fitted on height-residualized skeletal traits as well as excluding overall arms and leg length

**Fig. S16.**
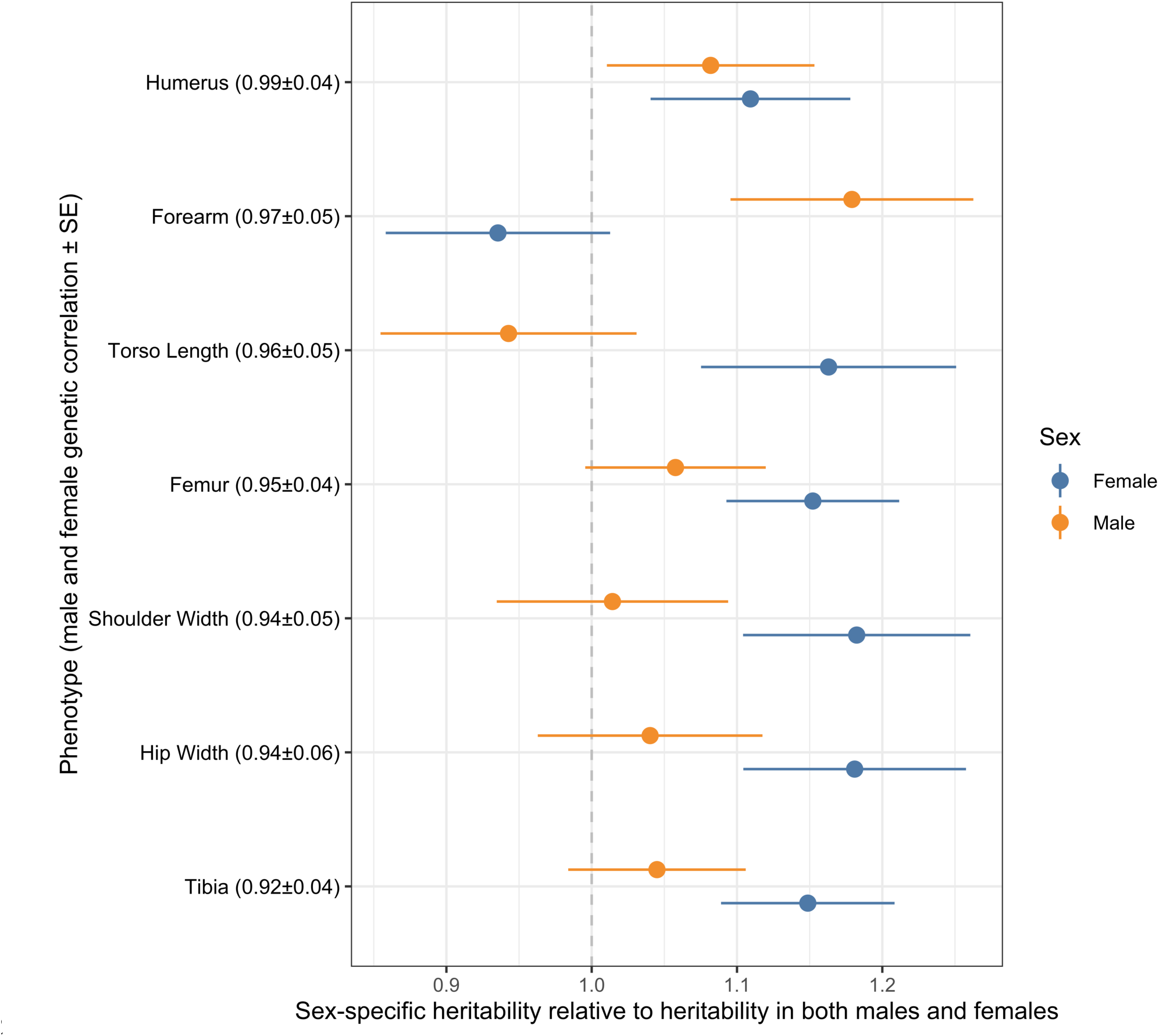
Genetic correlations between males and females, estimated using bi-variate LD Score Regression shown for each trait (y-axis). SNP heritability divided by the SNP heritability estimated in the sample with both sexes combined (x-axis) for all traits

**Fig. S17.**
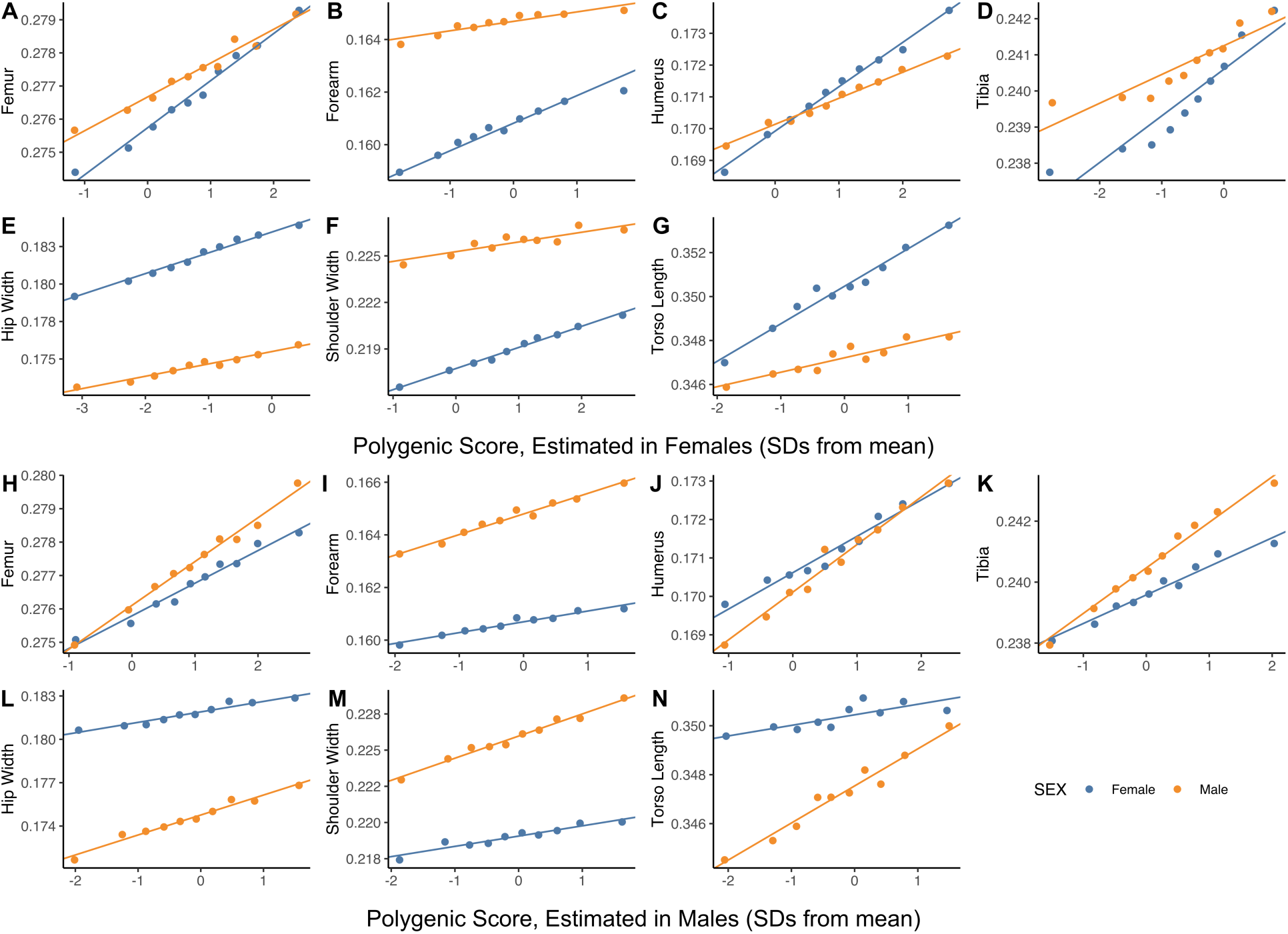
Regression of trait values in males (orange) and separately in females (blue) to a polygenic score estimated in an independent sample of females. Points show mean values in one decile of the polygenic score; the fitted line and associated effect estimate and *R*^2^ correspond to regressions on the raw, non-binned data.

**Fig. S18.**
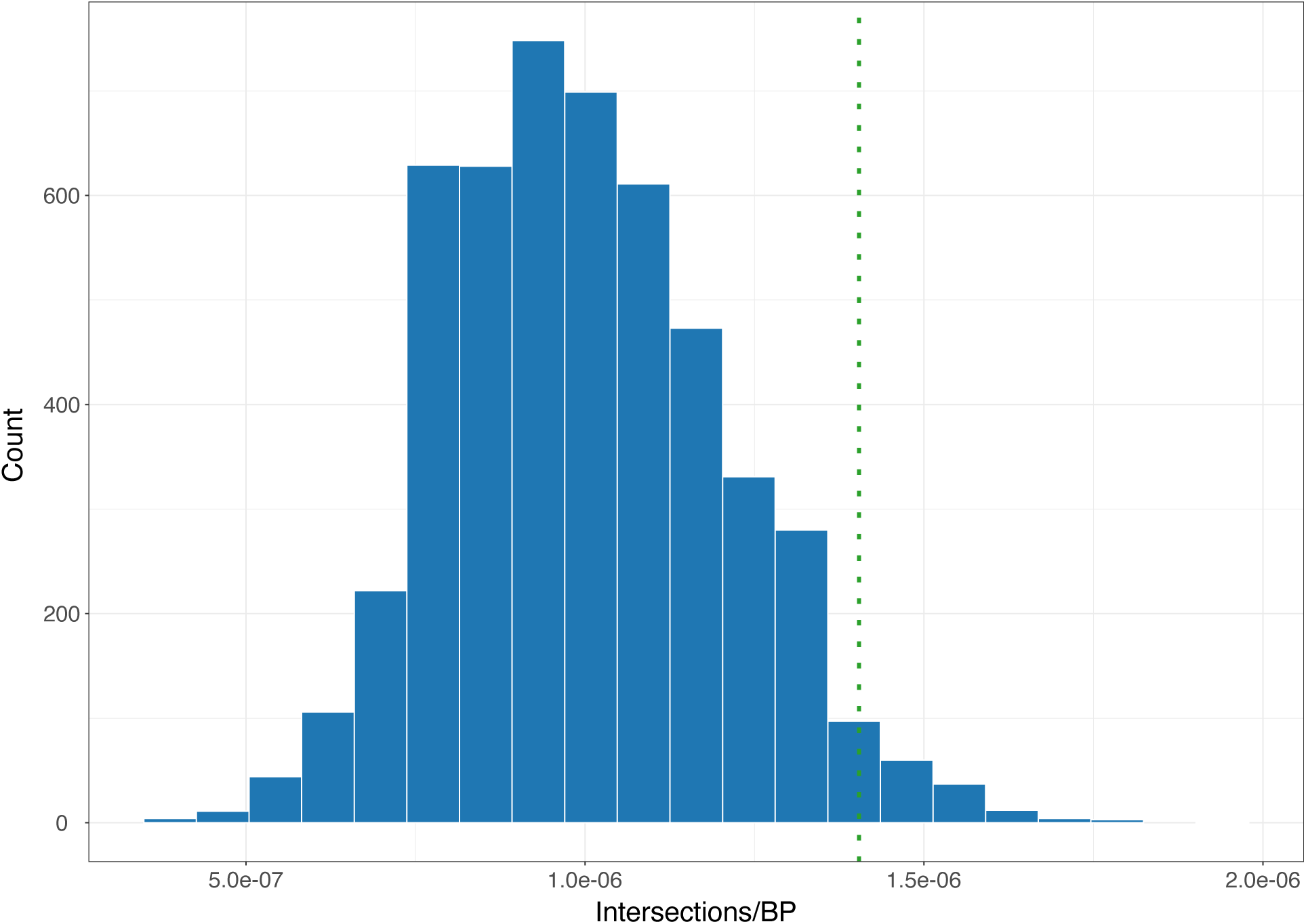
HAR background distributions. Intersections per base pair occurring between human accelerated regions (HARs) and phenotype-associated genes. Blue bars are background distributions generated from 5,000 simulations of matched element sets. An example is shown here for skin pigmentation.

## References

1. L. T. Gruss, D. Schmitt, The evolution of the human pelvis: changing adaptations to bipedalism, obstetrics and thermoregulation. Philosophical Transactions of the Royal Society B: Biological Sciences. 370 (2015), doi:10.1098/RSTB.2014.0063.

2. L. Aiello, C. Dean, J. Cameron, An introduction to human evolutionary anatomy (1990).

3. N. M. Young, G. P. Wagner, B. Hallgrímsson, Development and the evolvability of human limbs. Proc Natl Acad Sci U S A. 107, 3400–3405 (2010).

4. D. J. Futuyma, M. Kirkpatrick, Evolution (Sinauer Associates, Inc., 2017).

5. G. A. Orban, F. Caruana, The neural basis of human tool use. Front Psychol. 5, 310 (2014).

6. W. E. H. Harcourt-Smith, L. C. Aiello, Fossils, feet and the evolution of human bipedal locomotion. J Anat. 204, 403 (2004).

7. F. E. Grine, C. S. Mongle, J. G. Fleagle, A. S. Hammond, The taxonomic attribution of African hominin postcrania from the Miocene through the Pleistocene: Associations and assumptions. J Hum Evol. 173, 103255 (2022).

8. J. T. Stern, R. L. Susman, The locomotor anatomy of Australopithecus afarensis. Am J Phys Anthropol. 60, 279–317 (1983).

9. S. J. Shefelbine, C. Tardieu, D. R. Carter, Development of the femoral bicondylar angle in hominid bipedalism. Bone. 30, 765–770 (2002).

10. R. Schiess, M. Haeusler, No skeletal dysplasia in the nariokotome boy KNM-WT 15000 (homo erectus)—A reassessment of congenital pathologies of the vertebral column. Am J Phys Anthropol. 150, 365–374 (2013).

11. J. Garcia-Fernàndez, The genesis and evolution of homeobox gene clusters. Nature Reviews Genetics 2005 6:12. 6, 881–892 (2005).

12. R. L. Johnson, C. J. Tabin, Molecular Models for Vertebrate Limb Development. Cell. 90, 979–990 (1997).

13. I. Willis, G. Strohl, A homoeotic mutation transforming leg to antenna in Drosophila. Nature 1981 292:5824. 292, 635–638 (1981).

14. J. G. Roscito, K. Sameith, B. M. Kirilenko, N. Hecker, S. Winkler, A. Dahl, M. T. Rodrigues, M. Hiller, Convergent and lineage-specific genomic differences in limb regulatory elements in limbless reptile lineages. Cell Rep. 38, 110280 (2022).

15. E. Z. Kvon, O. K. Kamneva, U. S. Melo, I. Barozzi, M. Osterwalder, B. J. Mannion, V. Tissières, C. S. Pickle, I. Plajzer-Frick, E. A. Lee, M. Kato, T. H. Garvin, J. A. Akiyama, V. Afzal, J. Lopez-Rios, E. M. Rubin, D. E. Dickel, L. A. Pennacchio, A. Visel, Progressive Loss of Function in a Limb Enhancer during Snake Evolution. Cell. 167, 633–642.e11 (2016).

16. A. Saxena, V. Sharma, P. Muthuirulan, S. J. Neufeld, M. P. Tran, H. L. Gutierrez, K. D. Chen, J. M. Erberich, A. Birmingham, T. D. Capellini, J. Cobb, M. Hiller, K. L. Cooper, Interspecies transcriptomics identify genes that underlie disproportionate foot growth in jerboas. Current Biology. 32, 289–303.e6 (2022).

17. B. Xia, W. Zhang, A. Wudzinska, E. Huang, R. Brosh, M. Pour, A. Miller, J. S. Dasen, M. T. Maurano, S. Y. Kim, J. D. Boeke, I. Yanai, The genetic basis of tail-loss evolution in humans and apes. bioRxiv (2021), doi:10.1101/2021.09.14.460388.

18. D. Richard, Z. Liu, J. Cao, A. M. Kiapour, J. Willen, S. Yarlagadda, E. Jagoda, V. B. Kolachalama, J. T. Sieker, G. H. Chang, P. Muthuirulan, M. Young, A. Masson, J. Konrad, S. Hosseinzadeh, D. E. Maridas, V. Rosen, R. Krawetz, N. Roach, T. D. Capellini, Evolutionary Selection and Constraint on Human Knee Chondrocyte Regulation Impacts Osteoarthritis Risk. Cell. 181, 362–381.e28 (2020).

19. S. Chatterjee, N. Das, P. Chatterjee, The estimation of the heritability of anthropometric measurements. Appl Human Sci. 18, 1–7 (1999).

20. K. Silventoinen, S. Sammalisto, M. Perola, D. I. Boomsma, B. K. Cornes, C. Davis, L. Dunkel, M. de Lange, J. R. Harris, J. V. B. Hjelmborg, M. Luciano, N. G. Martin, J. Mortensen, L. Nisticò, N. L. Pedersen, A. Skytthe, T. D. Spector, M. A. Stazi, G. Willemsen, J. Kaprio, Heritability of Adult Body Height: A Comparative Study of Twin Cohorts in Eight Countries. Twin Research and Human Genetics. 6, 399–408 (2003).

21. L. Yengo, S. Vedantam, E. Marouli, J. Sidorenko, E. Bartell, S. Sakaue, M. Graff, A. U. Eliasen, Y. Jiang, S. Raghavan, J. Miao, J. D. Arias, S. E. Graham, R. E. Mukamel, C. N. Spracklen, X. Yin, S.-H. Chen, T. Ferreira, H. H. Highland, Y. Ji, T. Karaderi, K. Lin, K. Lüll, D. E. Malden, C. Medina-Gomez, M. Machado, A. Moore, S. Rüeger, X. Sim, S. Vrieze, T. S. Ahluwalia, M. Akiyama, M. A. Allison, M. Alvarez, M. K. Andersen, A. Ani, V. Appadurai, L. Arbeeva, S. Bhaskar, L. F. Bielak, S. Bollepalli, L. L. Bonnycastle, J. Bork-Jensen, J. P. Bradfield, Y. Bradford, P. S. Braund, J. A. Brody, K. S. Burgdorf, B. E. Cade, H. Cai, Q. Cai, A. Campbell, M. Cañadas-Garre, E. Catamo, J.-F. Chai, X. Chai, L.-C. Chang, Y.-C. Chang, C.-H. Chen, A. Chesi, S. H. Choi, R.-H. Chung, M. Cocca, M. P. Concas, C. Couture, G. Cuellar-Partida, R. Danning, E. W. Daw, F. Degenhard, G. E. Delgado, A. Delitala, A. Demirkan, X. Deng, P. Devineni, A. Dietl, M. Dimitriou, L. Dimitrov, R. Dorajoo, A. B. Ekici, J. E. Engmann, Z. Fairhurst-Hunter, A.-E. Farmaki, J. D. Faul, J.-C. Fernandez-Lopez, L. Forer, M. Francescatto, S. Freitag-Wolf, C. Fuchsberger, T. E. Galesloot, Y. Gao, Z. Gao, F. Geller, O. Giannakopoulou, F. Giulianini, A. P. Gjesing, A. Goel, S. D. Gordon, M. Gorski, J. Grove, X. Guo, S. Gusta fsson, J. Haessler, T. F. Hansen, A. S. Havulinna, S. J. Haworth, J. He, N. Heard-Costa, P. Hebbar, G. Hindy, Y.-L. A. Ho, E. Hofer, E. Holliday, K. Horn, W. E. Hornsby, J.-J. Hottenga, H. Huang, J. Huang, A. Huerta-Chagoya, J. E. Huffman, Y.-J. Hung, S. Huo, M. Y. Hwang, H. Iha, D. D. Ikeda, M. Isono, A. U. Jackson, S. Jäger, I. E. Jansen, I. Johansson, J. B. Jonas, A. Jonsson, T. Jørgensen, I.-P. Kalafati, M. Kanai, S. Kanoni, L. L. Kårhus, A. Kasturiratne, T. Katsuya, T. Kawaguchi, R. L. Kember, K. A. Kentistou, H.-N. Kim, Y. J. Kim, M. E. Kleber, M. J. Knol, A. Kurbasic, M. Lauzon, P. Le, R. Lea, J.-Y. Lee, H. L. Leonard, S. A. Li, X. Li, X. Li, J. Liang, H. Lin, S.-Y. Lin, J. Liu, X. Liu, K. S. Lo, J. Long, L. Lores-Motta, J. Luan, V. Lyssenko, L.-P. Lyytikäinen, A. Mahajan, V. Mamakou, M. Mangino, A. Manichaikul, J. Marten, M. Mattheisen, L. Mavarani, A. F. McDaid, K. Meidtner, T. L. Melendez, J. M. Mercader, Y. Milaneschi, J. E. Miller, I. Y. Millwood, P. P. Mishra, R. E. Mitchell, L. T. Møllehave, A. Morgan, S. Mucha, M. Munz, M. Nakatochi, C. P. Nelson, M. Nethander, C. W. Nho, A. A. Nielsen, I. M. Nolte, S. S. Nongmaithem, R. Noordam, I. Ntalla, T. Nutile, A. Pandit, P. Christofidou, K. Pärna, M. Pauper, E. R. B. Petersen, L. v. Petersen, N. Pitkänen, O. Polašek, A. Poveda, M. H. Preuss, S. Pyarajan, L. M. Raffield, H. Rakugi, J. Ramirez, A. Rasheed, D. Raven, N. W. Rayner, C. Riveros, R. Rohde, D. Ruggiero, S. E. Ruotsalainen, K. A. Ryan, M. Sabater-Lleal, R. Saxena, M. Scholz, A. Sendamarai, B. Shen, J. Shi, J. H. Shin, C. Sidore, C. M. Sitlani, R. C. Slieker, R. A. J. Smit, A. v. Smith, J. A. Smith, L. J. Smyth, L. Southam, V. Steinthorsdottir, L. Sun, F. Takeuchi, D. S. P. Tallapragada, K. D. Taylor, B. O. Tayo, C. Tcheandjieu, N. Terzikhan, P. Tesolin, A. Teumer, E. Theusch, D. J. Thompson, G. Thorleifsson, P. R. H. J. Timmers, S. Trompet, C. Turman, S. Vaccargiu, S. W. van der Laan, P. J. van der Most, J. B. van Klinken, J. van Setten, S. S. Verma, N. Verweij, Y. Veturi, C. A. Wang, C. Wang, L. Wang, Z. Wang, H. R. Warren, W. bin Wei, A. R. Wickremasinghe, M. Wielscher, K. L. Wiggins, B. S. Winsvold, A. Wong, Y. Wu, M. Wuttke, R. Xia, T. Xie, K. Yamamoto, J. Yang, J. Yao, H. Young, N. A. Yousri, L. Yu, L. Zeng, W. Zhang, X. Zhang, J.-H. Zhao, W. Zhao, W. Zhou, M. E. Zimmermann, M. Zoledziewska, L. S. Adair, H. H. H. Adams, C. A. Aguilar-Salinas, F. Al-Mulla, D. K. Arnett, F. W. Asselbergs, B. O. Åsvold, J. Attia, B. Banas, S. Bandinelli, D. A. Bennett, T. Bergler, D. Bharadwaj, G. Biino, H. Bisgaard, E. Boerwinkle, C. A. Böger, K. Bønnelykke, D. I. Boomsma, A. D. Børglum, J. B. Borja, C. Bouchard, D. W. Bowden, I. Brandslund, B. Brumpton, J. E. Buring, M. J. Caulfield, J. C. Chambers, G. R. Chandak, S. J. Chanock, N. Chaturvedi, Y.-D. I. Chen, Z. Chen, C.-Y. Cheng, I. E. Christophersen, M. Ciullo, J. W. Cole, F. S. Collins, R. S. Cooper, M. Cruz, F. Cucca, L. A. Cupples, M. J. Cutler, S. M. Damrauer, T. M. Dantoft, G. J. de Borst, L. C. P. G. M. de Groot, P. L. de Jager, D. P. v. de Kleijn, H. Janaka de Silva, G. v. Dedoussis, A. I. den Hollander, S. Du, D. F. Easton, P. J. M. Elders, A. H. Eliassen, P. T. Ellinor, S. Elmståhl, J. Erdmann, M. K. Evans, D. Fatkin, B. Feenstra, M. F. Feitosa, L. Ferrucci, I. Ford, M. Fornage, A. Franke, P. W. Franks, B. I. Freedman, P. Gasparini, C. Gieger, G. Girotto, M. E. Goddard, Y. M. Golightly, C. Gonzalez-Villalpando, P. Gordon-Larsen, H. Grallert, S. F. A. Grant, N. Grarup, L. Griffiths, V. Gudnason, C. Haiman, H. Hakonarson, T. Hansen, C. A. Hartman, A. T. Hattersley, C. Hayward, S. R. Heckbert, C.-K. Heng, C. Hengstenberg, A. W. Hewitt, H. Hishigaki, C. B. Hoyng, P. L. Huang, W. Huang, S. C. Hunt, K. Hveem, E. Hyppönen, W. G. Iacono, S. Ichihara, M. A. Ikram, C. R. Isasi, R. D. Jackson, M.-R. Jarvelin, Z.-B. Jin, K.-H. Jöckel, P. K. Joshi, P. Jousilahti, J. W. Jukema, M. Kähönen, Y. Kamatani, K. D. Kang, J. Kaprio, S. L. R. Kardia, F. Karpe, N. Kato, F. Kee, T. Kessler, A. v. Khera, C. C. Khor, L. A. L. M. Kiemeney, B.-J. Kim, E. K. Kim, H.-L. Kim, P. Kirchhof, M. Kivimaki, W.-P. Koh, H. A. Koistinen, G. D. Kolovou, J. S. Kooner, C. Kooperberg, A. Köttgen, P. Kovacs, A. Kraaijeveld, P. Kraft, R. M. Krauss, M. Kumari, Z. Kutalik, M. Laakso, L. A. Lange, C. Langenberg, L. J. Launer, L. le Marchand, H. Lee, N. R. Lee, T. Lehtimäki, H. Li, L. Li, W. Lieb, X. Lin, L. Lind, A. Linneberg, C.-T. Liu, J. Liu, M. Loeffler, B. London, S. A. Lubitz, S. J. Lye, D. A. Mackey, R. Mägi, P. K. E. Magnusson, G. M. Marcus, P. M. Vidal, N. G. Martin, W. März, F. Matsuda, R. W. McGarrah, M. McGue, A. J. McKnight, S. E. Medland, D. Mellström, A. Metspalu, B. D. Mitchell, P. Mitchell, D. O. Mook-Kanamori, A. D. Morris, L. A. Mucci, P. B. Munroe, M. A. Nalls, S. Nazarian, A. E. Nelson, M. J. Neville, C. Newton-Cheh, C. S. Nielsen, M. M. Nöthen, C. Ohlsson, A. J. Oldehinkel, L. Orozco, K. Pahkala, P. Pajukanta, C. N. A. Palmer, E. J. Parra, C. Pattaro, O. Pedersen, C. E. Pennell, B. W. J. H. Penninx, L. Perusse, A. Peters, P. A. Peyser, D. J. Porteous, D. Posthuma, C. Power, P. P. Pramstaller, M. A. Province, Q. Qi, J. Qu, D. J. Rader, O. T. Raitakari, S. Ralhan, L. S. Rallidis, D. C. Rao, S. Redline, D. F. Reilly, A. P. Reiner, S. Y. Rhee, P. M. Ridker, M. Rienstra, S. Ripatti, M. D. Ritchie, D. M. Roden, F. R. Rosendaal, J. I. Rotter, I. Rudan, F. Rutters, C. Sabanayagam, D. Saleheen, V. Salomaa, N. J. Samani, D. K. Sanghera, N. Sattar, B. Schmidt, H. Schmidt, R. Schmidt, M. B. Schulze, H. Schunkert, L. J. Scott, R. J. Scott, P. Sever, E. J. Shiroma, M. B. Shoemaker, X.-O. Shu, E. M. Simonsick, M. Sims, J. R. Singh, A. B. Singleton, M. F. Sinner, J. G. Smith, H. Snieder, T. D. Spector, M. J. Stampfer, K. J. Stark, D. P. Strachan, L. M. ‘t Hart, Y. Tabara, H. Tang, J.-C. Tardif, T. A. Thanaraj, N. J. Timpson, A. Tönjes, A. Tremblay, T. Tuomi, J. Tuomilehto, M.-T. Tusié-Luna, A. G. Uitterlinden, R. M. van Dam, P. van der Harst, N. van der Velde, C. M. van Duijn, N. M. van Schoor, V. Vitart, U. Völker, P. Vollenweider, H. Völzke, N. H. Wacher-Rodarte, M. Walker, Y. X. Wang, N. J. Wareham, R. M. Watanabe, H. Watkins, D. R. Weir, T. M. Werge, E. Widen, L. R. Wilkens, G. Willemsen, W. C. Willett, J. F. Wilson, T.-Y. Wong, J.-T. Woo, A. F. Wright, J.-Y. Wu, H. Xu, C. S. Yajnik, M. Yokota, J.-M. Yuan, E. Zeggini, B. S. Zemel, W. Zheng, X. Zhu, J. M. Zmuda, A. B. Zonderman, J.-A. Zwart, G. C. Partida, Y. Sun, D. Croteau-Chonka, J. M. Vonk, S. Chanock, L. le Marchand, D. I. Chasman, Y. S. Cho, I. M. Heid, M. I. McCarthy, M. C. Y. Ng, C. J. O’Donnell, F. Rivadeneira, U. Thorsteinsdottir, Y. v. Sun, E. S. Tai, M. Boehnke, P. Deloukas, A. E. Justice, C. M. Lindgren, R. J. F. Loos, K. L. Mohlke, K. E. North, K. Stefansson, R. G. Walters, T. W. Winkler, K. L. Young, P.-R. Loh, J. Yang, T. Esko, T. L. Assimes, A. Auton, G. R. Abecasis, C. J. Willer, A. E. Locke, S. I. Berndt, G. Lettre, T. M. Frayling, Y. Okada, A. R. Wood, P. M. Visscher, J. N. Hirschhorn, A saturated map of common genetic variants associated with human height. Nature 2022, 1–16 (2022).

22. A. M. Fredriks, S. van Buuren, W. J. M van Heel, R. H. M Dijkman-Neerincx, S. P. Verloove-Vanhorick, J. M. Wit, Nationwide age references for sitting height, leg length, and sitting height/height ratio, and their diagnostic value for disproportionate growth disorders. Arch Dis Child. 90, 807–812 (2005).

23. G. Livshits, A. Roset, K. Yakovenko, S. Trofimov, E. Kobyliansky, Genetics of human body size and shape: body proportions and indices. Ann Hum Biol. 29, 271–289 (2002).

24. S. L. Pulit, C. Stoneman, A. P. Morris, A. R. Wood, C. A. Glastonbury, J. Tyrrell, L. Yengo, T. Ferreira, E. Marouli, Y. Ji, J. Yang, S. Jones, R. Beaumont, D. C. Croteau-Chonka, T. W. Winkler, A. T. Hattersley, R. J. F. Loos, J. N. Hirschhorn, P. M. Visscher, T. M. Frayling, H. Yaghootkar, C. M. Lindgren, Meta-analysis of genome-wide association studies for body fat distribution in 694 649 individuals of European ancestry. Hum Mol Genet. 28, 166–174 (2019).

25. Y. Chan, R. M. Salem, Y. H. H. Hsu, G. McMahon, T. H. Pers, S. Vedantam, T. Esko, M. H. Guo, E. T. Lim, L. Franke, G. D. Smith, D. P. Strachan, J. N. Hirschhorn, Genome-wide Analysis of Body Proportion Classifies Height-Associated Variants by Mechanism of Action and Implicates Genes Important for Skeletal Development. Am J Hum Genet. 96, 695 (2015).

26. H. Currant, P. Hysi, T. W. Fitzgerald, P. Gharahkhani, P. W. M. Bonnemaijer, A. Senabouth, A. W. Hewitt, D. Atan, T. Aung, J. Charng, H. Choquet, J. Craig, P. T. Khaw, C. C. W. Klaver, M. Kubo, J. S. Ong, L. R. Pasquale, C. A. Reisman, M. Daniszewski, J. E. Powell, A. Pébay, M. J. Simcoe, A. A. H. J. Thiadens, C. M. van Duijn, S. Yazar, E. Jorgenson, S. MacGregor, C. J. Hammond, D. A. Mackey, J. L. Wiggs, P. J. Foster, P. J. Patel, E. Birney, A. P. Khawaja, Genetic variation affects morphological retinal phenotypes extracted from UK Biobank optical coherence tomography images. PLoS Genet. 17 (2021), doi:10.1371/JOURNAL.PGEN.1009497.

27. S. Agrawal, M. D. R. Klarqvist, N. Diamant, T. L. Stanley, P. T. Ellinor, N. N. Mehta, A. Philippakis, K. Ng, M. Claussnitzer, S. K. Grinspoon, P. Batra, A. v. Khera, Association of machine learning-derived measures of body fat distribution with cardiometabolic diseases in >40,000 individuals. medRxiv, in press, doi:10.1101/2021.05.07.21256854.

28. W. Bai, H. Suzuki, J. Huang, C. Francis, S. Wang, G. Tarroni, F. Guitton, N. Aung, K. Fung, S. E. Petersen, S. K. Piechnik, S. Neubauer, E. Evangelou, A. Dehghan, D. P. O’Regan, M. R. Wilkins, Y. Guo, P. M. Matthews, D. Rueckert, A population-based phenome-wide association study of cardiac and aortic structure and function. Nat Med. 26 (2020), doi:10.1038/s41591-020-1009-y.

29. J. P. Pirruccello, M. D. Chaffin, E. L. Chou, S. J. Fleming, H. Lin, M. Nekoui, S. Khurshid, S. F. Friedman, A. G. Bick, A. Arduini, L. C. Weng, S. H. Choi, A. D. Akkad, P. Batra, N. R. Tucker, A. W. Hall, C. Roselli, E. J. Benjamin, S. K. Vellarikkal, R. M. Gupta, C. M. Stegmann, D. Juric, J. R. Stone, R. S. Vasan, J. E. Ho, U. Hoffmann, S. A. Lubitz, A. A. Philippakis, M. E. Lindsay, P. T. Ellinor, Deep learning enables genetic analysis of the human thoracic aorta. Nature Genetics 2021 54:1. 54, 40–51 (2021).

30. Centers for Disease Control and Prevention (CDC), Prevalence and most common causes of disability among adults--United States, 2005. MMWR Morb Mortal Wkly Rep. 58, 421– 6 (2009).

31. A. A. Guccione, D. T. Felson, J. J. Anderson, J. M. Anthony, Y. Zhang, P. W. F. Wilson, M. Kelly-Hayes, P. A. Wolf, B. E. Kreger, W. B. Kannel, The effects of specific medical conditions on the functional limitations of elders in the Framingham Study. Am J Public Health. 84, 351–358 (1994).

32. D. Chen, J. Shen, W. Zhao, T. Wang, L. Han, J. L. Hamilton, H. J. Im, Osteoarthritis: toward a comprehensive understanding of pathological mechanism. Bone Res. 5, 16044 (2017).

33. K. J. Murray, M. F. Azari, Leg length discrepancy and osteoarthritis in the knee, hip and lumbar spine. J Can Chiropr Assoc. 59, 226 (2015).

34. J. C. Baker-LePain, N. E. Lane, Role of Bone Architecture and Anatomy in Osteoarthritis. Bone. 51, 197 (2012).

35. K. Sun, B. Xiao, D. Liu, J. Wang, Deep High-Resolution Representation Learning for Human Pose Estimation. Proceedings of the IEEE Computer Society Conference on Computer Vision and Pattern Recognition. 2019-**June**, 5686–5696 (2019).

36. J. Deng, W. Dong, R. Socher, L.-J. Li, Kai Li, Li Fei-Fei, ImageNet: A large-scale hierarchical image database, 248–255 (2010).

37. T.-Y. Lin, M. Maire, S. Belongie, L. Bourdev, R. Girshick, J. Hays, P. Perona, D. Ramanan, C. L. Zitnick, P. Dolí, Microsoft COCO: Common Objects in Context (2015).

38. MPII Human Pose Benchmark (Pose Estimation) | Papers With Code, (available at https://paperswithcode.com/sota/pose-estimation-on-mpii-human-pose).

39. COCO test-dev Benchmark (Pose Estimation) | Papers With Code, (available at https://paperswithcode.com/sota/pose-estimation-on-coco-test-dev).

40. K. He, X. Zhang, S. Ren, J. Sun, Deep Residual Learning for Image Recognition. Proceedings of the IEEE Computer Society Conference on Computer Vision and Pattern Recognition. 2016**-December**, 770–778 (2015).

41. C. Bycroft, C. Freeman, D. Petkova, G. Band, L. T. Elliott, K. Sharp, A. Motyer, D. Vukcevic, O. Delaneau, J. O’Connell, A. Cortes, S. Welsh, A. Young, M. Effingham, G. McVean, S. Leslie, N. Allen, P. Donnelly, J. Marchini, The UK Biobank resource with deep phenotyping and genomic data. Nature 2018 562:7726. 562, 203–209 (2018).

42. K. Robinette, T. Churchill, J. McConville, A Comparison of Male and Female Body Sizes and Proportions. Anthropology Research Project, (1979).

43. P. R. Loh, G. Kichaev, S. Gazal, A. P. Schoech, A. L. Price, Mixed model association for biobank-scale data sets. Nat Genet. 50, 906 (2018).

44. B. Bulik-Sullivan, P. R. Loh, H. K. Finucane, S. Ripke, J. Yang, N. Patterson, M. J. Daly, L. Price, B. M. Neale, A. Corvin, J. T. R. Walters, K. H. Farh, P. A. Holmans, P. Lee, D. A. Collier, H. Huang, T. H. Pers, I. Agartz, E. Agerbo, M. Albus, M. Alexander, F. Amin, S. A. Bacanu, M. Begemann, R. A. Belliveau, J. Bene, S. E. Bergen, E. Bevilacqua, T. B. Bigdeli, D. W. Black, R. Bruggeman, N. G. Buccola, R. L. Buckner, W. Byerley, W. Cahn, G. Cai, M. J. Cairns, D. Campion, R. M. Cantor, V. J. Carr, N. Carrera, S. v. Catts, K. D. Chambert, R. C. K. Chan, R. Y. L. Chen, E. Y. H. Chen, W. Cheng, E. F. C. Cheung, S. A. Chong, C. R. Cloninger, D. Cohen, N. Cohen, P. Cormican, N. Craddock, B. Crespo-Facorro, J. J. Crowley, D. Curtis, M. Davidson, K. L. Davis, F. Degenhardt, J. del Favero, L. E. DeLisi, D. Demontis, D. Dikeos, T. Dinan, S. Djurovic, G. Donohoe, E. Drapeau, J. Duan, F. Dudbridge, N. Durmishi, P. Eichhammer, J. Eriksson, V. Escott-Price, L. Essioux, A. H. Fanous, M. S. Farrell, J. Frank, L. Franke, R. Freedman, N. B. Freimer, M. Friedl, J. I. Friedman, M. Fromer, G. Genovese, L. Georgieva, E. S. Gershon, I. Giegling, P. Giusti-Rodríguez, S. Godard, J. I. Goldstein, V. Golimbet, S. Gopal, J. Gratten, L. de Haan, C. Hammer, M. L. Hamshere, M. Hansen, T. Hansen, V. Haroutunian, A. M. Hartmann, F. A. Henskens, S. Herms, J. N. Hirschhorn, P. Hoffmann, Hofman, M. v. Hollegaard, D. M. Hougaard, M. Ikeda, I. Joa, A. Juliá, R. S. Kahn, L. Kalaydjieva, S. Karachanak-Yankova, J. Karjalainen, D. Kavanagh, M. C. Keller, B. J. Kelly, J. L. Kennedy, A. Khrunin, Y. Kim, J. Klovins, J. A. Knowles, B. Konte, V. Kucinskas, Z. A. Kucinskiene, H. Kuzelova-Ptackova, A. K. Kähler, C. Laurent, J. L. C. Keong, S. H. Lee, S. E. Legge, B. Lerer, M. Li, T. Li, K. Y. Liang, J. Lieberman, S. Limborska, C. M. Loughland, J. Lubinski, J. Lönnqvist, M. Macek, P. K. E. Magnusson, S. Maher, W. Maier, J. Mallet, S. Marsal, M. Mattheisen, M. Mattingsdal, R. W. McCarley, C. McDonald, A. M. McIntosh, S. Meier, C. J. Meijer, B. Melegh, I. Melle, R. I. Mesholam-Gately, A. Metspalu, P. T. Michie, L. Milani, V. Milanova, Y. Mokrab, D. W. Morris, O. Mors, K. C. Murphy, R. M. Murray, I. Myin-Germeys, B. Müller-Myhsok, M. Nelis, I. Nenadic, D. A. Nertney, G. Nestadt, K. K. Nicodemus, L. Nikitina-Zake, L. Nisenbaum, A. Nordin, E. O’Callaghan, C. O’Dushlaine, F. A. O’Neill, S. Y. Oh, A. Olincy, L. Olsen, J. van Os, C. Pantelis, G. N. Papadimitriou, S. Papiol, E. Parkhomenko, M. T. Pato, T. Paunio, M. Pejovic-Milovancevic, D. O. Perkins, O. Pietiläinen, J. Pimm, A. J. Pocklington, J. Powell, A. E. Pulver, S. M. Purcell, D. Quested, H. B. Rasmussen, A. Reichenberg, M. A. Reimers, A. L. Richards, J. L. Roffman, P. Roussos, D. M. Ruderfer, V. Salomaa, A. R. Sanders, U. Schall, C. R. Schubert, T. G. Schulze, S. G. Schwab, E. M. Scolnick, R. J. Scott, L. J. Seidman, J. Shi, E. Sigurdsson, T. Silagadze, J. M. Silverman, K. Sim, P. Slominsky, J. W. Smoller, H. C. So, C. C. A. Spencer, E. A. Stahl, H. Stefansson, S. Steinberg, E. Stogmann, R. E. Straub, E. Strengman, J. Strohmaier, T. S. Stroup, M. Subramaniam, J. Suvisaari, D. M. Svrakic, J. P. Szatkiewicz, E. Söderman, S. Thirumalai, D. Toncheva, P. A. Tooney, S. Tosato, J. Veijola, J. Waddington, D. Walsh, D. Wang, Q. Wang, B. T. Webb, M. Weiser, D. B. Wildenauer, N. M. Williams, S. Williams, S. H. Witt, A. R. Wolen, E. H. M. Wong, B. K. Wormley, J. Q. Wu, H. S. Xi, C. C. Zai, X. Zheng, F. Zimprich, N. R. Wray, K. Stefansson, P. M. Visscher, R. Adolfsson, O. A. Andreassen, D. H. R. Blackwood, E. Bramon, J. D. Buxbaum, A. D. Børglum, S. Cichon, A. Darvasi, E. Domenici, H. Ehrenreich, T. Esko, P. v. Gejman, M. Gill, H. Gurling, C. M. Hultman, N. Iwata, A. v. Jablensky, E. G. Jönsson, K. S. Kendler, G. Kirov, J. Knight, T. Lencz, D. F. Levinson, Q. S. Li, J. Liu, A. K. Malhotra, S. A. McCarroll, A. McQuillin, J. L. Moran, P. B. Mortensen, B. J. Mowry, M. M. Nöthen, R. A. Ophoff, M. J. Owen, A. Palotie, C. N. Pato, T. L. Petryshen, D. Posthuma, M. Rietschel, B. P. Riley, D. Rujescu, P. C. Sham, P. Sklar, D. St Clair, D. R. Weinberger, J. R. Wendland, T. Werge, P. F. Sullivan, M. C. O’Donovan, LD Score regression distinguishes confounding from polygenicity in genome-wide association studies. Nature Genetics 2015 47:3. 47, 291–295 (2015).

45. J. Yang, B. Benyamin, B. P. McEvoy, S. Gordon, A. K. Henders, D. R. Nyholt, P. A. Madden, A. C. Heath, N. G. Martin, G. W. Montgomery, M. E. Goddard, P. M. Visscher, Common SNPs explain a large proportion of the heritability for human height. Nature Genetics 2010 42:7. **42**, 565–569 (2010).

46. A. D. Grotzinger, M. Rhemtulla, R. de Vlaming, S. J. Ritchie, T. T. Mallard, W. D. Hill, H. F. Ip, R. E. Marioni, A. M. McIntosh, I. J. Deary, P. D. Koellinger, K. P. Harden, M. G. Nivard, E. M. Tucker-Drob, Genomic structural equation modelling provides insights into the multivariate genetic architecture of complex traits. Nature Human Behaviour 2019 3:5. 3, 513–525 (2019).

47. C. Zhu, M. J. Ming, J. M. Cole, M. Kirkpatrick, A. Harpak, Amplification is the Primary Mode of Gene-by-Sex Interaction in Complex Human Traits. bioRxiv, in press, doi:10.1101/2022.05.06.490973.

48. K. Watanabe, E. Taskesen, A. van Bochoven, D. Posthuma, Functional mapping and annotation of genetic associations with FUMA. Nature Communications 2017 8:1. 8, 1–11 (2017).

49. Online Mendelian Inheritance in Man, OMIM®. McKusick-Nathans Institute of Genetic Medicine, Johns Hopkins University (Baltimore, MD).

50. J. A. Blake, R. Baldarelli, J. A. Kadin, J. E. Richardson, C. L. Smith, C. J. Bult, Mouse Genome Database (MGD): Knowledgebase for mouse-human comparative biology. Nucleic Acids Res. 49, D981–D987 (2021).

51. I. Delgado, G. Giovinazzo, S. Temiño, Y. Gauthier, A. Balsalobre, J. Drouin, M. Torres, Control of mouse limb initiation and antero-posterior patterning by Meis transcription factors. Nature Communications 2021 12:1. 12, 1–13 (2021).

52. Z. Liu, A. A. Hussien, Y. Wang, T. Heckmann, R. Gonzalez, C. M. Karner, J. G. Snedeker, R. S. Gray, An adhesion g protein-coupled receptor is required in cartilaginous and dense connective tissues to maintain spine alignment. Elife. 10 (2021), doi:10.7554/ELIFE.67781.

53. F. Aguet, A. A. Brown, S. E. Castel, J. R. Davis, Y. He, B. Jo, P. Mohammadi, Y. S. Park, P. Parsana, A. v. Segrè, B. J. Strober, Z. Zappala, B. B. Cummings, E. T. Gelfand, K. Hadley, K. H. Huang, M. Lek, X. Li, J. L. Nedzel, D. Y. Nguyen, M. S. Noble, T. J. Sullivan, T. Tukiainen, D. G. MacArthur, G. Getz, A. Addington, P. Guan, S. Koester, A. R. Little, N. C. Lockhart, H. M. Moore, A. Rao, J. P. Struewing, S. Volpi, L. E. Brigham, R. Hasz, M. Hunter, C. Johns, M. Johnson, G. Kopen, W. F. Leinweber, J. T. Lonsdale, A. McDonald, B. Mestichelli, K. Myer, B. Roe, M. Salvatore, S. Shad, J. A. Thomas, G. Walters, M. Washington, J. Wheeler, J. Bridge, B. A. Foster, B. M. Gillard, E. Karasik, R. Kumar, M. Miklos, M. T. Moser, S. D. Jewell, R. G. Montroy, D. C. Rohrer, D. Valley, D. C. Mash, D. A. Davis, L. Sobin, M. E. Barcus, P. A. Branton, N. S. Abell, B. Balliu, O. Delaneau, L. Frésard, E. R. Gamazon, D. Garrido-Martín, A. D. H. Gewirtz, G. Gliner, M. J. Gloudemans, B. Han, A. Z. He, F. Hormozdiari, X. Li, B. Liu, E. Y. Kang, I. C. McDowell, H. Ongen, J. J. Palowitch, C. B. Peterson, G. Quon, S. Ripke, A. Saha, A. A. Shabalin, T. C. Shimko, J. H. Sul, N. A. Teran, E. K. Tsang, H. Zhang, Y. H. Zhou, C. D. Bustamante, N. J. Cox, R. Guigó, M. Kellis, M. I. McCarthy, D. F. Conrad, E. Eskin, G. Li, A. B. Nobel, C. Sabatti, B. E. Stranger, X. Wen, F. A. Wright, K. G. Ardlie, E. T. Dermitzakis, T. Lappalainen, A. Battle, C. D. Brown, B. E. Engelhardt, S. B. Montgomery, R. E. Handsaker, S. Kashin, K. J. Karczewski, D. T. Nguyen, C. A. Trowbridge, R. Barshir, O. Basha, G. K. Bogu, L. S. Chen, C. Chiang, F. N. Damani, P. G. Ferreira, I. M. Hall, C. Howald, H. K. Im, Y. Kim, S. Kim-Hellmuth, S. Mangul, J. Monlong, M. Muñoz-Aguirre, A. W. Ndungu, D. L. Nicolae, M. Oliva, N. Panousis, P. Papasaikas, A. J. Payne, J. Quan, F. Reverter, M. Sammeth, A. J. Scott, R. Sodaei, M. Stephens, S. Urbut, M. van de Bunt, G. Wang, H. S. Xi, E. Yeger-Lotem, J. B. Zaugg, J. M. Akey, D. Bates, J. Chan, M. Claussnitzer, K. Demanelis, M. Diegel, J. A. Doherty, A. P. Feinberg, M. S. Fernando, J. Halow, K. D. Hansen, E. Haugen, P. F. Hickey, L. Hou, F. Jasmine, R. Jian, L. Jiang, A. Johnson, R. Kaul, M. G. Kibriya, K. Lee, J. B. Li, Q. Li, J. Lin, S. Lin, S. Linder, C. Linke, Y. Liu, M. T. Maurano, B. Molinie, J. Nelson, F. J. Neri, Y. Park, B. L. Pierce, N. J. Rinaldi, L. F. Rizzardi, R. Sandstrom, A. Skol, K. S. Smith, M. P. Snyder, J. Stamatoyannopoulos, H. Tang, L. Wang, M. Wang, N. van Wittenberghe, F. Wu, R. Zhang, C. R. Nierras, L. J. Carithers, J. B. Vaught, S. E. Gould, N. C. Lockart, C. Martin, A. M. Addington, S. E. Koester, A. H. Undale, A. M. Smith, D. E. Tabor, N. v. Roche, J. A. McLean, N. Vatanian, K. L. Robinson, K. M. Valentino, L. Qi, S. Hunter, P. Hariharan, S. Singh, K. S. Um, T. Matose, M. M. Tomaszewski, L. K. Barker, M. Mosavel, L. A. Siminoff, H. M. Traino, P. Flicek, T. Juettemann, M. Ruffier, D. Sheppard, K. Taylor, S. J. Trevanion, D. R. Zerbino, B. Craft, M. Goldman, M. Haeussler, W. J. Kent, C. M. Lee, B. Paten, K. R. Rosenbloom, J. Vivian, J. Zhu, Genetic effects on gene expression across human tissues. Nature. 550 (2017), doi:10.1038/nature24277.

54. T. Funck-Brentano, M. Nethander, S. Movérare-Skrtic, P. Richette, C. Ohlsson, Causal Factors for Knee, Hip, and Hand Osteoarthritis: A Mendelian Randomization Study in the UK Biobank. Arthritis and Rheumatology. 71, 1634–1641 (2019).

55. T. Ge, C. Y. Chen, Y. Ni, Y. C. A. Feng, J. W. Smoller, Polygenic prediction via Bayesian regression and continuous shrinkage priors. Nature Communications 2019 10:1. 10, 1–10 (2019).

56. K. Xu, E. E. Schadt, K. S. Pollard, P. Roussos, J. T. Dudley, Genomic and Network Patterns of Schizophrenia Genetic Variation in Human Evolutionary Accelerated Regions. Mol Biol Evol. 32, 1148 (2015).

57. P. M. Thompson, N. Jahanshad, C. R. K. Ching, L. E. Salminen, S. I. Thomopoulos, J. Bright, B. T. Baune, S. Bertolín, J. Bralten, W. B. Bruin, R. Bülow, J. Chen, Y. Chye, U. Dannlowski, C. G. F. de Kovel, G. Donohoe, L. T. Eyler, S. v. Faraone, P. Favre, C. A. Filippi, T. Frodl, D. Garijo, Y. Gil, H. J. Grabe, K. L. Grasby, T. Hajek, L. K. M. Han, S. N. Hatton, K. Hilbert, T. C. Ho, L. Holleran, G. Homuth, N. Hosten, J. Houenou, I. Ivanov, T. Jia, S. Kelly, M. Klein, J. S. Kwon, M. A. Laansma, J. Leerssen, U. Lueken, A. Nunes, J. O. Neill, N. Opel, F. Piras, F. Piras, M. C. Postema, E. Pozzi, N. Shatokhina, C. Soriano-Mas, G. Spalletta, D. Sun, A. Teumer, A. K. Tilot, L. Tozzi, C. van der Merwe, E. J. W. van Someren, G. A. van Wingen, H. Völzke, E. Walton, L. Wang, A. M. Winkler, K. Wittfeld, M. J. Wright, J. Y. Yun, G. Zhang, Y. Zhang-James, B. M. Adhikari, I. Agartz, M. Aghajani, A. Aleman, R. R. Althoff, A. Altmann, O. A. Andreassen, D. A. Baron, B. L. Bartnik-Olson, J. Marie Bas-Hoogendam, A. R. Baskin-Sommers, C. E. Bearden, L. A. Berner, P. S. W. Boedhoe, R. M. Brouwer, J. K. Buitelaar, K. Caeyenberghs, C. A. M. Cecil, R. A. Cohen, J. H. Cole, P. J. Conrod, S. A. de Brito, S. M. C. de Zwarte, E. L. Dennis, S. Desrivieres, D. Dima, S. Ehrlich, C. Esopenko, G. Fairchild, S. E. Fisher, J. P. Fouche, C. Francks, S. Frangou, B. Franke, H. P. Garavan, D. C. Glahn, N. A. Groenewold, T. P. Gurholt, B. A. Gutman, T. Hahn, I. H. Harding, D. Hernaus, D. P. Hibar, F. G. Hillary, M. Hoogman, H. E. Hulshoff Pol, M. Jalbrzikowski, G. A. Karkashadze, E. T. Klapwijk, R. C. Knickmeyer, P. Kochunov, I. K. Koerte, X. Z. Kong, S. L. Liew, A. P. Lin, M. W. Logue, E. Luders, F. Macciardi, S. Mackey, A. R. Mayer, C. R. McDonald, A. B. McMahon, S. E. Medland, G. Modinos, R. A. Morey, S. C. Mueller, P. Mukherjee, L. Namazova-Baranova, T. M. Nir, A. Olsen, P. Paschou, D. S. Pine, F. Pizzagalli, M. E. Rentería, J. D. Rohrer, P. G. Sämann, L. Schmaal, G. Schumann, M. S. Shiroishi, S. M. Sisodiya, D. J. A. Smit, I. E. Sønderby, D. J. Stein, J. L. Stein, M. Tahmasian, D. F. Tate, J. A. Turner, O. A. van den Heuvel, N. J. A. van der Wee, Y. D. van der Werf, T. G. M. van Erp, N. E. M. van Haren, D. van Rooij, L. S. van Velzen, I. M. Veer, D. J. Veltman, J. E. Villalon-Reina, H. Walter, C. D. Whelan, E. A. Wilde, M. Zarei, V. Zelman, ENIGMA and global neuroscience: A decade of large-scale studies of the brain in health and disease across more than 40 countries. Translational Psychiatry 2020 10:1. 10, 1–28 (2020).

58. M. Sohail, Investigating relative contributions to psychiatric disease architecture from sequence elements originating across multiple evolutionary time-scales. bioRxiv (2022), doi:10.1101/2022.02.28.482389.

59. M. L. A. Hujoel, S. Gazal, F. Hormozdiari, B. van de Geijn, A. L. Price, Disease Heritability Enrichment of Regulatory Elements Is Concentrated in Elements with Ancient Sequence Age and Conserved Function across Species. Am J Hum Genet. 104, 611–624 (2019).

60. S. K. Reilly, J. Yin, A. E. Ayoub, D. Emera, J. Leng, J. Cotney, R. Sarro, P. Rakic, J. P. Noonan, Evolutionary changes in promoter and enhancer activity during human corticogenesis. Science (1979). 347, 1155–1159 (2015).

61. M. W. Vermunt, S. C. Tan, B. Castelijns, G. Geeven, P. Reinink, E. de Bruijn, I. Kondova, S. Persengiev, R. Bontrop, E. Cuppen, W. de Laat, M. P. Creyghton, Epigenomic annotation of gene regulatory alterations during evolution of the primate brain. Nat Neurosci. 19, 494–503 (2016).

62. S. R. Browning, B. L. Browning, Y. Zhou, S. Tucci, J. M. Akey, Analysis of Human Sequence Data Reveals Two Pulses of Archaic Denisovan Admixture. Cell. 173, 53–61.e9 (2018).

63. S. Peyregne, M. J. Boyle, M. Dannemann, K. Prufer, Detecting ancient positive selection in humans using extended lineage sorting. Genome Res. 27, 1563–1572 (2017).

64. H. K. Finucane, B. Bulik-Sullivan, A. Gusev, G. Trynka, Y. Reshef, P. R. Loh, V. Anttila, H. Xu, C. Zang, K. Farh, S. Ripke, F. R. Day, S. Purcell, E. Stahl, S. Lindstrom, J. R. B. Perry, Y. Okada, S. Raychaudhuri, M. J. Daly, N. Patterson, B. M. Neale, A. L. Price, Partitioning heritability by functional annotation using genome-wide association summary statistics. Nature Genetics 2015 47:11. 47, 1228–1235 (2015).

65. S. Gazal, H. K. Finucane, N. A. Furlotte, P. R. Loh, P. F. Palamara, X. Liu, A. Schoech, B. Bulik-Sullivan, B. M. Neale, A. Gusev, A. L. Price, Linkage disequilibrium–dependent architecture of human complex traits shows action of negative selection. Nature Genetics 2017 49:10. 49, 1421–1427 (2017).

66. D. Marnetto, F. Mantica, I. Molineris, E. Grassi, I. Pesando, P. Provero, Evolutionary Rewiring of Human Regulatory Networks by Waves of Genome Expansion. Am J Hum Genet. 102, 207 (2018).

67. Roadmap Epigenomics Consortium, A. Kundaje, W. Meuleman, J. Ernst, M. Bilenky, A. Yen, A. Heravi-Moussavi, P. Kheradpour, Z. Zhang, J. Wang, M. J. Ziller, V. Amin, J. W. Whitaker, M. D. Schultz, L. D. Ward, A. Sarkar, G. Quon, R. S. Sandstrom, M. L. Eaton, Y. C. Wu, A. R. Pfenning, X. Wang, M. Claussnitzer, Y. Liu, C. Coarfa, R. A. Harris, N. Shoresh, C. B. Epstein, E. Gjoneska, D. Leung, W. Xie, R. D. Hawkins, R. Lister, C. Hong, P. Gascard, A. J. Mungall, R. Moore, E. Chuah, A. Tam, T. K. Canfield, R. S. Hansen, R. Kaul, P. J. Sabo, M. S. Bansal, A. Carles, J. R. Dixon, K. H. Farh, S. Feizi, R. Karlic, A. R. Kim, A. Kulkarni, D. Li, R. Lowdon, G. Elliott, T. R. Mercer, S. J. Neph, V. Onuchic, P. Polak, N. Rajagopal, P. Ray, R. C. Sallari, K. T. Siebenthall, N. A. Sinnott-Armstrong, M. Stevens, R. E. Thurman, J. Wu, B. Zhang, X. Zhou, A. E. Beaudet, L. A. Boyer, P. L. de Jager, P. J. Farnham, S. J. Fisher, D. Haussler, S. J. M. Jones, W. Li, M. A. Marra, M. T. McManus, S. Sunyaev, J. A. Thomson, T. D. Tlsty, L. H. Tsai, W. Wang, R. A. Waterland, M. Q. Zhang, L. H. Chadwick, B. E. Bernstein, J. F. Costello, J. R. Ecker, M. Hirst, A. Meissner, A. Milosavljevic, B. Ren, J. A. Stamatoyannopoulos, T. Wang, M. Kellis, Integrative analysis of 111 reference human epigenomes. Nature 2015 518:7539. 518, 317–330 (2015).

68. X. Wei, C. R. Robles, A. Pazokitoroudi, A. Ganna, A. Gusev, A. Durvasula, S. Gazal, P.-R. Loh, D. Reich, S. Sankararaman, The lingering effects of Neanderthal introgression on human complex traits. bioRxiv, in press, doi:10.1101/2022.06.07.495223.

69. M. Marchini, L. M. Sparrow, M. N. Cosman, A. Dowhanik, C. B. Krueger, B. Hallgrimsson, C. Rolian, Impacts of genetic correlation on the independent evolution of body mass and skeletal size in mammals. BMC Evol Biol. 14 (2014), doi:10.1186/S12862-014-0258-0.

70. H. Aschard, B. J. Vilhjálmsson, A. D. Joshi, A. L. Price, P. Kraft, Adjusting for Heritable Covariates Can Bias Effect Estimates in Genome-Wide Association Studies. Am J Hum Genet. 96, 329 (2015).

71. F. R. Day, P. R. Loh, R. A. Scott, K. K. Ong, J. R. B. Perry, A Robust Example of Collider Bias in a Genetic Association Study. Am J Hum Genet. 98, 392 (2016).

72. A. S. Nicholls, A. Kiran, T. C. B. Pollard, D. J. Hart, C. P. A. Arden, T. Spector, H. S. Gill, D. W. Murray, A. J. Carr, N. K. Arden, The association between hip morphology parameters and nineteen-year risk of end-stage osteoarthritis of the hip: A nested case– control study. Arthritis Rheum. 63, 3392 (2011).

73. T. M. Ecker, M. Tannast, M. Puls, K. A. Siebenrock, S. B. Murphy, Pathomorphologic alterations predict presence or absence of hip osteoarthrosis. Clin Orthop Relat Res. 465, 46–52 (2007).

74. J. Cushnaghan, P. Dieppe, Study of 500 patients with limb joint osteoarthritis. I. Analysis by age, sex, and distribution of symptomatic joint sites. Ann Rheum Dis. 50, 8–13 (1991).

75. R. Ganz, M. Leunig, K. Leunig-Ganz, W. H. Harris, The etiology of osteoarthritis of the hip: an integrated mechanical concept. Clin Orthop Relat Res. 466, 264–272 (2008).

76. M. Grotle, K. B. Hagen, B. Natvig, F. A. Dahl, T. K. Kvien, Obesity and osteoarthritis in knee, hip and/or hand: an epidemiological study in the general population with 10 years follow-up. BMC Musculoskelet Disord. 9 (2008), doi:10.1186/1471-2474-9-132.

77. A. A. Wright, C. Cook, J. H. Abbott, Variables associated with the progression of hip osteoarthritis: a systematic review. Arthritis Rheum. 61, 925–936 (2009).

78. Studies on Dysplastic Acetabula and Congenital Subluxation of the Hip Joint with Special Reference to the Complication of Osteo-Arthritis. J Am Med Assoc. 115, 81–81 (1940).

79. P. L. S. Li, R. Ganz, Morphologic features of congenital acetabular dysplasia: one in six is retroverted. Clin Orthop Relat Res. 416, 245–253 (2003).

80. K. K. Gosvig, S. Jacobsen, H. Palm, S. Sonne-Holm, E. Magnusson, A new radiological index for assessing asphericity of the femoral head in cam impingement. J Bone Joint Surg Br. 89, 1309–1316 (2007).

81. W. Harris, The correlation between minor or unrecognized developmental deformities and the development of osteoarthritis of the hip - PubMed. Instr Course Lect. 58, 257–259 (2009).

82. H. P. Nötzli, T. F. Wyss, C. H. Stoecklin, M. R. Schmid, K. Treiber, J. Hodler, The contour of the femoral head-neck junction as a predictor for the risk of anterior impingement. J Bone Joint Surg Br. 84, 556–560 (2002).

83. T. C. B. Pollard, R. N. Villar, M. R. Norton, E. D. Fern, M. R. Williams, D. J. Simpson, W. Murray, A. J. Carr, Femoroacetabular impingement and classification of the cam deformity: the reference interval in normal hips. Acta Orthop. 81, 134–141 (2010).

84. T. W. Holliday, Postcranial evidence of cold adaptation in European Neandertals. American Journal of Biological Anthropology. 104, 245–258 (1998).

85. E. Trinkaus, The Neandertals and Modern Human Origins. Annu Rev Anthropol. 15, 193– 218 (1986).

86. C. B. Ruff, Climate and body shape in hominid evolution. J Hum Evol. 21, 81–105 (1991).

87. K. L. Steudel-Numbers, M. J. Tilkens, The effect of lower limb length on the energetic cost of locomotion: implications for fossil hominins. J Hum Evol. 47, 95–109 (2004).

88. M. J. Tilkens, C. Wall-Scheffler, T. D. Weaver, K. Steudel-Numbers, The effects of body proportions on thermoregulation: an experimental assessment of Allen’s rule. J Hum Evol. 53, 286–291 (2007).

89. J. Howard, S. Gugger, Fastai: A Layered API for Deep Learning. Information 2020, Vol. 11, Page 108. 11, 108 (2020).

90. D. Mason, scaramallion, mrbean-bremen, rhaxton, J. Suever, Vanessasaurus, D. P. Orfanos, G. Lemaitre, A. Panchal, A. Rothberg, M. D. Herrmann, J. Massich, J. Kerns, K. van Golen, T. Robitaille, S. Biggs, moloney, C. Bridge, M. Shun-Shin, B. Conrad, pawelzajdel, M. Mattes, Y. Lyu, F. C. Morency, T. Cogan, H. Meine, J. Wortmann, pydicom/pydicom: pydicom 2.3.0 (2022), doi:10.5281/ZENODO.6394735.

91. C. R. Harris, K. J. Millman, S. J. van der Walt, R. Gommers, P. Virtanen, D. Cournapeau, Wieser, J. Taylor, S. Berg, N. J. Smith, R. Kern, M. Picus, S. Hoyer, M. H. van Kerkwijk, M. Brett, A. Haldane, J. F. del Río, M. Wiebe, P. Peterson, P. Gérard-Marchant, K. Sheppard, T. Reddy, W. Weckesser, H. Abbasi, C. Gohlke, T. E. Oliphant, Array programming with NumPy. Nature 2020 585:7825. 585, 357–362 (2020).

92. P. Virtanen, R. Gommers, T. E. Oliphant, M. Haberland, T. Reddy, D. Cournapeau, E. Burovski, P. Peterson, W. Weckesser, J. Bright, S. J. van der Walt, M. Brett, J. Wilson, K. J. Millman, N. Mayorov, A. R. J. Nelson, E. Jones, R. Kern, E. Larson, C. J. Carey, İ. Polat, Y. Feng, E. W. Moore, J. VanderPlas, D. Laxalde, J. Perktold, R. Cimrman, I. Henriksen, E. A. Quintero, C. R. Harris, A. M. Archibald, A. H. Ribeiro, F. Pedregosa, P. van Mulbregt, A. Vijaykumar, A. pietro Bardelli, A. Rothberg, A. Hilboll, A. Kloeckner, A. Scopatz, A. Lee, A. Rokem, C. N. Woods, C. Fulton, C. Masson, C. Häggström, C. Fitzgerald, D. A. Nicholson, D. R. Hagen, D. v. Pasechnik, E. Olivetti, E. Martin, E. Wieser, F. Silva, F. Lenders, F. Wilhelm, G. Young, G. A. Price, G. L. Ingold, G. E. Allen, G. R. Lee, H. Audren, I. Probst, J. P. Dietrich, J. Silterra, J. T. Webber, J. Slavič, J. Nothman, J. Buchner, J. Kulick, J. L. Schönberger, J. V. de Miranda Cardoso, J. Reimer, J. Harrington, J. L. C. Rodríguez, J. Nunez-Iglesias, J. Kuczynski, K. Tritz, M. Thoma, M. Newville, M. Kümmerer, M. Bolingbroke, M. Tartre, M. Pak, N. J. Smith, N. Nowaczyk, N. Shebanov, O. Pavlyk, P. A. Brodtkorb, P. Lee, R. T. McGibbon, R. Feldbauer, S. Lewis, S. Tygier, S. Sievert, S. Vigna, S. Peterson, S. More, T. Pudlik, T. Oshima, T. J. Pingel, T. P. Robitaille, T. Spura, T. R. Jones, T. Cera, T. Leslie, T. Zito, T. Krauss, U. Upadhyay, Y. O. Halchenko, Y. Vázquez-Baeza, SciPy 1.0: fundamental algorithms for scientific computing in Python. Nature Methods 2020 17:3. 17, 261–272 (2020).

93. S. van der Walt, J. L. Schönberger, J. Nunez-Iglesias, F. Boulogne, J. D. Warner, N. Yager, E. Gouillart, T. Yu, Scikit-image: Image processing in python. PeerJ. 2014, e453 (2014).

94. C. C. Chang, C. C. Chow, L. C. A. M. Tellier, S. Vattikuti, S. M. Purcell, J. J. Lee, Second-generation PLINK: rising to the challenge of larger and richer datasets. Gigascience. 4, 7 (2015).

95. P. R. Loh, G. Tucker, B. K. Bulik-Sullivan, B. J. Vilhjálmsson, H. K. Finucane, R. M. Salem, D. I. Chasman, P. M. Ridker, B. M. Neale, B. Berger, N. Patterson, A. L. Price, Efficient Bayesian mixed model analysis increases association power in large cohorts. Nat Genet. 47, 284 (2015).

96. D. M. Church, V. A. Schneider, T. Graves, K. Auger, F. Cunningham, N. Bouk, H. C. Chen, R. Agarwala, W. M. McLaren, G. R. S. Ritchie, D. Albracht, M. Kremitzki, S. Rock, H. Kotkiewicz, C. Kremitzki, A. Wollam, L. Trani, L. Fulton, R. Fulton, L. Matthews, S. Whitehead, W. Chow, J. Torrance, M. Dunn, G. Harden, G. Threadgold, J. Wood, J. Collins, P. Heath, G. Griffiths, S. Pelan, D. Grafham, E. E. Eichler, G. Weinstock, E. R. Mardis, R. K. Wilson, K. Howe, P. Flicek, T. Hubbard, Modernizing reference genome assemblies. PLoS Biol. 9 (2011), doi:10.1371/JOURNAL.PBIO.1001091.

97. J. Yang, S. H. Lee, M. E. Goddard, P. M. Visscher, GCTA: A Tool for Genome-wide Complex Trait Analysis. Am J Hum Genet. 88, 76 (2011).

98. R. Border, G. Athanasiadis, A. Buil, A. Schork, N. Cai, A. Young, T. Werge, J. Flint, K. Kendler, S. Sankararaman, A. Dahl, N. Zaitlen, Cross-trait assortative mating is widespread and inflates genetic correlation estimates. bioRxiv, in press, doi:10.1101/2022.03.21.485215.

99. R. Border, S. O’Rourke, T. de Candia, M. E. Goddard, P. M. Visscher, L. Yengo, M. Jones, M. C. Keller, Assortative mating biases marker-based heritability estimators. Nat Commun. 13, 660–660 (2022).

100. Mathematical contributions to the theory of evolution.—On a form of spurious correlation which may arise when indices are used in the measurement of organs. Proceedings of the Royal Society of London. 60, 489–498 (1897).

101. J. W. Belmont, P. Hardenbol, T. D. Willis, F. Yu, H. Yang, L. Y. Ch’Ang, W. Huang, B. Liu, Y. Shen, P. K. H. Tam, L. C. Tsui, M. M. Y. Waye, J. T. F. Wong, C. Zeng, Q. Zhang, M. S. Chee, L. M. Galver, S. Kruglyak, S. S. Murray, A. R. Oliphant, A. Montpetit, F. Chagnon, V. Ferretti, M. Leboeuf, M. S. Phillips, A. Verner, S. Duan, D. L. Lind, R. D. Miller, J. Rice, N. L. Saccone, P. Taillon-Miller, M. Xiao, A. Sekine, K. Sorimachi, Y. Tanaka, T. Tsunoda, E. Yoshino, D. R. Bentley, S. Hunt, D. Powell, H. Zhang, I. Matsuda, Y. Fukushima, D. R. Macer, E. Suda, C. Rotimi, C. A. Adebamowo, T. Aniagwu, P. A. Marshall, O. Matthew, C. Nkwodimmah, C. D. M. Royal, M. F. Leppert, M. Dixon, F. Cunningham, A. Kanani, G. A. Thorisson, P. E. Chen, D. J. Cutler, C. S. Kashuk, P. Donnelly, J. Marchini, G. A. T. McVean, S. R. Myers, L. R. Cardon, A. Morris, B. S. Weir, J. C. Mullikin, M. Feolo, M. J. Daly, R. Qiu, A. Kent, G. M. Dunston, K. Kato, N. Niikawa, J. Watkin, R. A. Gibbs, E. Sodergren, G. M. Weinstock, R. K. Wilson, L. L. Fulton, J. Rogers, B. W. Birren, H. Han, H. Wang, M. Godbout, J. C. Wallenburg, P. L’Archevêque, G. Bellemare, K. Todani, T. Fujita, S. Tanaka, A. L. Holden, F. S. Collins, L. D. Brooks, J. E. McEwen, M. S. Guyer, E. Jordan, J. L. Peterson, J. Spiegel, L. M. Sung, L. F. Zacharia, K. Kennedy, M. G. Dunn, R. Seabrook, M. Shillito, B. Skene, J. G. Stewart, D. L. Valle, E. W. Clayton, L. B. Jorde, A. Chakravarti, M. K. Cho, T. Duster, M. W. Foster, M. Jasperse, B. M. Knoppers, P. Y. Kwok, J. Licinio, J. C. Long, P. Ossorio, V. O. Wang, C. N. Rotimi, P. Spallone, S. F. Terry, E. S. Lander, E. H. Lai, D. A. Nickerson, G. R. Abecasis, D. Altshuler, M. Boehnke, P. Deloukas, J. A. Douglas, S. B. Gabriel, R. R. Hudson, T. J. Hudson, L. Kruglyak, Y. Nakamura, R. L. Nussbaum, S. F. Schaffner, S. T. Sherry, L. D. Stein, T. Tanaka, The International HapMap Project. Nature 2004 426:6968. 426, 789–796 (2003).

102. N. J. Higham, Computing the nearest correlation matrix - A problem from finance. IMA Journal of Numerical Analysis. 22, 329–343 (2002).

103. J. L. Horn, A rationale and test for the number of factors in factor analysis. Psychometrika 1965 30:2. 30, 179–185 (1965).

104. J. Yang, A. Bakshi, Z. Zhu, G. Hemani, A. A. E. Vinkhuyzen, I. M. Nolte, J. v. van Vliet- Ostaptchouk, H. Snieder, T. Esko, L. Milani, R. Mägi, A. Metspalu, A. Hamsten, P. K. E. Magnusson, N. L. Pedersen, E. Ingelsson, P. M. Visscher, Genome-wide genetic homogeneity between sexes and populations for human height and body mass index. Hum Mol Genet. 24, 7445–7449 (2015).

105. S. Purcell, B. Neale, K. Todd-Brown, L. Thomas, M. A. R. Ferreira, D. Bender, J. Maller, P. Sklar, P. I. W. de Bakker, M. J. Daly, P. C. Sham, PLINK: A Tool Set for Whole-Genome Association and Population-Based Linkage Analyses. Am J Hum Genet. 81, 559 (2007).

106. A. R. Quinlan, I. M. Hall, BEDTools: a flexible suite of utilities for comparing genomic features. Bioinformatics. 26, 841–842 (2010).

107. C. A. de Leeuw, J. M. Mooij, T. Heskes, D. Posthuma, MAGMA: Generalized Gene-Set Analysis of GWAS Data. PLoS Comput Biol. 11, e1004219 (2015).

108. A. A. Gusev, A. Ko, H. Shi, G. Bhatia, W. Chung, B. W. J. H. Penninx, R. Jansen, E. J. C. de Geus, D. I. Boomsma, F. A. Wright, P. F. Sullivan, E. Nikkola, M. Alvarez, M. Civelek, J. Lusis, T. Lehtimäki, E. Raitoharju, M. Kähönen, I. Seppälä, O. T. Raitakari, J. Kuusisto, M. Laakso, A. L. Price, P. Pajukanta, B. Pasaniuc, Integrative approaches for large-scale transcriptome-wide association studies. Nat Genet. 48, 245–252 (2016).

109. N. E. Renthal, P. Nakka, J. M. Baronas, H. M. Kronenberg, J. N. Hirschhorn, Genes with specificity for expression in the round cell layer of the growth plate are enriched in genomewide association study (GWAS) of human height. Journal of Bone and Mineral Research. 36, 2300–2308 (2021).

110. S. Durinck, P. T. Spellman, E. Birney, W. Huber, Mapping identifiers for the integration of genomic datasets with the R/Bioconductor package biomaRt. Nature Protocols 2009 4:8. 4, 1184–1191 (2009).

111. J. C. Lui, Y. H. Jee, P. Garrison, J. R. Iben, S. Yue, M. Ad, Q. Nguyen, B. Kikani, Y. Wakabayashi, J. Baron, Differential aging of growth plate cartilage underlies differences in bone length and thus helps determine skeletal proportions. PLoS Biol. 16 (2018), doi:10.1371/JOURNAL.PBIO.2005263.

112. C. Palazzo, C. Nguyen, M. M. Lefevre-Colau, F. Rannou, S. Poiraudeau, Risk factors and burden of osteoarthritis. Ann Phys Rehabil Med. 59, 134–138 (2016).

113. L. A. C. Millard, N. M. Davies, T. R. Gaunt, G. D. Smith, K. Tilling, Software Application Profile: PHESANT: a tool for performing automated phenome scans in UK Biobank. Int J Epidemiol. 47, 29–35 (2018).

114. B. Neale, Neale Lab UK_Biobank_GWAS. Github (2021).

115. B. Bulik-Sullivan, H. K. Finucane, V. Anttila, A. Gusev, F. R. Day, P. R. Loh, L. Duncan, J. R. B. Perry, N. Patterson, E. B. Robinson, M. J. Daly, A. L. Price, B. M. Neale, An atlas of genetic correlations across human diseases and traits. Nature Genetics 2015 47:11. 47, 1236–1241 (2015).

116. C. P. Bird, B. E. Stranger, M. Liu, D. J. Thomas, C. E. Ingle, C. Beazley, W. Miller, M. E. Hurles, E. T. Dermitzakis, Fast-evolving noncoding sequences in the human genome. Genome Biol. 8 (2007), doi:10.1186/GB-2007-8-6-R118.

117. E. C. Bush, B. T. Lahn, A genome-wide screen for noncoding elements important in primate evolution. BMC Evol Biol. 8 (2008), doi:10.1186/1471-2148-8-17.

118. R. M. Gittelman, E. Hun, F. Ay, J. Madeoy, L. Pennacchio, W. S. Noble, R. D. Hawkins, J. M. Akey, Comprehensive identification and analysis of human accelerated regulatory DNA. Genome Res. 25, 1245–1255 (2015).

119. K. S. Pollard, S. R. Salama, B. King, A. D. Kern, T. Dreszer, S. Katzman, A. Siepel, J. S. Pedersen, G. Bejerano, R. Baertsch, K. R. Rosenbloom, J. Kent, D. Haussler, Forces shaping the fastest evolving regions in the human genome. PLoS Genet. 2, 1599–1611 (2006).

120. S. Prabhakar, J. P. Noonan, S. Pääbo, E. M. Rubin, Accelerated evolution of conserved noncoding sequences in humans. Science. 314, 786 (2006).

121. W. McLaren, L. Gil, S. E. Hunt, H. S. Riat, G. R. S. Ritchie, A. Thormann, P. Flicek, F. Cunningham, The Ensembl Variant Effect Predictor. Genome Biol. 17, 1–14 (2016).

122. M. L. Benton, A. Abraham, A. L. LaBella, P. Abbot, A. Rokas, J. A. Capra, The influence of evolutionary history on human health and disease. Nature Reviews Genetics 2021 22:5. 22, 269–283 (2021).

123. B. Vernot, S. Tucci, J. Kelso, J. G. Schraiber, A. B. Wolf, R. M. Gittelman, M. Dannemann, S. Grote, R. C. McCoy, H. Norton, L. B. Scheinfeldt, D. A. Merriwether, G. Koki, J. S. Friedlaender, J. Wakefield, S. Pääbo, J. M. Akey, Excavating Neandertal and Denisovan DNA from the genomes of Melanesian individuals. Science. 352, 235–239 (2016).

124. K. C. Tashman, R. Cui, L. J. O’Connor, B. M. Neale, H. K. Finucane, Significance testing for small annotations in stratified LD-Score regression. medRxiv, in press, doi:10.1101/2021.03.13.21249938.

